# AI-based Prediction of Protein Corona Composition on DNA Nanostructures

**DOI:** 10.1101/2024.08.25.609594

**Authors:** Jared Huzar, Roxana Coreas, Markita P. Landry, Grigory Tikhomirov

## Abstract

DNA nanotechnology has emerged as a powerful approach to engineering biophysical tools, therapeutics, and diagnostics because it enables the construction of designer nanoscale structures with high programmability. Based on DNA base pairing rules, nanostructure size, shape, surface functionality, and structural reconfiguration can be programmed with a degree of spatial, temporal, and energetic precision that is difficult to achieve with other methods. However, the properties and structure of DNA constructs are greatly altered *in vivo* due to spontaneous protein adsorption from biofluids. These adsorbed proteins, referred to as the protein corona, remain challenging to control or predict, and subsequently, their functionality and fate *in vivo* are difficult to engineer. To address these challenges, we prepared a library of diverse DNA nanostructures and investigated the relationship between their design features and the composition of their protein corona. We identified protein characteristics important for their adsorption to DNA nanostructures and developed a machine-learning model that predicts which proteins will be enriched on a DNA nanostructure based on the DNA structures’ design features and protein properties. Our work will help to understand and program the function of DNA nanostructures *in vivo* for biophysical and biomedical applications.

## Introduction

Simple base pairing rules (A-T & G-C) have enabled engineering very complex nanometer-precise DNA structures. Nanostructures consisting of 10,000 different DNA strands and reaching gigadaltons size with programmable behavior have been constructed^1,2,3,4^. These nanostructures can be designed to have precise arrangements of targeting ligands and dynamic reconfigurations with programmable kinetics^5–7^. DNA nanotechnology is now emerging as a versatile toolkit to study and alter biological processes^8,9^. As sensors, DNA constructs have been developed that can sense piconewton scale forces^9^, changes in temperature^10^, acidity^11^, and analyte presence^12^. As medicines, DNA nanostructures can sequester and on-demand release cargos^13,14^ including functional nucleic acids (DNAzymes^15^, siRNA^16^, gene-encoding DNAs^17^, etc.) and proteins^18^. By modulating the size, shape, or addition of chemical moieties to nanostructures, DNA nanostructures can be programmed to reconfigure^14,19^, to target specific tissues^20–22^, and to be preferentially internalized by certain cell types^23,24^. However, all this powerful programmability of DNA nanotechnology is impacted by protein adsorption when nanostructures are immersed in biological environments. Analogous to protein adsorption that occurs on other materials, protein corona changes DNA nanostructures’ capabilities *in vivo*.

When DNA nanostructures are introduced to biological fluids (plasma, serum, etc.), they are spontaneously covered by a multitude of biomolecules forming the biomolecular corona^20,25^. The proteins that comprise the biomolecular corona drastically change how nanomaterials interact with biosystems^26^ by altering nanoparticle size^27^, shape^28^, and physicochemical properties^29^. As the outermost entity, the biomolecular corona provides nanomaterials with a surrogate biological identity that can have unintended effects on nanostructure uptake, biodistribution, and immunogenicity^26,30,31^. Despite the biomolecular corona being a well-documented phenomenon across nanobiotechnology, the factors underlying biomolecular corona formation, especially for DNA nanostructures, remain insufficiently understood, limiting the applications of DNA nanotechnologies in biological fluids. Machine learning models have recently been used to elucidate the factors governing protein absorption on inorganic nanoparticles, liposomes, and carbon nanotubes^32–34^. Machine learning models have also been implemented to predict binding interactions between proteins and short nucleic acids^35^, however, *in silico* methods that accurately predict the interactions between biomolecules and DNA nanostructures, both *in vitro* and *in vivo* have yet to be developed and require extensive datasets built upon fundamental research. We sought to develop an interpretable machine learning classifier that can accurately predict which proteins will be found in the biomolecular coronas of DNA nanostructures.

To this end, we designed, synthesized, and characterized an array of DNA nanostructures with diverse design features including sizes, shapes, charges, and surface modifications including aptamers and cholesterol. We also synthesized several DNA nanostructures coated with oligolysine, a common polycationic polymer used to enhance cellular uptake and stability of DNA in biological environments^36^. With this library of nanostructures, we quantitatively measured the abundance of proteins adsorbed to the DNA nanostructures in human serum using gel electrophoresis shift assays and ultra-high performance liquid chromatography tandem mass spectrometry (UHPLC-MS/MS). We observed differences along all DNA nanostructure design feature axes, i.e. certain proteins preferably adsorbed onto certain nanostructures.

With this rich dataset, we developed an explainable machine learning model that can, based on basic DNA nanostructure and protein features, predict with 92% accuracy whether a protein will be present in the biomolecular corona. Thus, the protein corona can be predicted and even engineered with well-established DNA nanostructure design approaches. We leveraged this model to quantitatively probe relationships between size, shape, surface charge, and other features of DNA nanostructures and corona proteins. Thus, we gained insights into the factors governing protein adsorption and the biological pathways likely influenced by the adsorbed proteins. These findings will help guide the design of DNA nanostructures for biophysical and biomedical applications that are subject to biomolecular corona formation.

## Results

### Synthesis of DNA Nanostructures and Design Characterization

We synthesized a compact yet diverse library of DNA nanostructures (**Fig. 1a**) to elucidate the effect of various design features on the protein corona composition. We tested the effect of (1) size, (2) shape, (3) surface functionalization, and (4) surface charge (**Fig. S1**). These features are known to influence cellular uptake and other biological functions of DNA nanostructures^23^. DNA nanostructures are often functionalized with the lipophilic molecule cholesterol to facilitate insertion into lipid membranes^37^, and aptamers for the targeting of cell-surface receptors^38^; thus we synthesized several structures with either aptamers or cholesterol. Cholesterol-modified DNA nanostructures are also known to seed a larger protein corona than its unmodified DNA-only counterpart^20^. Finally, we modified the surface charge by coating our DNA nanostructures with a polylysine (**Fig. 1b**), as it was reported to enhance stability and cellular uptake^36^. In addition, polylysine-PEG polymer coating affects protein corona formation by stabilizing the integrity of DNA nanostructures in complex milieus^25^. Surface charge is affected by the cationic polymer coating, so we measured many structures’ zeta (ζ) potential using an electrophoretic light scattering zetasizer (**Table S1**).

**Figure 1.**
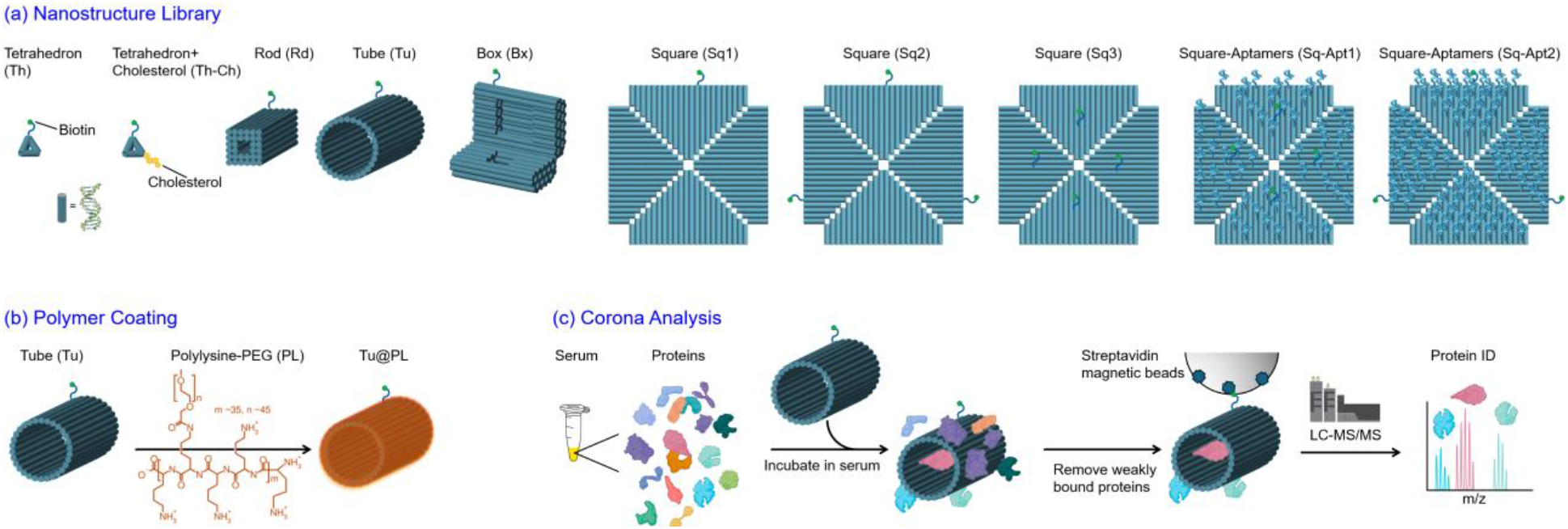
DNA nanostructures and protein corona analysis approach. **(a)** DNA nanostructures used in this study. **(b)** Schematic of nanostructures coating with PLL-g-PEG5K cationic polymer. **(c)** Schematic of protein corona analysis by magnetic bead separation and liquid chromatography tandem mass-spectrometry.

Based on these prior findings, we synthesized 17 DNA nanostructures (**Fig. 1**): (1) a DNA tetrahedron with 20mer duplexes per edge (Th)^20^, (2) the same tetrahedron functionalized with a single cholesterol (Th-Ch), (3) a slightly modified origami box (Bx)^39^, (4) square origami tiles (Sq)^4^ with differing numbers and positions of aptamers and biotins, (5) a hollow origami tube (Tu)^1^, and (6) a 32 helix origami rod (Rd)^40^; many of these structures were also synthesized with a polycationic PLL-PEG coating (@PL) (e.g., tube in **Fig. 1b**). Overall, these structures surveyed a wide design space over the various biologically relevant parameters (**Fig. 1a and S1**).

### Experimental Determination of Protein Corona on DNA nanostructures

We synthesized and purified DNA nanostructures containing a ssDNA oligonucleotide modified with a terminal biotin for attachment to streptavidin-coated magnetic beads (**Fig. 1**). Initially, we incubated the nanostructures at a final concentration of 0.5 μM in 20-fold diluted pooled human serum (protein concentration ∼3.5 mg/mL) and isolated the nanostructures along with their adsorbed proteins from unbound, serum proteins via a pull-down assay (**Fig. 1c**). We performed gel electrophoretic analysis and observed that the proteins bound to DNA nanostructures were distinct from those bound to the magnetic bead in serum, our negative control (**Fig. S2a**). In addition, we observed a substantial difference in corona composition between the cholesterol-modified and non-cholesterol-modified tetrahedron (**Fig. S2b**), corroborating previously reported results^20^. Having thus validated this protocol, we expanded this corona extraction protocol to test several nanostructures at lower, biologically relevant concentrations^41^. For higher-throughput and quantitative determination of corona composition, we performed SDS-PAGE separation followed by UHPLC-MS/MS.

### UHPLC-MS/MS Analysis of DNA Nanostructures’ Protein Corona

Studies have shown that dose-dependent effects can be observed with DNA nanostructures; biologically relevant doses (10 pM – 100 nM) can modulate cell inflammatory responses^41,42^. To characterize the protein corona adsorbed onto DNA nanostructures, we incubated the 17 unique DNA structures at a final concentration of 50 pM in pooled human sera with a final protein concentration of 5mg/mL and performed UHPLC-MS/MS to identify and quantify the relative abundances of adsorbed proteins. Across all DNA nanostructure coronas, we identified 575 proteins that showed differential abundance in the corona when compared to their prevalence in serum. Enriched corona proteins, (log_2_(fold change) >0), were identified as proteins that were more abundant in the nanostructure corona than in serum. Conversely, proteins that were more abundant in the controls, (log_2_(fold change) <0), were highly abundant in sera, and suggest that they may possess minimal to no affinity to the nanostructures. The magnitude of change in protein prevalence for each nanostructure relative to the controls is shown in **Fig. S3**; volcano plots for each nanostructure show the statistical significance between relative abundances (-log_10_(*p-*value)) versus the magnitude of change (log_2_(fold change)) of protein relative abundances (**Fig. S4**). Within our dataset, there were unique proteins that were present on the nanostructures but were not identified in serum nor on the magnetic bead, likely due to their low abundance in serum (**Fig. S5**). Of these uniquely present proteins, several were involved in binding: Bx-PL enriched proteins involved in purine nucleotide and guanosine diphosphate binding; Sq-Apt1@PL and Sq3@PL enriched proteins associated in membrane adhesion, intracellular transport, and vesicle-mediated transport; Th-Ch@PL, Sq1, Rd, Sq3@PL, Sq-Apt1@PL and Tu enriched proteins involved with membrane docking; Th@PL enriched proteins that interact with biomolecules within the extracellular space and/or exosomes. Moreover, proteins associated with the positive regulation of early endosome to late endosome transport were enriched on Sq-Apt1, Sq-Apt1@PL, Sq3@PL, Sq-Apt2@PL, Rd, Th-Ch@PL, Sq1, and Tu. Several nanostructures enriched unique proteins associated with the positive regulation and/or activation of immunological processes; most notably, Sq3@PL enriched proteins involved in antimicrobial humoral responses, fibrinolysis, integrin activation, and regulation of ERK1 and ERK2 cascades; Sq-Apt1@PL enriched autocrine signaling proteins; molecular chaperones that fold stress-denatured proteins were enriched on Sq-Apt1, Sq-Apt2@PL, and Th; proteins associated with positive regulation of establishment of T cell polarity were enriched on Sq2; and proteins involved in wound responses and healing were enriched on Th-Ch@PL. Contrastingly, proteins involved in the negative regulation of immunological processes, including complement activation and regulation of extrinsic apoptotic signaling via death domain receptors, were enriched on Sq-Apt1@PL and Sq3@PL.

The magnetic beads (MB) used to collect the DNA nanostructure-protein corona complexes (**Fig. 1c**) also adsorbed serum proteins even when no nanostructures were present (**Fig. S2**). Thus, we compared the composition of differentially expressed proteins adsorbed on the MB relative to serum, with the collective composition of corona proteins adsorbed on all 17 of the nanostructures. We found that all differentially expressed proteins on the MB were adsorbed across the different nanostructures’ coronas, many of which were more abundant on the complex formed between the DNA nanostructures and the MB than on the MB itself. To factor out the influence of the proteins adsorbed directly onto the magnetic beads, the spectral counts for the proteins adsorbed on the nanostructures were subtracted by the average spectral count identified on the magnetic beads. The magnitude of change in protein prevalence for each nanostructure relative to serum, extracting the enrichment on the magnetic beads, is shown in **Fig. S6**. Recognizing these distinct protein preferences, we aimed to get deeper insights into these interactions. To do so we performed two different binary classifications of the individual proteins for each nanostructure, using the dataset in which the magnetic bead enrichment was subtracted. Proteins that composed the nanostructure corona included those that were calculated as enriched or depleted, as well as those that were uniquely present on the corona but not in the sera controls. These thresholds were decided as we expect all proteins present in the biomolecular corona will affect the nanostructures’ physiochemical and pharmacokinetic properties, making it important to predict and understand the entirety of the corona’s composition. However, we anticipate exploring enriched proteins will produce a model that is independent of protein concentration in the serum, making it more generalizable and even combinable with data from other biological fluids. Herein, we use the term ‘enriched’ to describe proteins found in higher levels in the corona than in serum, ‘depleted’ to represent proteins found in lower abundances on nanostructures relative to serum, and ‘present’ to describe all proteins found in the corona irrespective of its abundance relative to the serum levels.

### Differences and Similarities Across Origami

Using our library of nanostructures, we probed the effects of nanostructure design features and protein properties on the composition of the corona. We developed a database of functional, structural, and physicochemical properties of proteins to identify the protein features important in determining a protein’s abundance in the corona. Data was obtained by scraping UniProt^43^, with the Quantiprot Python package^44^, and NetSurf 2.0^45^ (see the methods section for more information on the specific metrics). Overall, our database leveraged single amino acid level properties, secondary structure information, and functional information to encompass protein properties most likely to affect protein corona adsorption. We next quantified variation in protein corona composition between different DNA nanostructures. Considering the total corona composition of each nanostructure, we observed all nanostructures differentiate themselves by their unique corona compositions relative to base serum composition (**Fig. 2a**). Interestingly, we find that corona content is not a simple reflection of each protein’s relative abundance in the serum, rather that each nanostructure corona has certain proteins that are enriched or depleted relative to their presence in human serum. To analyze this trend further, we considered the clustering of nanostructures based on their corona compositions. This revealed two distinct groups divided by the presence or absence of the polymer coating on the nanostructure. Among nanostructures with a polymer coating, there was a high degree, ∼75%, of protein compositional similarity (**Fig. 2b**). This homogeneity is even more pronounced among nanoparticles without a polymer coating (**Fig. 2b**). Despite this polymer-driven clustering, 117 proteins – relative to 534 total analyzed proteins – were universally adsorbed across all nanostructures. These 114 universally adsorbed proteins exhibited a significant level of connectivity, referring to similarity regarding their endogenous biological roles (**Fig. 2c**). Of the 117 proteins, several functional clusters emerged. Most predominantly, histone proteins represented a large fraction of these proteins, many of which are involved in the formation of nucleosomes, suggesting their propensity to interact with DNA^46^. In addition, other clusters were present consisting primarily of ribosomal proteins, tubulin proteins, and apolipoproteins among others. Finally, several immunoglobulin proteins were universally present. Immunoglobin proteins have been associated with the opsonization of nanoparticles^47^, and thus these results support the notion that the presence of foreign DNA in the form of nanostructures is likely to trigger their clearance. Notably, this opsonization phenomenon is likely to occur regardless of nanostructure size and shape. Both apolipoproteins and immunoglobins are commonly present in the coronas of other organic nanoparticles, like liposomes^48,49^, and inorganic nanoparticles^50^.

**Figure 2.**
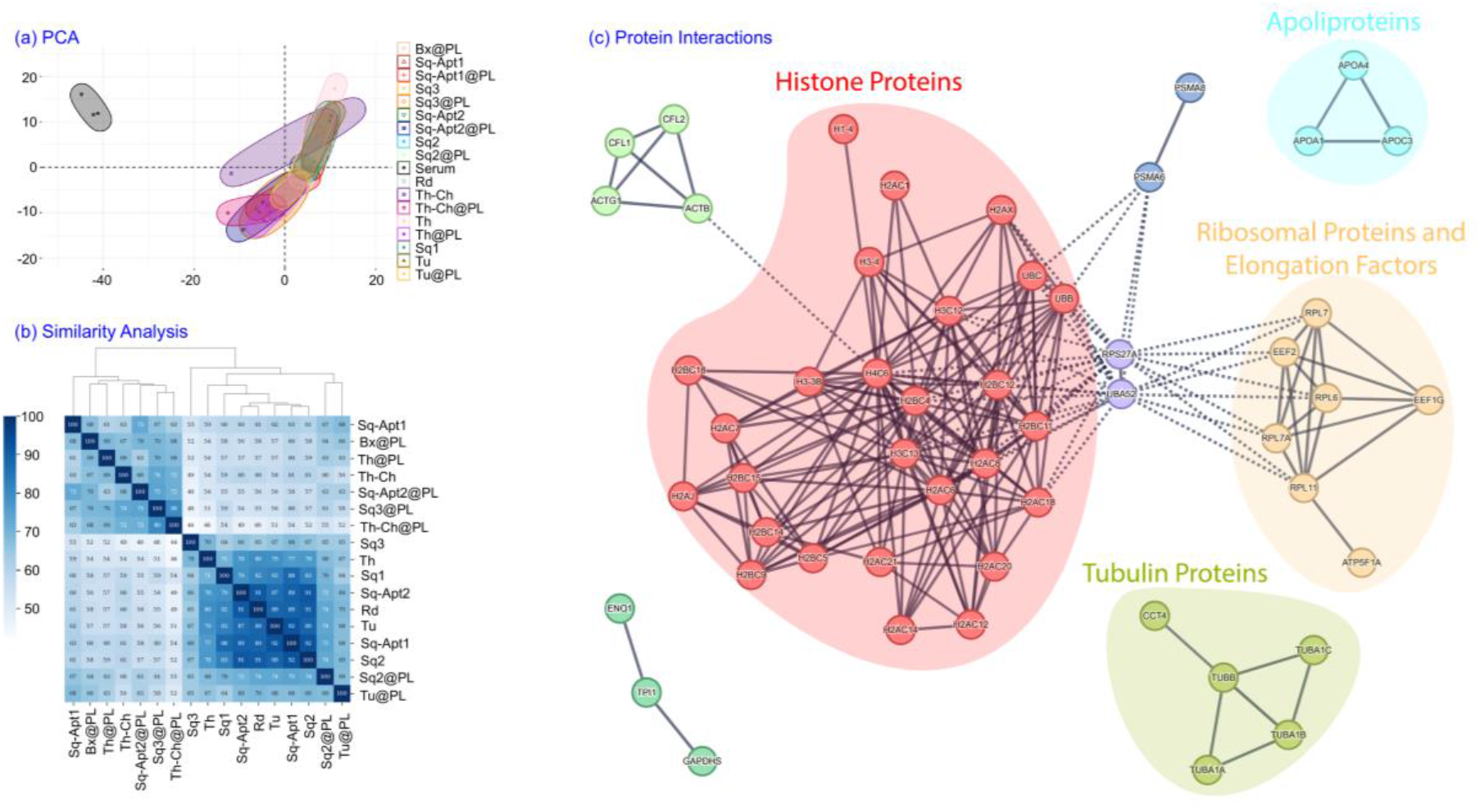
Protein corona analysis for polymer coated and uncoated DNA nanostructures. **(a)** PCA of the protein coronas of all nanostructures and serum, with magnetic bead subtracted. PC1 and PC2 accounted for 37.3% (25.6% and 11.7%, respectively) of the total variance in the data. **(b)** Similarity heatmap plot with hierarchical clustering of the protein corona compositions for all nanostructures. **(c)** Interaction networks between proteins that are universally present across all nanostructures. Proteins with no interactions were removed.

Notably, thrombospondin-1 (TSP1), which interacts with several cell adhesion receptors^51^, was preferentially absorbed on all nanostructures with the @PL coating, with the highest enrichment observed on Th@PL. Nanostructures coated with @PL also enriched plasma serine and protein z-dependent protease inhibitors; the highest enrichment of these modulatory serpins that regulate inflammatory responses^52^ was observed on Box@PL. Other immunomodulatory proteins that were preferentially adsorbed onto the @PL coated nanostructures include carboxypeptidase B2^53^, most abundant on Th@PL, alpha-2-HS-glycoprotein (fetuin-A) and its less abundant homolog fetuin-B^54^, most abundant on Th@PL and Sq3@PL, respectively, phospholipid transfer protein^55^, most abundant on Sq-Apt1@PL, lipopolysaccharide-binding protein^56^, most abundant on Sq3@PL, CD5 antigen-like (CD5L)^57^, most abundant on Bx@PL, complement components C6, C9, and factor B^58^, most abundant on Th@PL, Sq2@PL, and Tu@PL, respectively. Contrastingly, ELAV-like protein 1, which suppresses inflammatory responses^59^, was preferentially adsorbed on the non-coated nanostructures, with the highest enrichment observed on Tu. Inter-alpha-trypsin inhibitor heavy chain 2 (ITIH2), which inhibits complement activation^60^, was also preferentially adsorbed by non-coated nanostructures, with the highest enrichment observed in Sq1. However, several immunogenic proteins were also preferentially absorbed on the non-coated nanostructures and not their @PL coated counterparts, including elongation factor 2^61^, which was most abundant on Rd, vitronectin^62^, most abundant on Sq1, and complement C1q subcomponent subunit C^58^, most abundant on Sq3. Additionally, histone proteins H1, H2, and H3, involved in packaging DNA into chromatin and transcriptional activation^63^, were preferentially enriched on nanostructures lacking the @PL coating; Sq3 had the highest enrichment of histone proteins. Apolipoproteins A-I and B-100, the former having immunogenic properties and the latter potentially exhibiting such properties^64,65^, were enriched on several nanostructures, irrespective of @PL coating presence, yet Tu@PL and Th-Ch@PL had the highest enrichments, respectively.

### Effect of protein composition

We anticipated the 117 universally present corona proteins would contain common structural features that influence their adsorption to nanostructures irrespective of nanostructure polymer coating presence or absence. Indeed, when compared with proteins that were not found in the corona of any nanostructures, clear trends emerge differentiating the universally adsorbed and universally absent proteins. A set of 27 protein physiochemical properties exhibited statistically significant distributions (p-value < 0.01). Highlighting a subset of these 27, several interesting patterns emerged. We observed that proteins with greater flexibility and a more positive charge exhibited an increased propensity for adsorption to the nanostructures (**Fig. 3a**), which makes sense considering that DNA nanostructures are negatively charged and relatively rigid. A similar phenomenon of flexible proteins being enriched in a nanoparticle corona was observed with single-walled carbon nanotubes (SWCNTs)^33^, a rigid, inorganic nanoparticle. Interestingly, larger proteins were less likely to be adsorbed (**Fig. 3b)**. It has been previously reported that the adsorption of small proteins onto nanostructures is an enthalpy-driven process, versus larger proteins that adsorb via an entropy-driven mechanism^47^. As such, we expect this protein size-dependent adsorption observation may be temperature dependent, as well as nanostructure dependent. For this reason, we performed all corona adsorption experiments at 37°C to best mimic conditions experienced *in vivo*. We found that proteins’ different amino acid compositions elicited different effects on the protein’s adsorption to nanostructures. For example, proteins with larger fractions of amino acids with hydrophobic side chains (phenylalanine, leucine, and tryptophan) were less likely to be adsorbed to hydrophilic DNA nanostructures (**Fig. 3c**). In addition, positively charged lysine residues were associated with greater protein adsorption (**Fig. 3d**).

**Figure 3.**
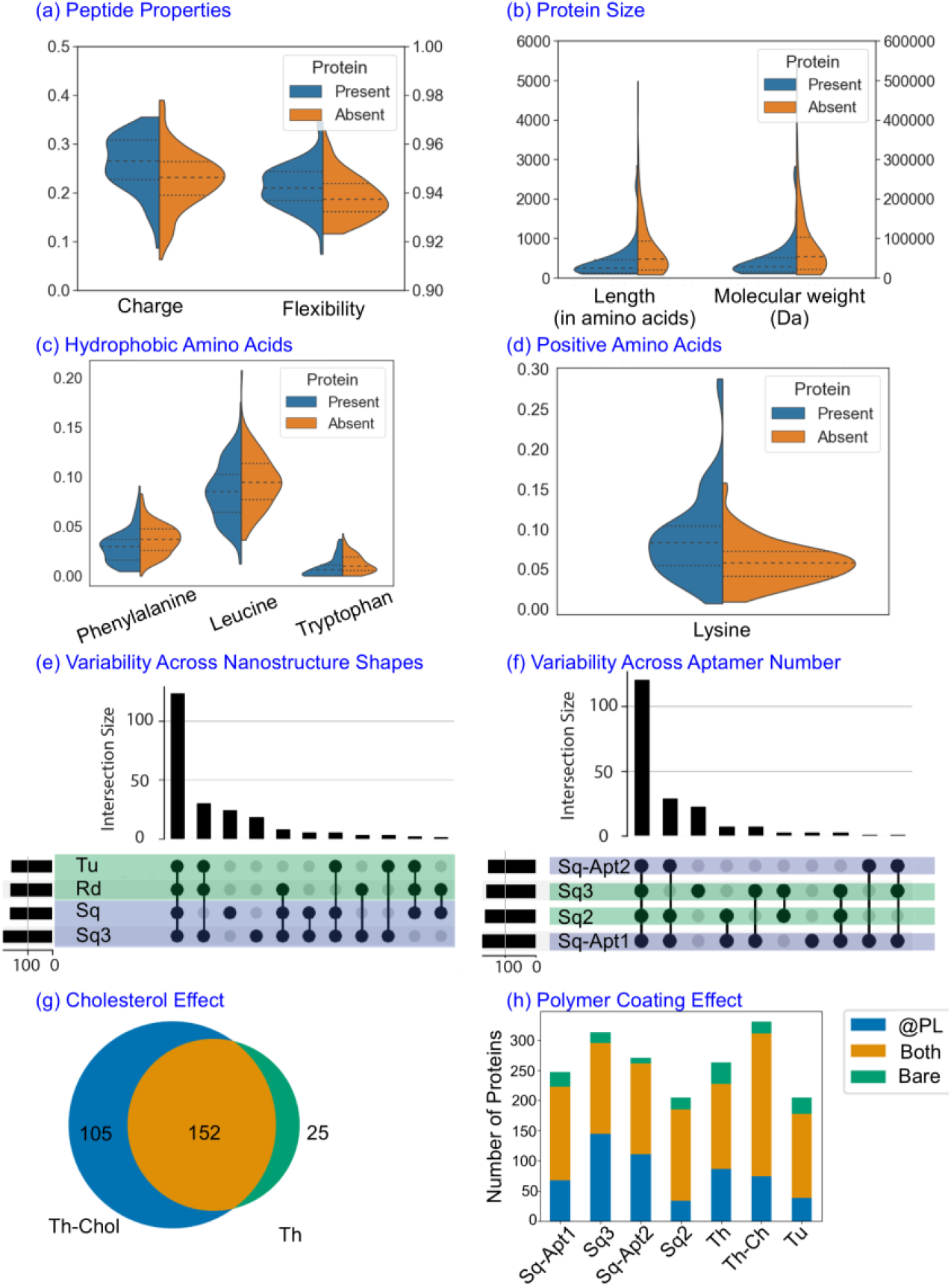
Effect of protein and DNA nanostructure composition on corona content. **(a-d)** Subgroups of protein properties significantly different between universally present and universally absent proteins from the corona. **(a)** Peptide properties. **(b)** Peptide size. **(c)** Hydrophobic amino acids. **(d)** Amino acids with positively charged side chains. **(e)** Effect of the nanostructure shape: number of proteins belonging to one, some, or all the Rd, Tu, Sq-Apt1, and Sq nanostructures’ coronas. Blue samples are 2D square sheets, green samples are elongated more rigid 3D shapes. **(f)** Effect of the number and position of aptamers or biotin tags on the same square origami: number of proteins belonging to one, some, or all the Sq-Apt2, Sq3, Sq2, Sq-Apt1 nanostructures’ coronas. Blue samples are aptamer functionalized, and green samples are bare. **(g)** Effect of Cholesterol attachment: number of proteins in the corona of tetrahedron with cholesterol only, tetrahedron only, and both. **(h)** Effect of polymer coating: number of proteins present in the coronas of coated structures, their bare counterparts, and both.

### Effect of nanostructure shape and aptamer attachments

We next investigated the influence of nanostructure shape on the protein corona composition. Comparing Sq3 and Sq1, to Rd and Tu, we can consider the effect of aspect ratio and dimensionality (2D versus 3D) while keeping size and physiochemical properties constant. Overall, we observed that many proteins are commonly found in all four structures, further demonstrating the marginal effect of DNA origami structure on protein corona composition (**Fig. 3e**). Many proteins are commonly found in structures even with dissimilar shapes. Of the proteins found in the corona of any of the four structures, 55% are commonly found in all 4 nanostructures’ coronas. This represents a significant departure from trends observed between cholesterol-modified/unmodified nanostructures, where the presence of cholesterol greatly increased the diversity of proteins in the corona. Consequently, we conclude DNA nanostructure shape has a limited role in protein adsorption for nanostructure shapes tested herein. Next, given the utility of aptamers in biological applications involving DNA nanostructures, we investigated the effect aptamer number and positioning had on corona formation on an otherwise identical DNA nanostructure. We chose to functionalize our nanostructures with the CoV2-RBD-1C aptamer, discovered by Song et al^66^, an aptamer targeting the receptor-binding domain of the SARS-CoV-2 spike glycoprotein. We implemented this aptamer because we chose to explore whether aptamers would have any non-specific effect on protein adsorption. To accomplish that, we needed an aptamer targeting a protein that would not be in the pooled human serum, and one that had undergone counter-selection, both criteria that the CoV2-RBD-1C aptamer meets. Sq-Apt2 is the aptamer functionalized equivalent of Sq2, and similarly, Sq-Apt1 is the aptamer functionalized version of Sq3. By comparing Sq-Apt2 to Sq2 and Sq-Apt1 to Sq3, we tested the effect of the aptamers on protein corona formation, and we also tested the effect of their placement and multivalency by comparing Sq-Apt1 to Sq-Apt2. Sq-Apt1 has all aptamers on a single face, while Sq-Apt2 has all aptamers on both faces of the square DNA nanostructure. We decided to explore the effects of multivalency and positioning on corona formation due to their well-documented influence in various biological applications, such as cell labeling^67^. In addition, we hypothesized a greater number of aptamers and more complete decoration (Sq-Apt2), could result in a reduction in protein adsorption due to steric hindrance, an effect previously observed with other polymers^68^. Of proteins found in the corona of any of the four structures, most - 63% - are commonly found across all four structures, and no clear distinctions appear among the aptamer-modified structures to differentiate themselves from their non-functionalized counterparts (**Fig. 3f**). These results emphasize the limited effect of aptamer number or position on the protein corona.

We next investigated the effects of cholesterol modification as well as the @PL polymer coating, since non-DNA modifications are known to significantly affect corona composition. Comparing the tetrahedron with and without cholesterol, we observed that a cholesterol modification causes an almost entirely unique protein corona in addition to the typical corona of the DNA-only nanostructure. The Th-Ch nanostructure corona had 105 proteins found exclusively in the structure, in addition to 152 proteins found in both the Th-Ch corona and Th corona. While Th only had 25 unique proteins in its corona (**Fig. 3g**). In a similar manner, by comparing seven uncoated nanostructures to their polymer coated counterparts, we were able to study the effect of the coating and change in surface charge independently of other variables (**Fig. 3h**). Across the seven nanostructures, each of which was coated with the @PL polymer, most (up to 74%) of proteins adsorbed were present in both the polymer-coated nanostructure and uncoated nanostructure coronas, although several were also uniquely present to either the bare or polymer coated nanostructures’ coronas.

### Developing a Machine Learning Model to Understand and Predict Nanostructure Protein Corona Formation

To better understand the factors influencing the differences in corona formation on DNA nanostructures, we sought to develop an explainable machine learning model. The ability to predict which proteins will be present in a given nanostructure’s corona can help engineer nanostructures with better performance in biofluids and *in vivo*. We chose to implement the XGBoost^69^ algorithm as it is an implementation of gradient-boosting decision trees that maintains the interpretability of decision trees while offering advantages in speed and accuracy. Also, XGBoost has previously been demonstrated as an effective algorithm for protein corona classification^70^. To evaluate the validity of this approach, we tested two other ensemble methods that use sklearn^71^: (1) a Random Forest classifier^72^ and (2) a Gradient boosting classifier^73^. Models were all evaluated using a 10-fold cross-validation with a 90/10 train-test split on the datasets described above. Their performance in identifying both enriched and present proteins was evaluated using the area under the receiver operating curve (AUC), accuracy, f1, precision, and recall. Analyses revealed that XGBoost was the superior model across all metrics (**Fig. S7**). We subsequently chose to use this architecture for the remainder of the analyses. With this approach, we then classified the proteins as enriched/depleted, or present/absent in the corona for all the samples previously described (**Fig. 4a**). We elected to study two classification tasks: the prediction of enriched proteins and the prediction of all proteins present within a DNA nanostructure protein corona. The two classification tasks both have their unique advantages. We expect the prediction of enriched proteins will create a more generalizable model with less of an effect from initial protein concentration in the serum. However, prediction of all proteins present is essential to understanding and engineering nanoparticle fate.

**Figure 4.**
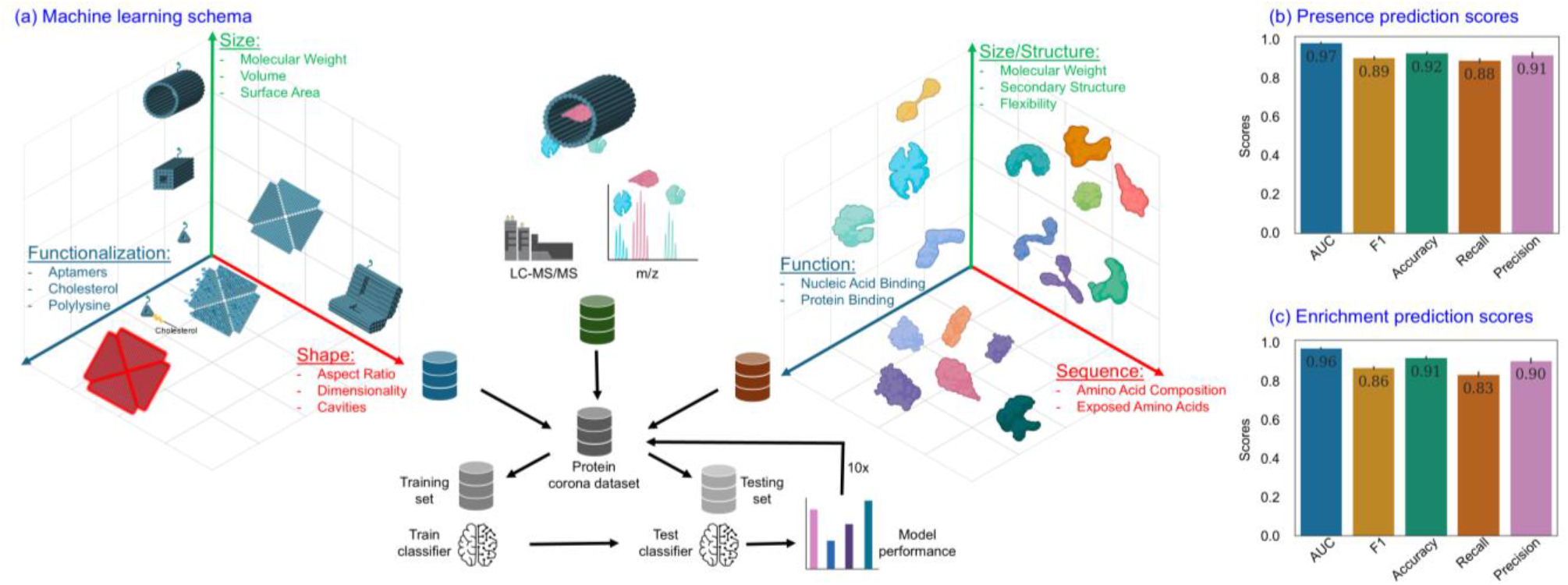
Machine learning model scheme and performance. **(a)** Schema for machine learning model and evaluation protocol. **(b)** Performance metrics of the XGBoost model classifying proteins as present or absent in the nanostructures’ coronas across 10 splits. **(c)** Performance metrics of the XGBoost model classifying proteins as enriched or depleted in the nanostructures’ coronas.

We explored two different ways to train and implement our model. Firstly, we combined the UHPLC-MS/MS datasets for each nanostructure all into one database, and combined protein data with nanostructure physicochemical properties (**Fig. 4a)**. Aggregating the data from each sample is advantageous in several ways. First, as a predictive tool, this architecture now makes the results generalizable to any given DNA nanostructure. A researcher can input the properties of their nanostructure, select a protein of interest, and receive a classification of whether this protein is likely to be enriched and/or abundant in the corona. Secondly, we hypothesized the increase in data accumulated by aggregating the 17 nanostructures’ UHPLC-MS/MS data together would improve the predictive power of the model. Thirdly, using the importance of each feature, this model can be informative about which nanostructure design characteristics can most influentially bias protein adsorption. And finally, we can use this model to identify characteristics of adsorbed proteins that are generalizable across nanostructures.

We then also experimented with training a unique model for individual nanostructures. This approach is advantageous for several reasons. Firstly, by comparing the important features across the different models, we can see which protein properties influence binding most for nanostructures with different properties. Secondly, we expect this model may have improved efficacy as the model does not have to learn the nanostructure features. While it then becomes a simpler classification task, the drawback is there is significantly less data to train on as compared to the bulk dataset.

### Model Performance on Predicting Nanostructure Protein Corona Composition

Implementing our XGBoost model on the aggregated dataset, we achieved 0.97 AUC and 92% accuracy in classifying a protein as present or absent from the protein corona. This demonstrates its high performance in identifying proteins likely to be adsorbed (**Fig. 4b**). Thus, we validated our model as a useful tool allowing for the prediction of proteins found within the protein corona on nanostructures. Utilizing these findings, researchers can now preemptively predict, and account for, the proteins likely to be found on their DNA nanostructures, a first-of-its-kind tool to our knowledge.

Using the same architecture, we classified proteins that are enriched (i.e. more abundant in the corona than in serum) on nanostructures. For this novel classification task, we found an XGBoost model was able to maintain a high level of performance, achieving an AUC of 0.96 and an accuracy of 91% (**Fig. 4c**). Since this model effectively subtracts the serum concentrations of proteins, we expect insights derived from this explainable model can be generalizable to protein adsorption onto DNA nanostructures from other biological fluids. Thus, we successfully implemented and characterized two models with high power for predicting proteins likely to impact the nanostructures’ physiological and pharmacokinetic properties. The models’ high performances also validate them as tools from which we can gain insights into the factors governing protein adsorption on DNA nanostructures.

### Important Protein and Nanostructure Features Governing Protein Adsorption

We next considered which protein features were most informative and predictive to the formation of the nanostructure protein corona. We tested over 600 features quantifying protein size, structure, amino acid composition, and functionality, and DNA nanostructure size, shape, and functionalization. With both models, we found that more than 150 out of over 600 tested features contributed to the classification of proteins as being in versus out of the protein corona (**Table S2**). This result underscores the vast complexity of factors governing protein corona adsorption. To elucidate on a broader scale the properties identified as drivers of protein corona formation, we subset our data into 3 classes: (1) Protein sequence and structure features, (2) Protein functional features, and (3) DNA nanostructure design features. Training the data on each of these subsets alone saw reduced performance as compared to the original dataset containing all this information. Protein sequence and structural features were the most effective at predicting protein presence in the corona, followed by protein function, and lastly origami design (**Fig. 5a**). Specifically, protein sequence and structural features alone effectively predicted protein adsorption with 86% accuracy, protein function predicted protein adsorption with 81% accuracy, and origami design predicted protein adsorption with 70% accuracy. Since the protein features alone were not as effective as when combined with origami design features, we conclude that protein features governing corona adsorption are not generalizable across all nanostructures. However, the moderate predictive power of protein structure alone suggests that protein structure plays a large role in determining protein in-corona presence. DNA nanostructure design features were the weakest predictor indicating that it is feasible to build a general predictive tool for protein corona formation on a large variety of DNA nanostructures of relevance across diverse biomedical applications.

**Figure 5.**
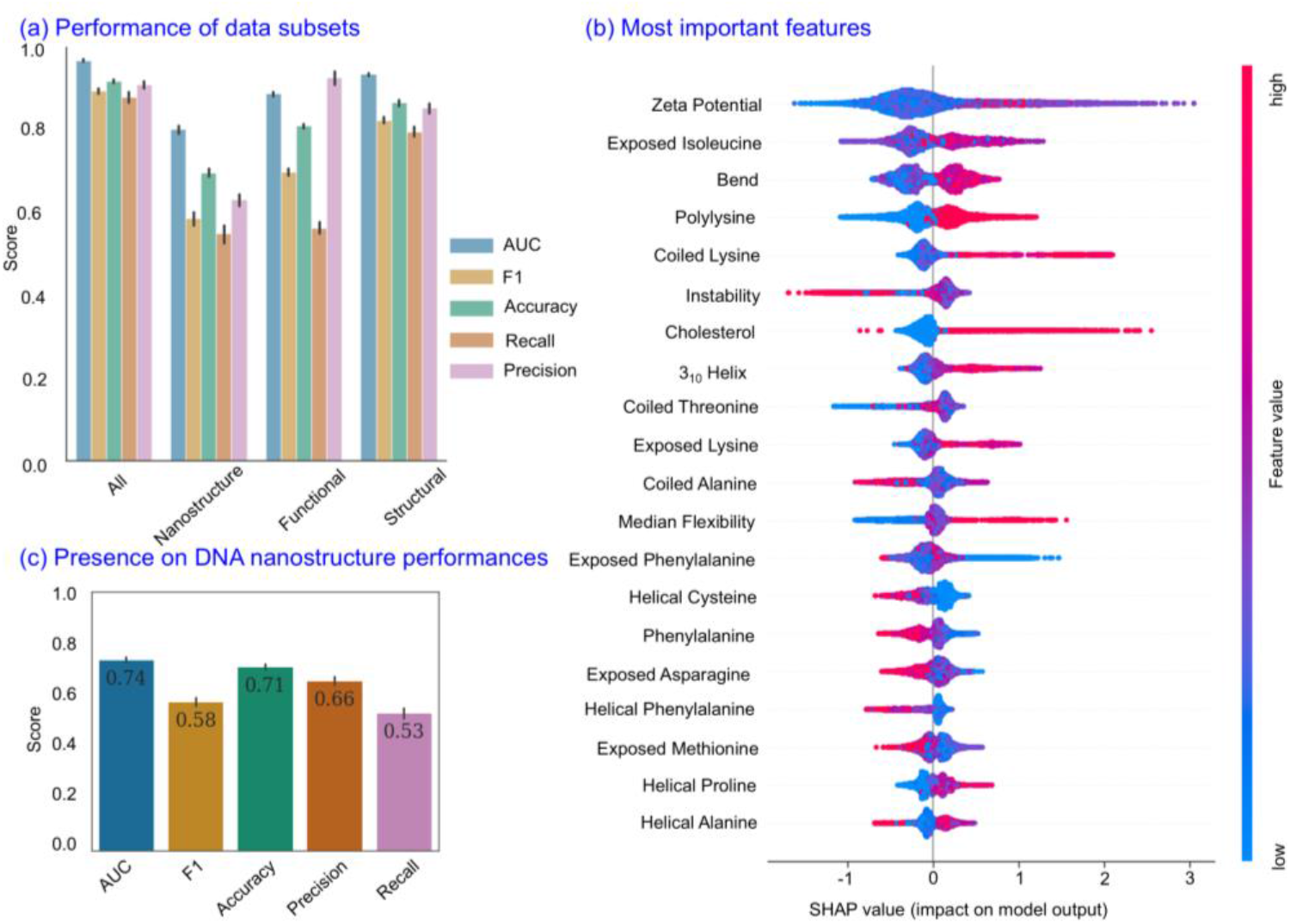
Model performance on data subsets and feature importances. **(a)** XGBoost model performance in classifying proteins as present in the corona across the different data subsets. **(b)** SHAP value plot of the 20 most important features for classifying proteins as present or absent. **(c)** Mean performances of XGBoost model trained individually on all nanostructures.

Having demonstrated the high performance of the model with all data available, we next utilized the model’s decision-making architecture to elucidate features governing protein adsorption to nanostructures. To interpret the importance values of different features, we calculated the Shadley Additive exPlanations (SHAP) for each feature^74^. Two of the five most important features were related to the modifications of DNA nanostructures: @PL coating and ζ-potential (**Fig. 5b**). Low values of ζ-potential and the absence of the @PL coating are both associated with proteins not being present in the corona. The presence of a cholesterol modified DNA strand is another influential design choice. Cholesterol had a particularly large impact on the model as its presence strongly and positively influenced whether a protein will be adsorbed. This finding further supports our conclusion that non-DNA modifications most significantly affect the protein corona composition on DNA nanostructures.

Several insights regarding protein properties governing adsorption can be gleaned from the SHAP values. For example, some protein secondary structures, like 3_10_ helices and bends, promoted positive predictions. A protein containing a large fraction of 3_10_ helices and bend secondary structures resulted in proteins being predicted as present in the corona. Different amino acids also had different effects on a protein’s likelihood to adsorb. For instance, a protein with a large fraction of exposed isoleucine residues corresponded with positive protein adsorption predictions, while a protein with a large fraction of exposed asparagine residues led to negative predictions of protein adsorption.

We then explored feature importances from the models trained on subgroups of DNA nanostructures with similar properties, i.e. are certain protein features better for protein adsorption on coated vs. uncoated nanostructures. When we trained models on individual nanostructures only, we observed that across nanostructures, there was a wide variance in model performance for each structure. Most individual nanostructure models could accurately classify a protein as present within the specific nanostructure’s corona with an accuracy ranging between 70%-80% (**Fig. 5c**), compared to the 92% predictive power when training the model on all DNA nanostructures together. Comparing the SHAP values for models trained on data from different structures can provide further insights into which protein properties are more likely to lead to adsorption across nanostructures of different design axes. But we had to consider the reduction in predictive power evident when we train models on individual nanostructures. So, to explore the effect of coating the nanostructures, we trained one model on all the nanostructures coated with @PL and separately trained another model on all nanostructures without a coating. We first validated these models and found that they maintained high levels of predictive power, with the models considering the uncoated and coated structures separately achieving accuracies of 94% and 90% respectively (**Fig. S8a, b)**. From the 20 most important protein and nanostructure features for each model, 5 protein properties (bend secondary structure, 3_10_ helix secondary structure, exposed isoleucine, coiled lysine, and exposed asparagine) are commonly influential to both coated versus uncoated nanostructure predictions, suggesting different factors cause the corona differences we see across coated and uncoated structures (**Fig. S8c, d)**. Among these 5 features, each had similar relationships regarding feature effect on protein corona presence across both models, indicating there are some conserved principles governing adsorption to DNA nanostructures regardless of the polymer coating.

Overall, our results demonstrate that several protein features like the amount of exposed isoleucine and phenylalanine promote protein adsorption onto DNA nanostructures of varying sizes, shapes, and modifications differently; while several other features like increased flexibility and decreased amino acid sequence length promote practically universal adsorption onto the DNA nanostructures. These results suggest it is possible to bias protein corona composition with nanostructure engineering, albeit with incomplete control over the entire proteome. Our results also suggest that engineering the biofluid itself, or perhaps pre-coating nanostructures with specific proteins, could enable greater control over DNA nanostructure physiochemical identity for subsequent use in a range of biofluids.

Lastly, we sought to understand the role of protein function on corona composition. We considered gene ontologies classified as molecular function as protein functions, which included ATP binding, actin binding, helicase activity, and many more. Specifically, for protein functions found in at least 23 of the 534 proteins identified across all of our trials and analyzed, we calculated the enrichment score for each protein function. We define enrichment score as the fold difference in the number of different proteins with that particular function present, as compared to the number of proteins with said function expected by chance to be in the corona. We find that of all tested protein functions, several are significantly enriched and depleted in proteins found within the corona (p-value <0.05) (**Fig. 6a**). These significantly enriched functions fall into two primary categories: (1) nucleic acid binding and (2) protein binding. As expected, DNA-binding is an enriched function, as well as RNA-binding, with a 1.7- and 1.2-fold increased diversity of corona proteins exhibiting DNA or RNA binding functions respectively, as opposed to the number expected by random. The most enriched functional group was the structural constituent of chromatin group, with a 2.2-fold enrichment of corona proteins being chromatin constituents. These proteins endogenously interact with DNA given they are part of the complex of DNA and proteins that make up chromatin, as such, it is reasonable that these protein functions would positively influence protein adsorption to nanostructures. The remaining functions comprise the second category: protein binding, suggesting proteins with a propensity to bind with other proteins are more likely to be present in nanostructures’ coronas. Specifically, protein heterodimerization activity exhibited a 1.9-fold enrichment, protein homodimerization activity a 1.3-fold enrichment, antigen binding a 1.2-fold enrichment, cadherin binding a 1.1-fold enrichment, and identical protein binding a 1.1-fold enrichment. We hypothesize that since the protein corona consists of several layers^76^, the outermost layer at any given time affects the interactions of remaining biofluid proteins. Therefore, the ability of a protein to be present in the outermost layer of a corona may be dependent on its ability to bind to and interact with other proteins forming the innermost corona layers. Interestingly, there was a large depletion in metal ion binding and ATP binding proteins, with both experiencing 0.7 and 0.7-fold depletions respectively. This occurs possibly because metal ion binding proteins have a greater affinity to the metal ions in the nanostructure buffer than the nanostructure itself. ATP binding proteins were depleted likely due to steric hindrances preventing nanostructures from compatibly interacting with the binding domain of the ATP binding proteins. We conclude that, of all considered protein features, a protein’s degree of nucleic acid binding and protein binding are the most influential in its ability to bind to DNA nanostructures.

**Figure 6.**
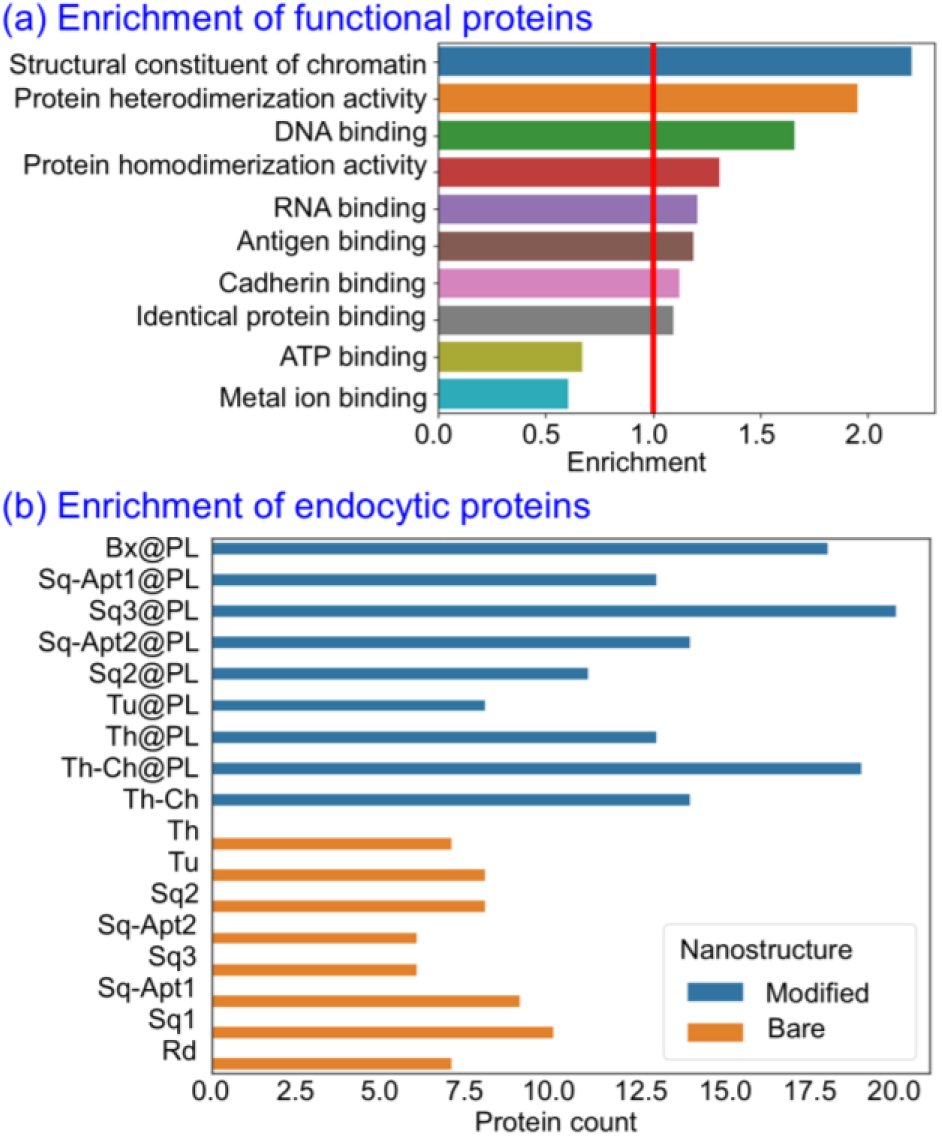
Differential adsorption of proteins with distinct functions in the corona. **(a)** Enrichment of functional protein families in nanostructure coronas. All enrichment and depletions are statistically significant (p-value < 0.05). **(b)** The number of endocytosis-associated proteins across all DNA nanostructures.

Having demonstrated statistically significant enrichment/depletions of various functional protein families, we explored if proteins associated with different biological processes can be differentially adsorbed to the surface of DNA nanostructures intentionally. Specifically, we examined whether the protein corona could be engineered by changing the different design parameters of the nanostructures. As a proof of concept, we explored engineering the nanostructure protein corona with proteins involved in endocytosis (gene ontology group GO:0006897) in different nanostructures. We selected this proof-of-principle experiment because controlling and better understanding endocytosis is important for cellular delivery of therapeutics and biophysical tools such as DNA nanostructures. We found that design axes like DNA nanostructure size and shape yielded minimal changes in the diversity of unique endocytic proteins adsorbed to the nanostructure. However, we observed an almost doubling in the diversity of endocytic proteins in the corona of modified (cationic polymer or cholesterol functionalized) nanostructures versus their bare, un-modified counterparts (**Fig. 6b**). Therefore, we hypothesize that cationic polymer coatings can enrich the protein corona for proteins associated with endocytosis. This effect is synergistic with the previously reported improvement in DNA nanostructure stability and ζ-potential increase when cationic polymers are used to coat nanostructures^25,36^. Our results suggest that other nanostructure surface modifications may be able to modulate the composition of the protein corona across other protein functional domains by introducing non-DNA modifications to DNA nanostructures.

## Discussion

Herein, we broadly surveyed the effect of DNA nanostructure design parameters on the composition of their protein coronas when incubated in human serum. We find that modulating structural properties of nanostructures (size, shape, etc.) can lead to differential adsorption of a minority of the overall proteins present in the corona, but that nanostructure design is largely less influential in driving protein corona composition than non-DNA nanostructure surface modification with polymers or cholesterol. Design features of the nanostructures only slightly bias the adsorption of different proteins, with at most 36% difference in protein corona composition between the two most dissimilar DNA-only nanostructures. Conversely, the addition of non-DNA modifications (cholesterol and cationic polymer coatings) leads to the most pronounced changes in coronas, with up to 52% difference in protein corona composition between the polymer-coated nanostructure relative to its uncoated counterpart. We hypothesize this non-DNA modification driven increase in protein corona compositional diversity is due to the hydrophobicity of cholesterol and the cationic charge of the polymer attracting new classes of proteins to the corona, thereby adding to those already binding to hydrophilic, negatively charged DNA. Our work demonstrates the potential to engineer the nanostructure corona using both non-DNA modifications to the nanostructure and, to a lesser extent, by modifying the DNA nanostructure itself.

To further promote corona engineering, we developed two explainable machine learning models that predict whether a protein will be present/absent or enriched/depleted from a given nanostructure’s corona. These models are first-of-their-kind tools enabling the accurate prediction of the protein corona on any given DNA nanostructure with up to 92% accuracy. In addition, by utilizing an explainable algorithm, the models offer valuable insights into the factors governing the adsorption of proteins, both in the case of proteins that ubiquitously bind all DNA nanostructures versus proteins that are unique to certain nanostructure constructs. Therefore, we envision our model will enable researchers to further reclaim the programmability of DNA nanostructures that is typically lost with spontaneous protein corona formation. With model-based predictability of nanostructure protein corona composition, researchers can account for spontaneous protein adsorption prior to experimentation. Furthermore, our approach can support the design of nanostructures with designer coronas. Utilizing the knowledge of what nanostructure features and protein properties drive protein corona formation, both independently and in concert, it is possible to intelligently design nanostructures to bias the corona favorably. Harnessing the protein corona can enhance the efficiency of nanostructures *in vivo* by utilizing protein properties to favorably improve circulation time, anatomical targeting, biocompatibility, and cellular uptake. We anticipate this work will serve as a step toward the future of DNA constructs as nanomedicines, biosensors, and general tools for probing and manipulating biological organisms.

While we intend for this work to improve the engineering of DNA nanostructures for *in vivo* applications, we also acknowledge that several limitations and hurdles remain. Most envisioned applications of DNA nanostructures *in vivo* are intended for nanostructure end-fate either on the cell membrane or within the cell. Therefore, while our study supports a better understanding of nanostructure physiochemical identity in human circulation, to thoroughly understand the role of the corona on nanostructure intracellular fate, this study bears repeating in other biological milieus like the cytoplasm. We expect results obtained using our enrichment classifier will be largely generalizable to other biological fluids, and other nanostructures, but this assumption needs to be experimentally verified. In addition, studies need to be performed with sequential incubation into different biologically relevant milieu. When a nanomaterial with pre-adsorbed protein corona enters circulation, certain *in vivo* proteins may adsorb and displace the original pre-adsorbed proteins as per the Vroman effect^77^. Nanostructures *in vivo* may traverse through numerous unique environments sequentially before reaching their intended target, and each of these environments drives the formation of protein coronas with unique identities. We expect this dynamic evolution of the corona would bias the corona’s final composition to proteins with greater binding affinity to components of the nanostructure-corona complex, like nucleic acids and proteins. Lastly, while these are all factors that can be determined through further experiments and their integration with machine learning, there are inherent limitations to using machine learning algorithms altogether. Our algorithm provides insight into which proteins are likely to adsorb into the nanostructure protein corona, but it is generally unable to provide mechanistic insights into how and why. For such mechanistic and structural insights, further studies using high-resolution imaging and biochemical assays are necessary as a complement to the predictive power of machine learning. Taken together, we anticipate our study can enhance the effectiveness of DNA nanostructures *in vitro* and *in vivo*, and also inspire further approaches and inquiries into the important question of how, why, and which proteins adsorb to DNA nanostructures.

## Acknowledgments

The authors thank Yifeng Shi, Myoungseok Kim, Durham Smith, and Arjun Banerjee for providing some of the DNA nanostructures and the UCSD Proteomics Facility for analyzing samples. JH was supported by NIH training grant (T32GM146614). GT was supported by Society of Hellman Fellows Fund, UC Berkeley, NSF CAREER (2240000) and NSF POSE Phase 2 (2346048) awards. We acknowledge support of a Burroughs Wellcome Fund (BWF) Career Award at the Scientific Interface (CASI) (MPL), a Dreyfus foundation award (MPL), the Philomathia foundation (MPL), an NSF CAREER award 2046159 (MPL), an NSF CBET award 1733575 (to MPL), a CZI imaging award (MPL), a Sloan Foundation Award (MPL), a McKnight Foundation award (MPL), a Simons Foundation Award (MPL), a Moore Foundation Award (MPL), a Brain Foundation Award (MPL), and a polymaths award from Schmidt Sciences, LLC (MPL). MPL is a Chan Zuckerberg Biohub investigator, and a Helen Wills Neuroscience Institute Investigator. RC was supported by the NSF PRFB (2305663) and BWF PDEP awards.

## Author contributions

JH synthesized, characterized, and purified DNA nanostructure library and developed machine learning models. RC performed experiments to obtain corona composition. JH wrote the manuscript with contributions from GT, RC, and MPL. MPL and GT conceived the study. GT guided the project.

