## Supporting Information for "AI-based Prediction of Protein Corona Composition on DNA Nanostructures"

### Table of Contents


**1 Materials and Methods ... 4**

1.1 Data and Code Availability ... 4

1.2 Nanostructure Synthesis ... 4

1.2.1 Box (Bx) ... 4

1.2.2 Tube (Tu) ... 4

1.2.3 Square (Sq) ... 4

1.2.4 Rod (Rd) ... 4

1.2.5 Tetrahedron with and without cholesterol (Th and Th-Ch) ... 5

1.3 Cationic Polymer Coating ... 5

1.4 Nanostructure Characterization Protocols ... 5

1.4.1 Agarose Gel Electrophoresis ... 5

1.4.2 Transmission Electron Microscopy Imaging ... 5

1.4.3 Atomic Force Microscopy Imaging ... 5

1.4.4 Zeta (ζ)Potential Measurements ... 6

1.5 Database Construction ... 6

1.5.1 Nanostructure Features ... 6

1.5.2 Protein Features ... 6

1.5.3 Protein Presence Classification ... 6

1.5.4 Protein Enrichment Classification ... 6

1.6 Protein Corona Analysis ... 7

1.6.1 Corona Extraction Pull-Down Assay ... 7

1.6.2 Gel Electrophoresis Shift Assay ... 7

1.6.3 In-gel Protein Proteolysis ... 7

1.6.4 Liquid-Chromatography Mass Spectrometry ... 8

1.6.5 UHPLC-MS/MS Abundance Calculations ... 8

1.7 Computational Analysis ... 8

1.7.1 Corona Similarity Calculations ... 8

1.7.2 Protein Interactions ... 9

1.7.3 Protein Property Differences ... 9

1.7.4 Protein Function Enrichment ... 9

**2 Parameter Space Schematic (S1) ... 10**

**3 Protein Corona Protocol Validation (S2) ... 11**

**4 Protein Corona Abundances ... 12**

4.1 Heatmap of protein corona relative to human serum (S3) ... 12

4.2 Volcano plots of proteins' relative abundance (S4) ... 13
4.3 Heatmap of unique corona proteins associated with enriched biological processes (S5) ... 14
4.3 Heatmap of protein corona with magnetic bead subtracted (S6) ... 15
**5 Machine Learning Model Architecture Validation (S7) ... 16**
**6 Model Performance on Data Subsets (S8) ... 17**
**7 Nanostructure Characterizations ... 18**
7.1 Rod (Rd) (S9) and coated rod (Rd@PL) (S10) ... 18
7.2 Tube (Tu) (S11) ... 19
7.3 Box (Bx) (S12) ... 19
7.4 Tile (Sq1) (S13) and aptamer modified tile (Sq-Apt) (S14) ... 20
7.5 Th (Th) (S15) and cholesterol conjugated Th (Th-Ch) (S15) ... 21
**8 Nanostructure Designs ... 22**
8.1 Rod (Rd) (S16) ... 22
8.2 Tube (Tu) (S17) ... 23
8.3 Square (Sq) (S18) ... 24
**9 Nanostructure Feature Quantifications (Table S1) ... 25**
**10 Model Features and Importance Values (Table S2) ... 26**
**11 DNA Sequences (Table S3) ... 77**
**12 References ... 131**

### 1 Materials and Methods

#### 1.1 Data and code availability:

All data and code used to draw the conclusions in both the main and supplementary text is available on Github (https://github.com/jaredhuzar/DNANano_ProteinCorona_ML).

#### 1.2 Nanostructure Synthesis

Single-stranded M13mp13 DNA was purchased from Bayou Biolabs (Catalog: P-107) at 1 µg/µL and stored at -20°C in 10 mM Tris-Cl and 1 mM EDTA at pH=8. DNA strands listed in table S3 were purchased from Integrated DNA Technologies and stored in 1x TE buffer. Structures Bx, Tu, Sq, and Rd were synthesized by adding a molar excess of staples to the DNA scaffold before being mixed in the following buffers with the following annealing ramps. Th and Th-Ch samples are scaffoldless and were synthesized by adding their respective 4 strands together in 1:1 molar ratios. After annealing, all samples were stored at 4°C. Bx, Tu, and Sq structures were purified from excess staples using either 100 kDa or 50 kDa Amicon Ultra-0.5 ml centrifugal filters (Millipore Corporation).

##### 1.2.1 Box (Bx)

The sample was synthesized following the previously published protocol[1]. Scaffold and staples were mixed in 1x Tris-Acetate-EDTA (TAE) buffer supplemented with 16 mM $MgCl_2$. Samples were then incubated at 75°C for 15 minutes, followed by a temperature ramp of -0.1°C/1.5 minutes until the reaction temperature reached 60°C, at which point the ramp was slowed down to -0.1°C/6 minutes until the temperature reached 20°C.

##### 1.2.2 Tube (Tu)

The sample was synthesized by mixing the scaffold and staples in 1x Tris-EDTA (TE) buffer supplemented with 12.5 mM $MgCl_2$. The reaction mixture was then incubated at 95°C for 2 minutes followed by a temperature ramp to 75°C at a rate of -0.1C/6 seconds, and then cooled from 75°C to 20°C at a rate of -0.1°C/12 seconds. The sample was purified using gel extraction. The band in an agarose gel corresponding to the desired tube structure was excised using a razorblade and placed inside a Freeze 'N Squeeze DNA Gel Extraction Spin Column (Bio-Rad, Catalog No: 7326165), the gel was then scrambled using tweezers, placed at -20°C for >5 minutes, and centrifuged at 7,000 RCF for 5 minutes to recover the purified sample.

##### 1.2.3 Square (Sq)

The Sq samples were synthesized following the previously published protocol[2]. Scaffold and staples were mixed in 1x TE buffer supplemented with 12.5 mM $MgCl_2$. The reaction mixture was then incubated at 90°C for two minutes followed by a temperature ramp from 90°C to 20°C at a rate of -0.1°C/6 seconds.

##### 1.2.4 Rod (Rd)

The Rd samples were synthesized following the previously published protocol[3]. Scaffold and staples were mixed in 1x TAE buffer supplemented with 20 mM $MgCl_2$. The reaction mixture was then incubated at 80°C for 2 minutes followed by a temperature ramp from 80°C to 14°C over 15 hours.

###### 1.2.5 Tetrahedron (Th) and cholesterol modified tetrahedron (Th-Ch)

Samples were synthesized following the previously published protocol[4]. Samples were mixed in 20 mM Tris-HCl buffer supplemented with 5 mM $MgCl_2$. The reaction mixture was incubated at 95°C for 5 minutes followed by a temperature ramp to -1.5C/1 minute to 5C, and then -1C/1 minute to 4°C.

#### 1.3 Cationic Polymer Coating

The poly(L-lysine) backbone and poly(ethylene glycol) side-chains (PLL-g-PEG) polymer was ordered from Fisher Scientific (Catalog No.: NC1728626) and supplied by Nanosoft Polymers. The polymer consisted of 200 repeating units of PLL and was grafted with 5 kDa molecular weight PEG with 20% PEG substitution. DNA nanostructures were incubated with the polymer at room temperature for >30 minutes. The polymer was added to a final concentration of 0.16 µg/mL.

#### 1.4 Nanostructure Characterization Protocols

###### 1.4.1 Agarose Gel Electrophoresis

All nanostructures except Th and Th-Ch were analyzed using 1% wt/vol agarose gel ran at 75 V for 90 minutes with a running buffer of 0.5x Tris-Borate-EDTA (TBE) and 12.5 mM $MgCl_2$. A 1 kilobase pair ladder purchased from New England Biolabs (Catalog No: N3232L) served as the reference. Th-Ch was analyzed using 4% wt/vol agarose gel with a running buffer of 1x TAE. The gel was run at 60 V for 120 minutes. A 100 basepair ladder purchased from New England Biolabs (Catalog No: N3231S) served as the reference. All samples were stained using Ethidium Bromide (Thermo Scientific, Catalog No: 17896) and bands were visualized using a ChemiDoc™ Imaging System (Bio-Rad, Catalog No: 12003153)

###### 1.4.2 Transmission Electron Microscopy Imaging

2-5 µL of DNA nanostructures in their folding buffer were deposited onto glow-discharged carbon coated copper grids (Electron Microscopy Sciences, Catalog No: CF300-Cu-Ul). They were then incubated for >5 minutes at room temperature, before being blot dried with filter paper. They were then washed 3 times in water droplets. Micrographs of the Rd structure reported in figures S11 and S12 were recorded using a low-voltage electron microscope (LVEM5, Delong Instrument, Montreal, Quebec, Canada). Electron micrographs of the box structure (Bx) and tube structure (Tu) in figures S13 and S15 were recorded using an FEI TECNAI 12 TEM. Bx and Tu structures were deposited following identical procedures as described above, but with an additional staining step for one minute using 2% Uranyl Acetate after washing with water.

###### 1.4.3 Atomic Force Microscopy Imaging

Samples were imaged using Bruker FastScan Bio (Bruker, USA) in ScanAsyst Air mode. Samples were first diluted to 1 nM in their respective folding buffers before being deposited on freshly cleaved mica flakes. Samples were incubated on the mica for >5 minutes before being washed with water and subsequently dried using an air-pump-bulb. Images were processed by flattening with the Nanoscope analysis software to subtract out effects of mica imperfections.

###### 1.4.4 Zeta (ζ)Potential Measurements

All DNA nanostructures were suspended in 5 mM $MgCl_2$ buffer, except the rod (Rd) and the DNA tile with biotin (Sq1), which were suspended in 20 and 12.5 mM $MgCl_2$, respectively. Nanostructures' ζ potential was measured using a Nano ZS (Malvern Panalytical); ζ potential measurements were performed with nanostructures at a final concentration of 100 pM and 0.1 mM NaCl to improve conductivity and analyzed by the Smoluchowski approximation model.

#### 1.5 Database Construction

###### 1.5.1 Nanostructure Features

Quantification of nanostructure size was achieved by counting the approximate number of nucleotides in each structure. The theoretical volume and surface area was quantified based on the a distance of 0.34 nm per nucleotide and 2.0 nm helical diameter[5]. We quantified the shape of a nanoparticle by estimating its aspect ratio, in addition we added a binary metric for encoding the dimensionality of the structure (3D vs 2D). The number of cavities and vertices for each structure was estimated from the designs. For surface modifications, we counted the number of cholesterol molecules, and separately the presence of absence of aptamers in the structure. Finally, we included a binary metric for whether the nanostructure was coated with polylysine. ζ potential was measured using an electrophoretic light scattering zetasizer (Nano ZS, Malvern Panalytical). For structures with no ζ potential data, we imputed values calculated based on similar structures' measured ζ potentials.

###### 1.5.2 Protein Features

Structural data was obtained by scraping from the UniProt database[6] (UP000005640), from the Quantiprot Python package[7], by obtaining structural prediction from NetSurf 2.0[8], and prediction of DNA binding by iDNA-Prot[9]. Functional information was obtained from GO annotations on the UniProt database. Proteins greater than 5000 amino acids long were removed due to NetSurf 2.0 limitations. Similar proteins were also removed. Finally, the number of binary interactions each protein has with other proteins was obtained through the UniProt database. This count served as a metric for interactivity. For a comprehensive list regarding the specific metrics please see Table S1.

###### 1.5.3 Protein Presence Classification

Proteins were classified as present in the corona if, after subtracting its abundance in the magnetic bead only sample, there remained a >0 amount of protein found in the corona. Therefore, proteins were classified as present even if they were found in lower abundance than in serum alone.

###### 1.5.4 Protein Enrichment Classification

Proteins were classified as enriched in the corona if, after subtracting its abundance in the magnetic bead only sample, there remained greater amount of protein found in the corona than in sera alone. Therefore, proteins were classified as enriched only if they were found in greater abundance in the corona than in serum alone.

#### 1.6 Protein Corona Analysis

##### 1.6.1 Corona Extraction Pull-Down Assay

Streptavidin-coated magnetic beads (Dynabeads™ MyOne Streptavidin™ T1, Catalog No: 65601) were obtained from ThermoFischer Scientific. The concentration of pooled human serum (Thomas Scientific, Catalog No: C817B49) was estimated using a Qubit Protein Broad Range Assay (ThermoFisher Scientific, Catalog No: A50668) and diluted to a working stock (10 mg/mL in 1× PBS). Next, 10 µL of the beads were washed three times with 100 µL of 1× PBS. The washed beads were resuspended with 50 µL of diluted pooled human serum and incubated at 37°C for 1h to pre-coat the magnetic beads with proteins. Proteins non-specifically bound to the magnetic beads were removed and the protein-coated magnetic beads were washed 3× with 50 µL of 2× PBS. The beads were resuspended with fresh 50 µL of diluted human serum and 50 µL of a singular biotinylated-DNA nanostructure (50 pM final nanostructure concentration) and incubated at 37°C for 1h. During the incubation, the nanostructures were subsequently immobilized to the streptavidin-coated magnetic beads through non-covalent biotin-streptavidin interactions. Magnetic beads were then separated from unbound serum proteins and washed three times with 1× PBS to further remove non-specifically bound proteins from the nanostructures. Formation of the protein corona on each nanostructure was performed in triplicate.

##### 1.6.2 Gel Electrophoresis Shift Assay

After the pull-down assay, the streptavidin-coated magnetic beads attached to the biotinylated-DNA nanostructures and their protein coronas were resuspended with 10 µL of 1× PBS and 10 µL of Laemmli sample buffer (138.9 mM Tris-HCl (pH 6.8), 2.2% LDS, 22.2% (w/v) glycerol, 0.01% (w/v) bromophenol blue, 1.43 M 2-mercaptoethanol; BioRad). Adsorbed proteins were denatured from DNA nanostructures by incubating the mixtures at 95°C for 10 min. The desorbed corona proteins were then separated with 4-20% SDS-PAGE at 110 V for 70 mins, stained with 1× Flamingo Fluorescent Stain (BioRad, Catalog No: 1610490 ), imaged with a GE Typhoon FLA 9000 Imaging Scanner, and gel band intensities were analyzed with ImageJ.

##### 1.6.3 In-gel Protein Proteolysis

Protein bands (~ 75-25 kDa) were excised from the SDS-PAGE gels, diced into 1mm × 1 mm cubes and unstained 3× by first washing with 100 µL of 100 mM ammonium bicarbonate ($NH_4HCO_3$) for 15 min followed by an addition of 100 µL of acetonitrile for 15 min. The supernatants were removed, and the gel pieces were dried with a SpeedVac. Samples were reduced by incubating the gel pieces with 200 µL of 10 mM DTT in 100 mM $NH_4HCO_3$ at 56°C for 30 min. After cooling to ambient temperature, the supernatants were removed and replaced with 200 µL of 55 mM IAA in 100 mM $NH_4HCO_3$ and the samples were incubated in the dark for 20 min. The supernatants were then removed, and the gel pieces were washed 1× with 200 µL of 100 mM $NH_4HCO_3$ for 15 min. Samples were then dehydrated with 200 µL acetonitrile and dried with a Speedvac. For protein digestion, enough solution of ice-cold trypsin (0.01 µg/µL), in 50 mM $NH_4HCO_3$, was added to cover the gel pieces and the samples were placed on ice for 30 min. After complete rehydration of the gel pieces, the trypsin solutions were removed and replaced with 50 mM $NH_4HCO_3$ and left overnight at 37°C. The peptides were extracted twice by adding 50 µL of 0.2% formic acid/5% acetonitrile and vortexing for 30 min. The supernatants, containing the peptides, were collected. Residual peptides were further extracted from the SDS-PAGE gel pieces by adding 50 µL of 0.2% formic acid/50% acetonitrile and vortexing for 30 min. The supernatant was collected and pooled with the peptides from the initial extraction and dried by SpeedVac.

##### 1.6.4 Liquid-Chromatography Mass Spectrometry

Peptides were redissolved in 10 µL of 5% formic acid and analzyed by ultra-high performance liquid chromatography coupled with tandem mass spectrometry (UHPLC-MS/MS) using nano-electrospray ionization (nano-ESI). The nano-ESI was performed using a timsTOF Pro 2 hybrid mass spectrometer (Bruker) interfaced with nanoscale reversed phased UHPLC (Evosep One), which utilized 10 cm × 150 µm reverse-phase columns packed with 1.5 µm C18-beads (PepSep, Bruker). The columns were connected with a fused silica ID emitter (10 µm ID; Bruker) inside a nano-ESI ion source (Captive spray source, Bruker). Mobile phase A was 0.1% formic acid and mobile phase B was composed of 0.1% formic acid/99.9% acetonitrile. The timsTOF Pro 2 MS was operated in the PASEF mode for standard proteomics. The values for mobility-dependent collision energy ramping were set to 95 eV at an inversed reduced mobility ($1/k_0$) of 1.6 V・s/cm$^2$ and 23 eV at 0.73 V・s/cm$^2$. Collision energies were linearly interpolated between these $1/k_0$ values and kept constant above or below. No merging of TIMS scans was performed. Target intensity per individual PASEF precursor was set to 20,000. The scan range was set between 0.6 and 1.6 V・s/cm$^2$ with a ramp time of 166 ms. Fourteen PASEF MS/MS scans were triggered per cycle (2.57 s) with a maximum of seven precursors per mobilogram. The precursor ions in an *m/z* range between 100 and 1700, with charge states ≥3+ and ≤8+ were selected for fragmentation. Active exclusion was enabled for 0.4 min (mass width 0.015 Th, $1/k_0$ width of 0.015 V・s/cm$^2$). Protein identification and label free quantification was carried out using Peaks Studio X (Bioinformatics Solutions, Inc.) against the UniProtKB *Homo sapiens* database (UP000005640).

##### 1.6.5 UHPLC-MS/MS Abundance Calculations

Spectral counts for triplicate nanostructure samples were obtained from the UHPLC-MS/MS measurements. Proteins with spectral counts <2 were filtered out. Spectral counts were normalized by the total spectral count for each sample to obtain relative abundances for each protein within each sample. To compare protein abundances between nanostructure coronas and control samples, average relative abundances were used to compute the fold change in protein abundances between the nanostructure coronas and diluted sera. Proteins were considered enriched in the corona if their $\log_2$(fold change) was greater than 0 (i.e.: proteins are considered enriched if they are found at greater abundance in the corona than in serum). Student's *t*-test was calculated for the statistical analysis between protein relative abundance arrays in the nanostructure coronas and the sera controls; the -log(*p*-values) were thus calculated for Volcano plots. To subtract out the effect of the protein corona on the magnetic beads from the measured protein corona on the captured DNA nanostructures, spectral counts of proteins found in nanostructure coronas were subtracted by the respective average spectral count in the magnetic bead sample. Fold changes were also calculated with the relative abundances that factored out the magnetic bead values. Proteins were also considered uniquely present in the nanostructure coronas if spectral counts were identified in the protein corona samples but not in the sera control; for these specific proteins, enrichment values could not be calculated due to the absence of spectral counts in the sera.

#### 1.7 Computational Analysis

##### 1.7.1 Corona Similarity Calculations

Nanostructure protein corona similarity values were calculated by:

$$(1) \qquad \text{Similarity} = 100 \times \frac{|A \cup B|}{|A \cap B|}$$

Where ‘A’ is the set of proteins present in one nanostructure’s corona, and ‘B’ is the set of proteins present in another nanostructure’s corona.

##### 1.7.2 Protein Interactions

All proteins that were present in all nanostructures’ coronas were queried in homo sapiens using the STRING database. Active interaction sources selected were: experiments, databases, co-expression, neighborhood, gene fusion, and co-occurrence. Minimum required interaction score was 0.900. Network type is full STRING network, meaning edges represent both functional and physical protein associations. Nodes were colored based on MCL clustering with an inflation parameter of 3. Proteins with no interactions were removed.

##### 1.7.3 Protein Property Differences

The difference of protein property between proteins found to be universally adsorbed on nanostructure coronas versus universally absent was determined using the t-test for the means of two independent samples. This assumes the two populations have identical variances.

##### 1.7.4 Protein Function Enrichment

The enrichment of functional proteins was defined as the fold difference in number of different proteins with that particular function present as compared to the number expected by random. It was calculated as:

$$\text{(2)} \qquad \text{Enrichment} = \frac{nN}{kM}$$

Where ‘n’ is the number of proteins that are both abundant and have the function of interest, ‘N’ is the total number of proteins, ‘k’ is the total number of proteins with the function of interest, and ‘M’ is the total number of proteins abundant in the corona. The p-values were calculated based on the cumulative distribution function of the hypergeometric distribution.

#### 2 Parameter Space Schematic

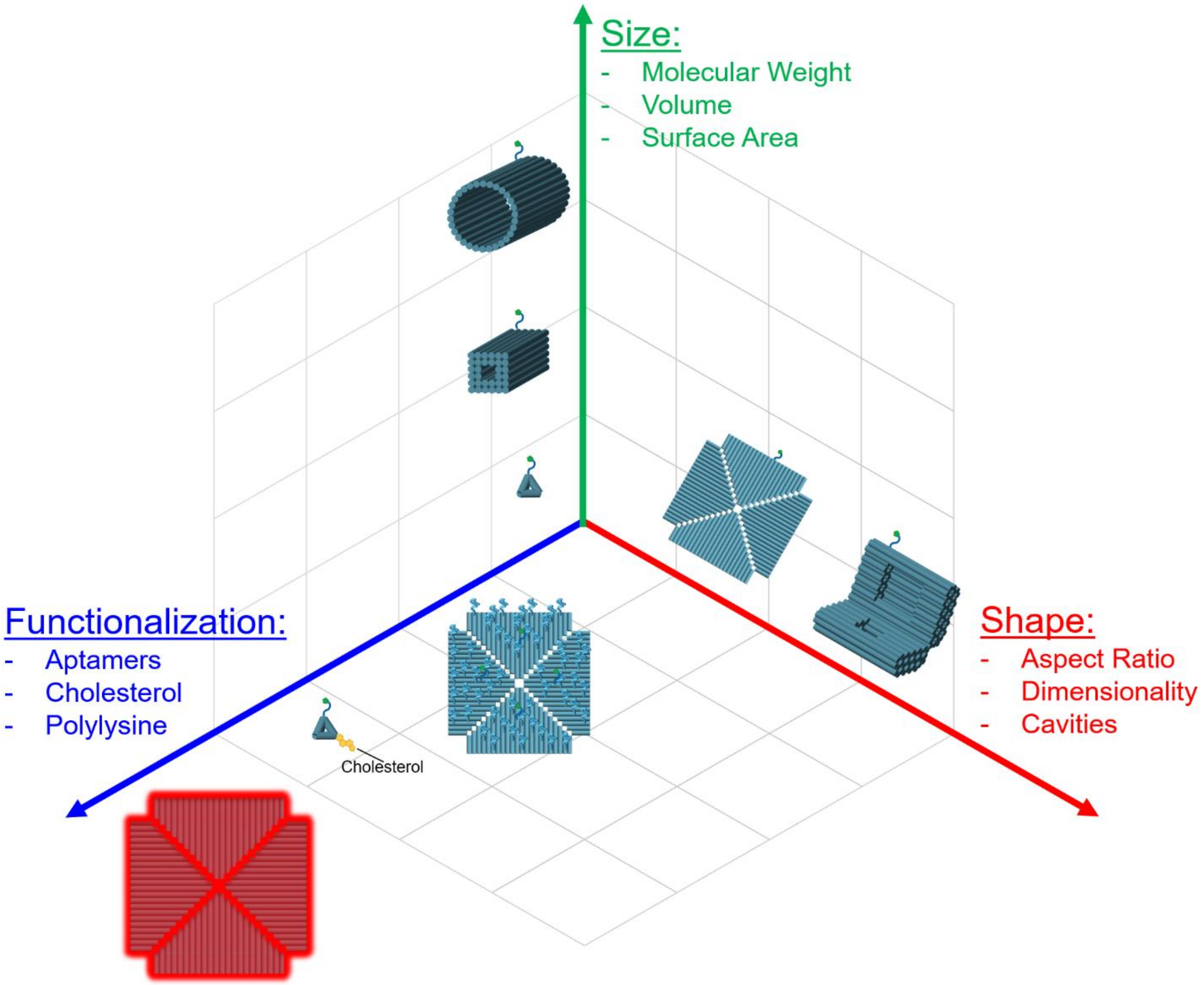


**Fig. S1.** Schematic representation of design parameters space of DNA nanostructures.

### 3 Protein Corona Protocol Validation

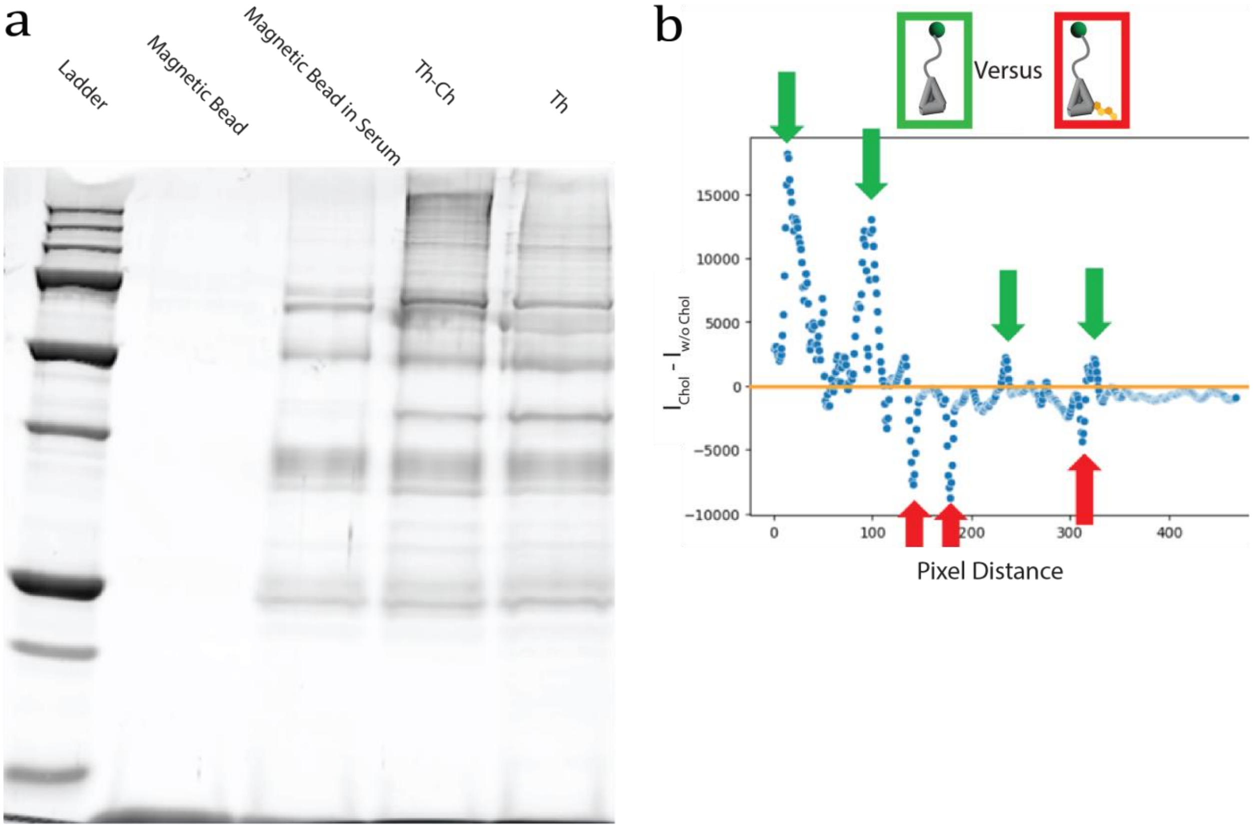


**Fig. S2. (a)** SDS-PAGE analysis of Th nanostructures comparing magnetic bead control with and without human sera. **(b)** Gel pixel intensity analysis on the SDS-PAGE of the cholesterol-modified and unmodified Th.

### 4 Protein Corona Abundances

#### 4.1 Heatmap of protein corona relative to human serum

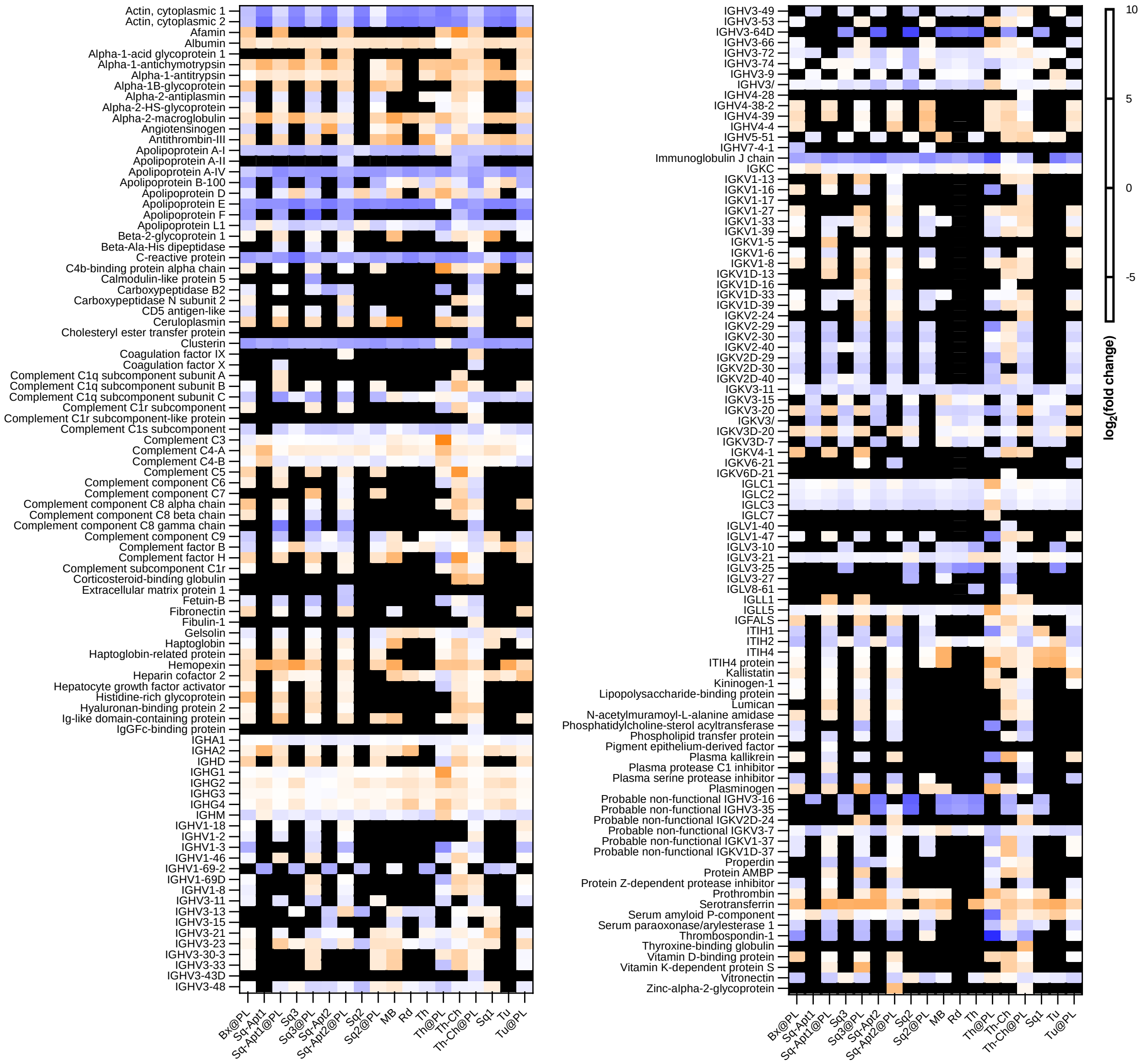


**Fig. S3.** Heatmap of protein expression change of the nanostructures' and magnetic beads' (MB) protein corona relative to the human serum control. Fold changes were quantified using average relative abundances of biological triplicates. Protein enrichment and depletion on the nanostructures' corona, where $\log_2$(fold change) >0 and $\log_2$(fold change) <0, respectively, is correspondingly shown with orange and blue cells. Black cells correspond to incalculable fold changes, where protein relative abundance is found solely in the nanostructures' protein corona or human serum. The heatmap was divided into 2 portions to optimize text visibility.

#### 4.2 Volcano plots of proteins' relative abundance

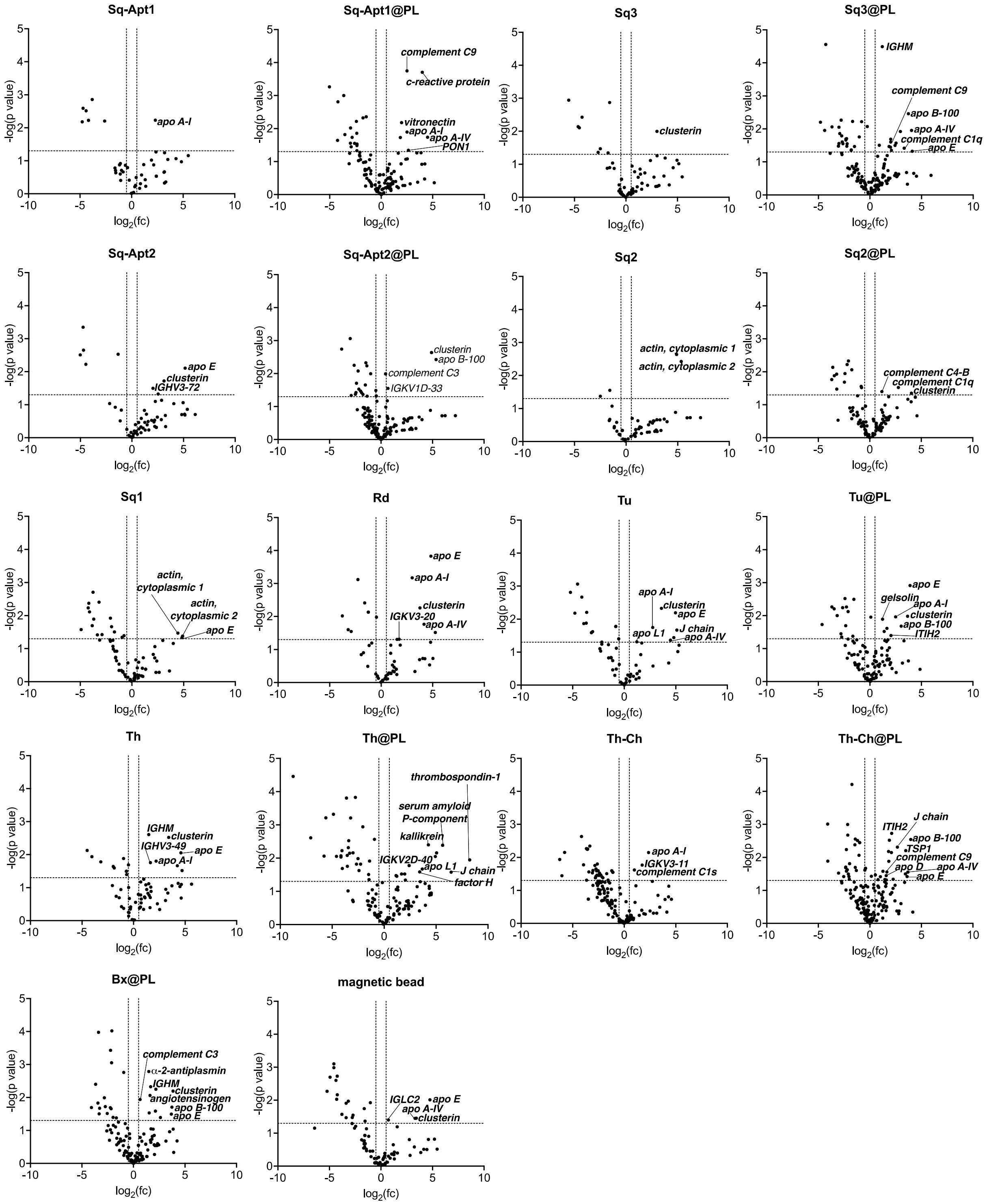


**Fig. S4.** Volcano plots depicting the statistical significance (-$\log_{10}$(*p*-value)) versus the magnitude of change ($\log_2$(fold-change)) of protein relative abundances (RA) across all 17 DNA nanostructures and the magnetic bead. MB spectral counts (SC) were not subtracted.

#### 4.3 Heatmap of unique corona proteins associated with enriched biological processes

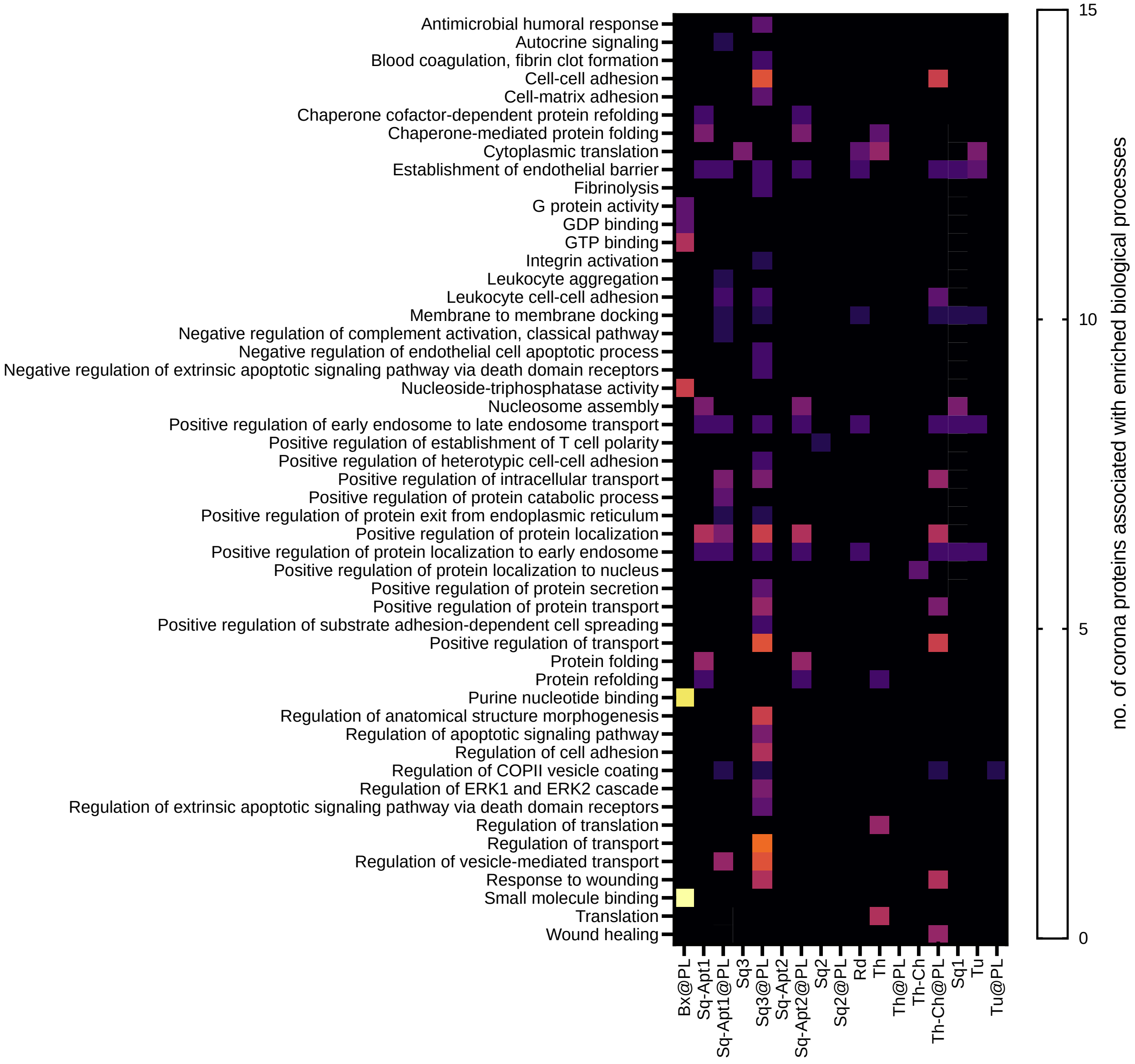


**Fig. S5.** Heatmap illustrating gene ontology analysis of the unique corona proteins associated with enriched biological processes. Each biological process listed had a maximum false discovery rate of ≤0.05.

#### 4.4 Heatmap of protein corona with magnetic bead subtracted

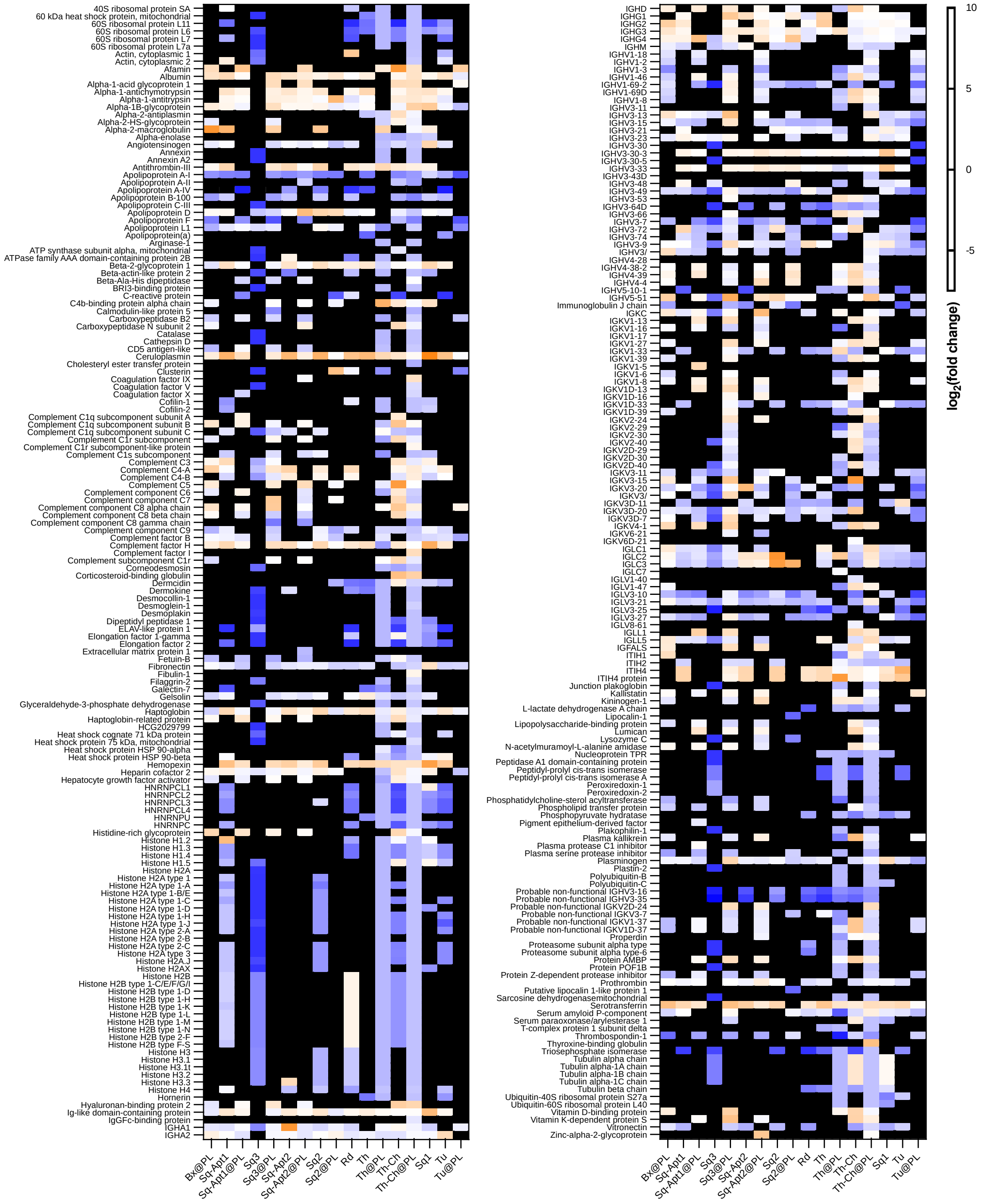


**Fig. S6.** Heatmap of the magnitude of protein expression changes in the nanostructures' protein corona, relative to the human serum control, with the SC from the MB subtracted. Protein enrichment on the nanostructures' corona, where $\log_2$(fold change) >0, is depicted with blue cells, and protein depletion in the corona, where $\log_2$(fold change) <0, is shown with orange cells. Black cells correspond to incalculable fold changes, where protein relative abundance is found either in the nanostructures' protein corona or the human serum control. The heatmap was divided into 2 portions to optimize text visibility.

### 5 Machine Learning Model Architecture Validation

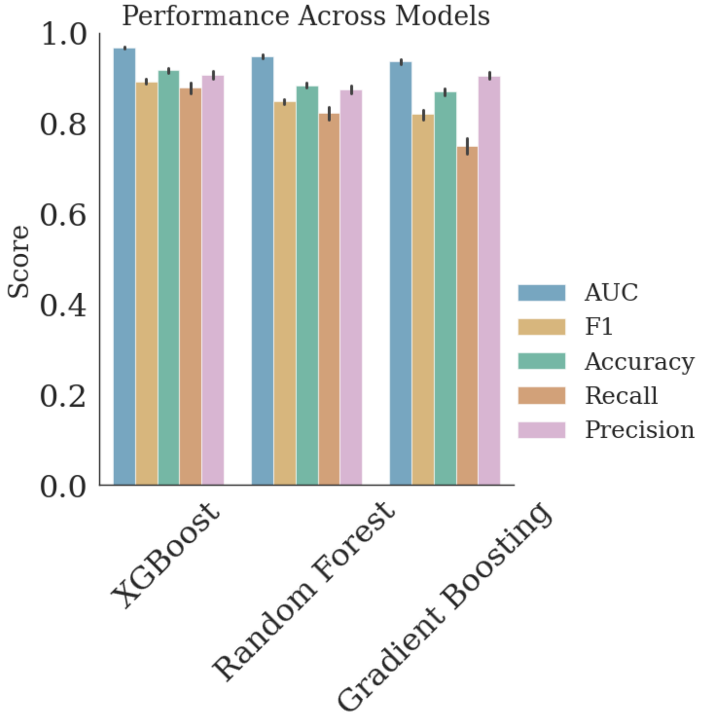


**Fig. S7.** Protein presence classification performance across various model architectures.

### 6 Model Performance on Data Subsets

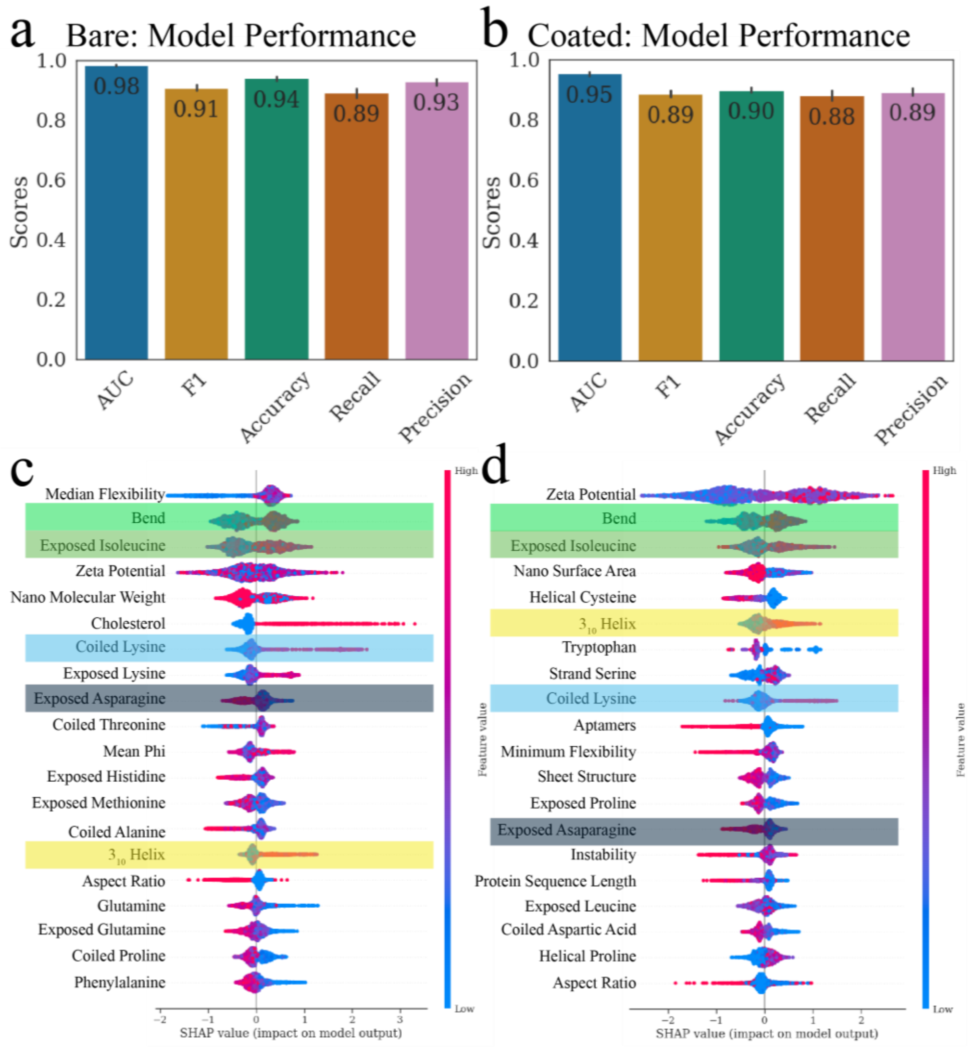


**Fig. S8. (a-b)** Mean model performance across 10-fold split. **(a)** Bare nanostructures only. **(b)** Coated nanostructures only. **(c-d)** SHAP values of 20 most important features for protein presence prediction. **(c)** Bare nanostructures. **(d)** Polymer coated nanostructures.

### 7 Nanostructure Characterizations

#### 7.1 Rod (Rd) and coated rod (Rd@Pl)

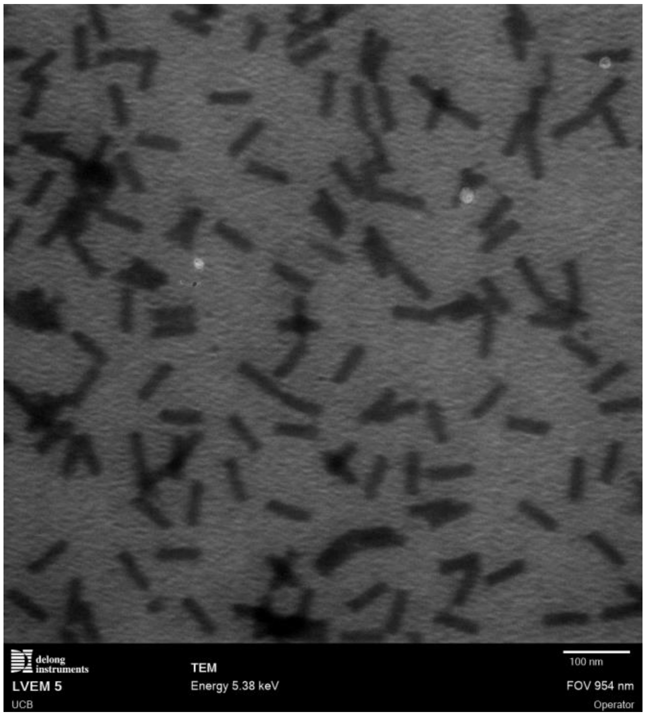


**Fig. S9.** TEM characterization of 32HB (Rd).

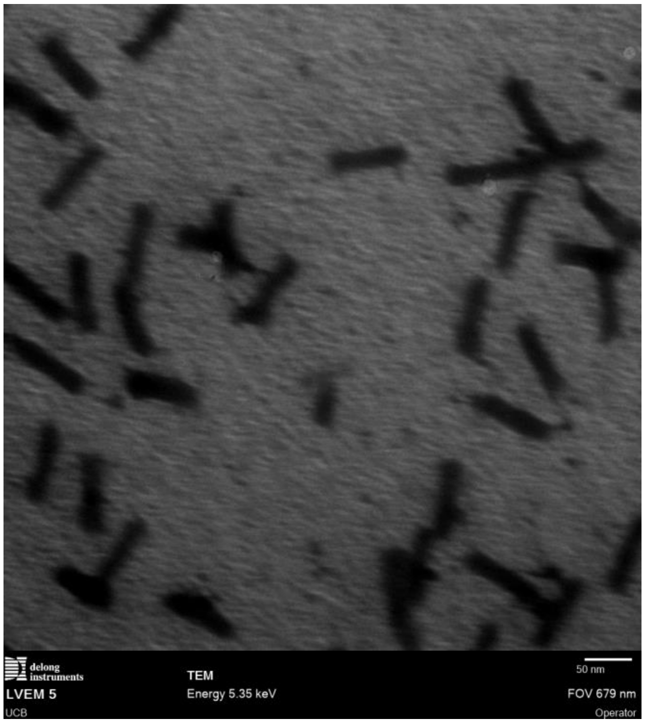


**Fig. S10.** TEM characterization of polymer coated 32HB (Rd@PL).

#### 7.2 Tube (Tu)

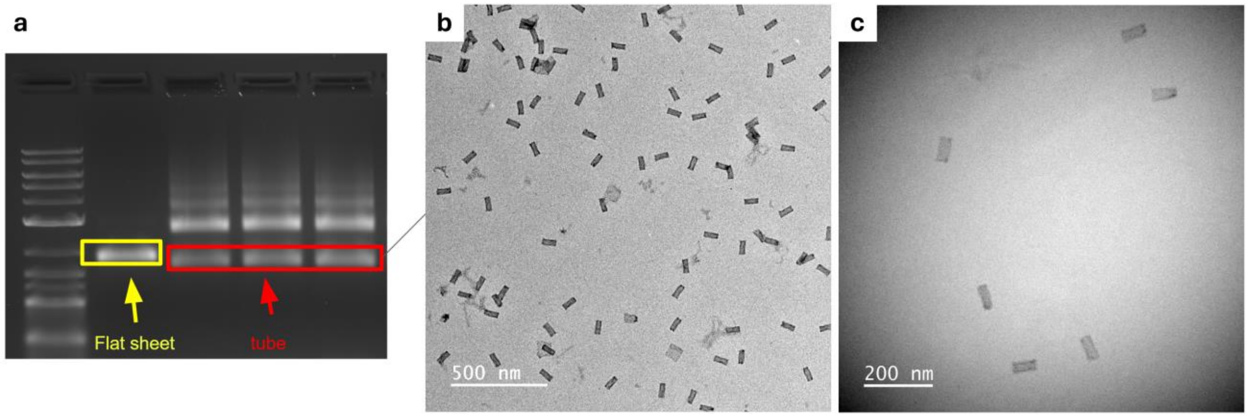


**Fig. S11.** Characterization of the tube (Tu). a) Agarose gel electrophoresis, (b, c) TEM of negatively stained Tu purified by gel electrophoresis.

#### 7.3 Box (Bx)

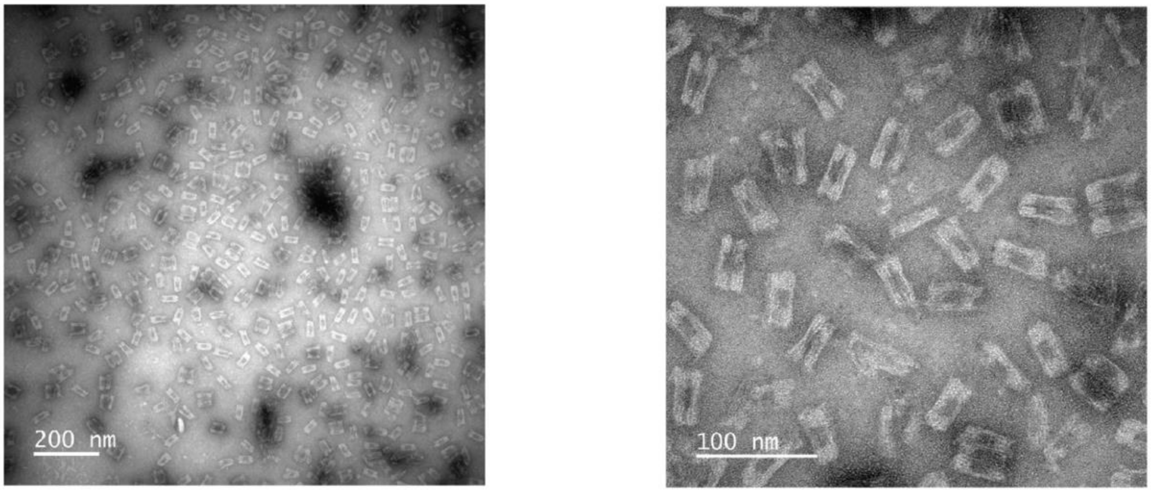


**Fig. S12.** Negatively-stained transmission electron microscopy of box (Bx).

#### 7.4 Tile (Sq) and aptamer modified tile (Sq-Apt)

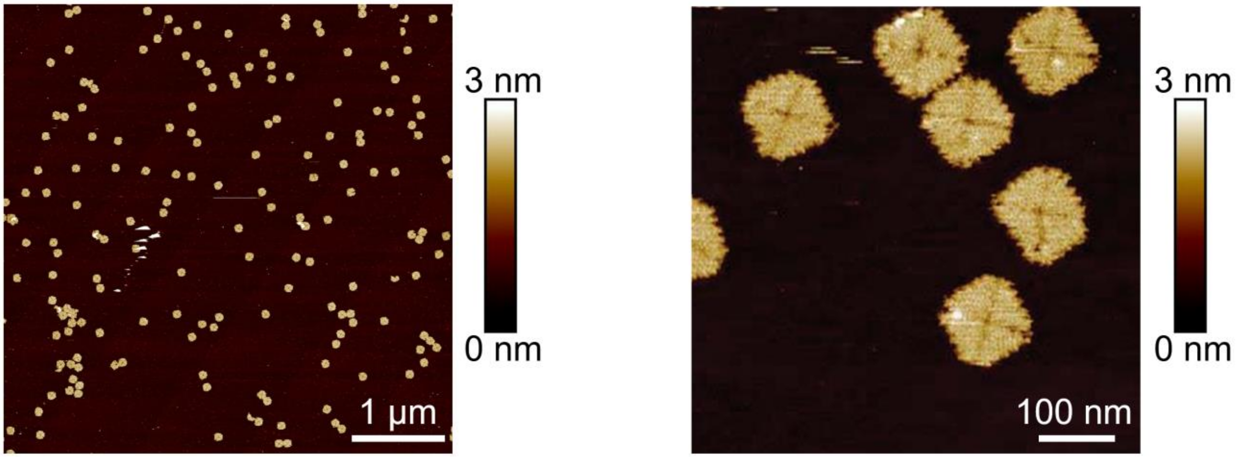


**Fig. S13.** Tapping Mode Atomic Force Microscopy of Square (Sq).

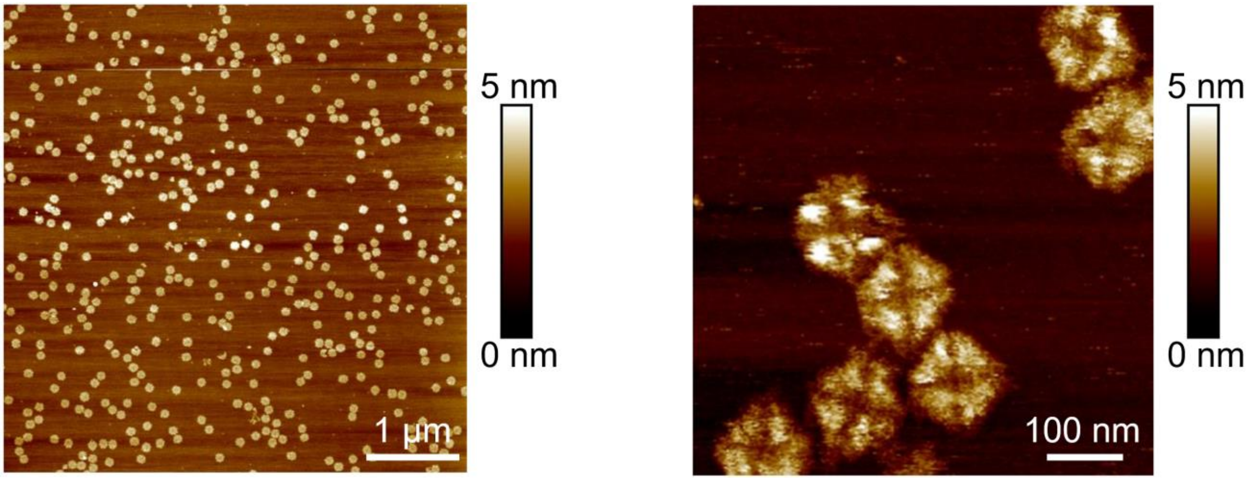


**Fig. S14.** Tapping Mode Atomic Force Microscopy of aptamer modified tile (Sq-Apt).

#### 7.5 Tetrahedron (Th) and cholesterol modified tetrahedron (Th-Ch)

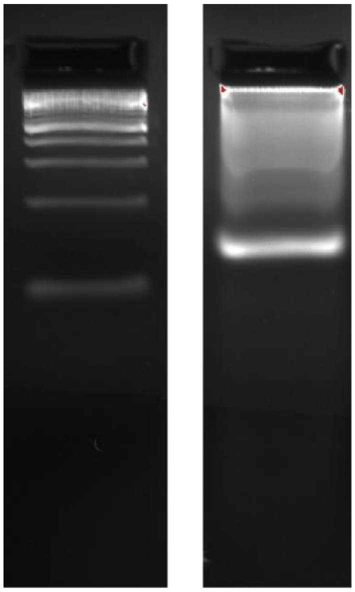

**Fig. S15.** Agarose gel characterization of tetrahedron with cholesterol (Th-Ch).

### 8 Nanostructure Designs

#### 8.1 Rod (Rd)

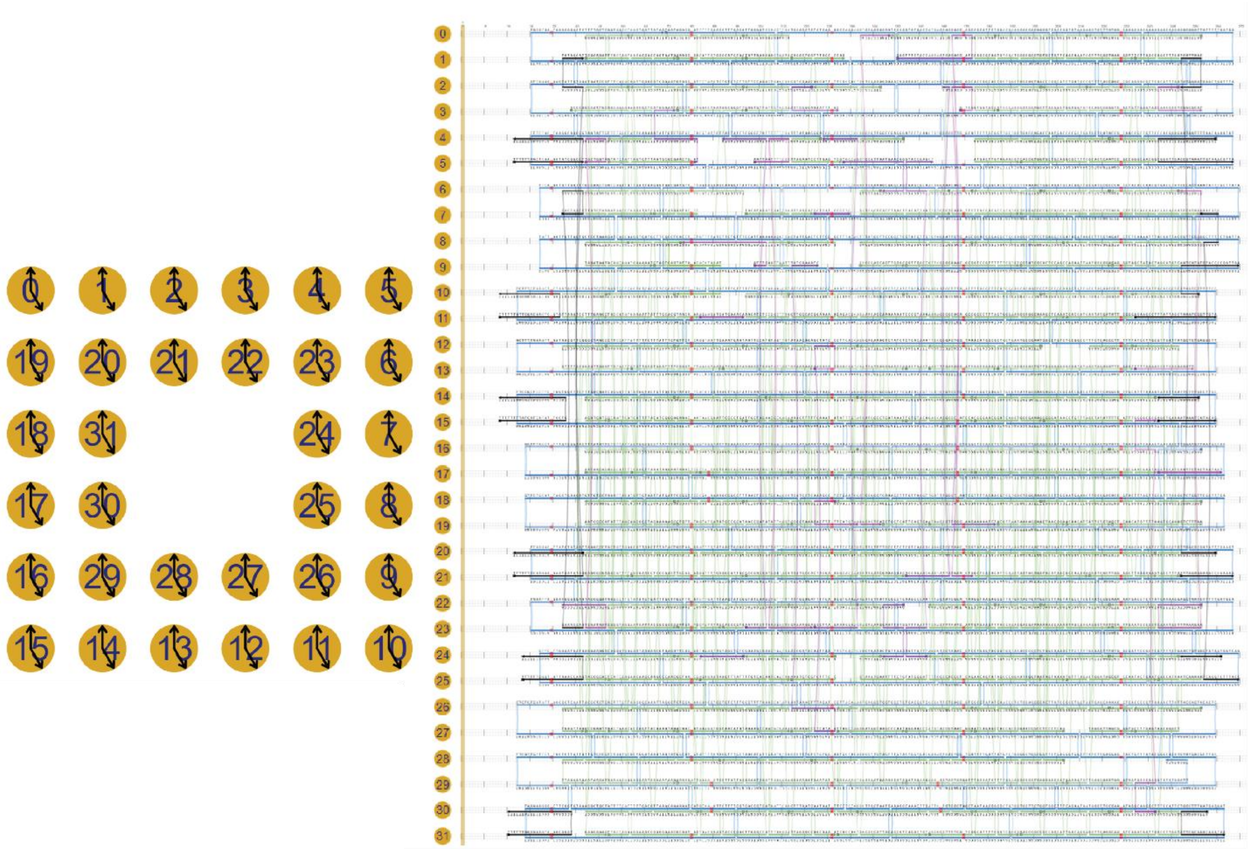

**Fig. S16.** Scadnano design of rod (Rd).

#### 8.2 Tube (Tu)

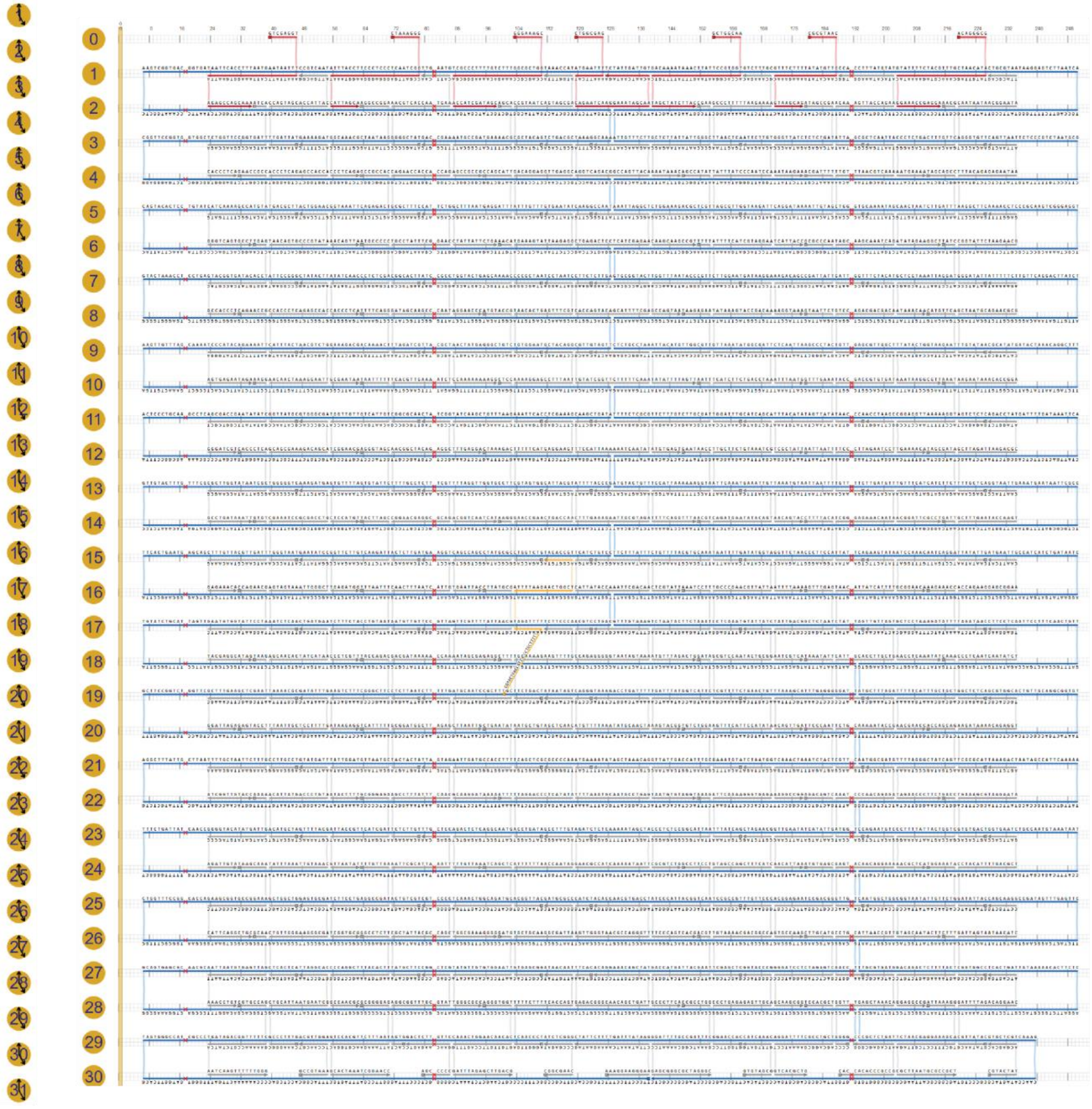

**Fig. S17.** Scadnano design of tube (Tu).

#### 8.3 Tile (Sq1)

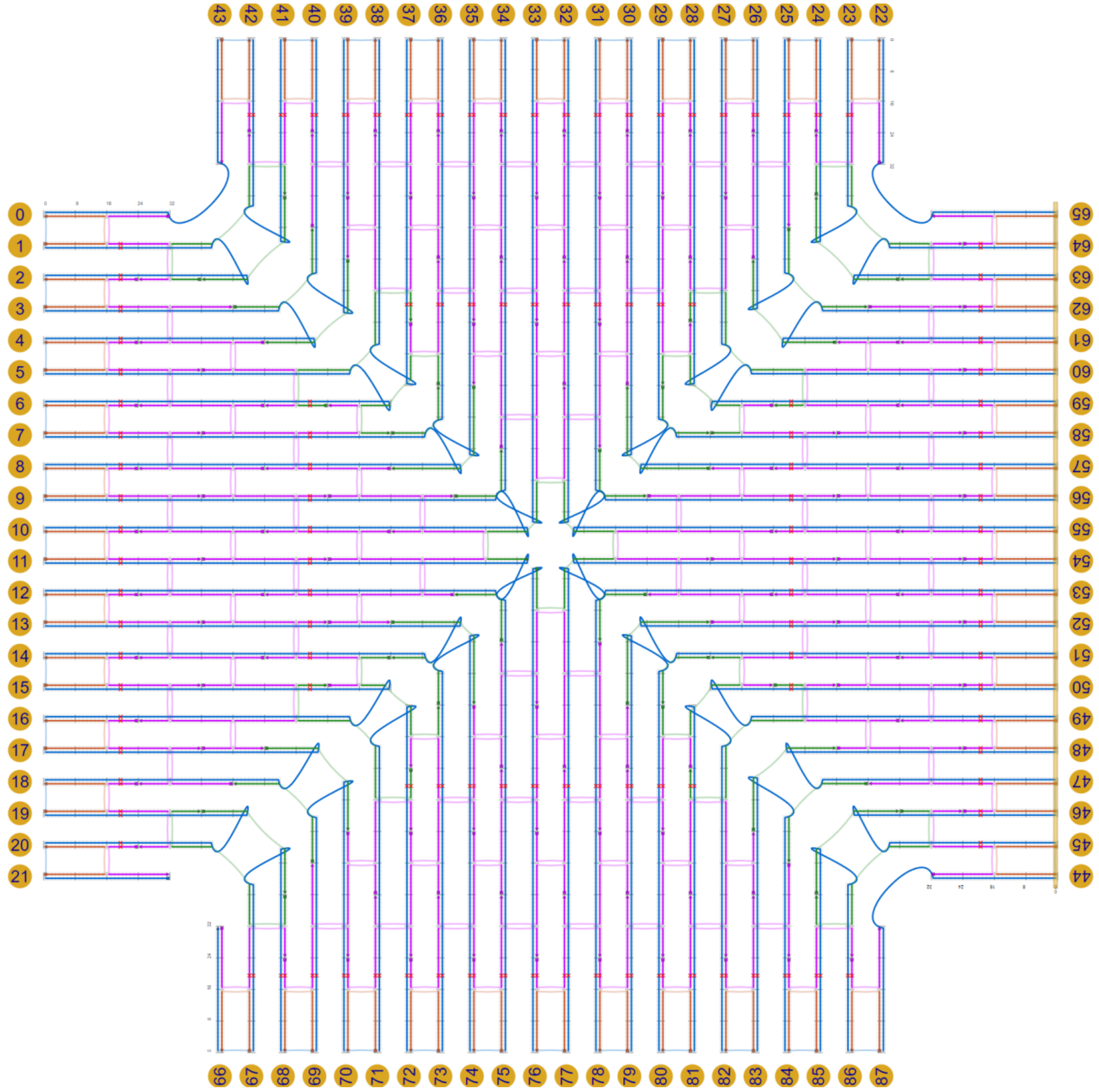


**Fig. S18.** Scadnano design of tile (Sq1).

### 9 Nanostructure Feature Quantifications

**Table S1.** The aspect ratio, surface area, $\zeta$ potential, number of nucleotides (NTs), 3D classification, vertices, cavities, and volume, as well as the presence/absence of aptamers, cholesterol moieties and Pll-g-Peg5K cationic polymer (PolyL), are listed for each of the nanostructures used in this study.

| | Bx@PL | Sq-Apt1 | Sq-Apt1@PL | Sq3 | Sq3@PL | Sq-Apt2 | Sq-Apt2@PL | Sq2 | Sq2@PL | Rd | Th | Th@PL | Th-Ch | Th-Ch@PL | Sq1 | Tu | Tu@PL |
|---|---|---|---|---|---|---|---|---|---|---|---|---|---|---|---|---|---|
| **Aspect Ratio** | 1.4 | 1.0 | 1.0 | 1.0 | 1.0 | 1.0 | 1.0 | 1.0 | 1.0 | 4.7 | 1.0 | 1.0 | 1.0 | 1.0 | 1.0 | 4.7 | 4.7 |
| **ZetaPotential** | -4.7 | -5.5 | -1.6 | -5.4 | -2.4 | -6.6 | -3.6 | -8.1 | -3.6 | -4.7 | -3.9 | -4.7 | -7.6 | 7.3 | -6.0 | -7.2 | -8.8 |
| **Scaffold Length** | 7249 | 7249 | 7249 | 7249 | 7249 | 7249 | 7249 | 7249 | 7249 | 7249 | 126 | 126 | 126 | 126 | 7249 | 7249 | 7249 |
| **Aptamers** | 0 | 1 | 1 | 0 | 0 | 1 | 1 | 0 | 0 | 0 | 0 | 0 | 0 | 0 | 0 | 0 | 0 |
| **3D** | 1 | 0 | 0 | 0 | 0 | 0 | 0 | 0 | 0 | 1 | 1 | 1 | 1 | 1 | 0 | 1 | 1 |
| **Vertices** | 12 | 4 | 4 | 4 | 4 | 4 | 4 | 4 | 4 | 8 | 4 | 4 | 4 | 4 | 4 | 0 | 0 |
| **Cavities** | 1 | 0 | 0 | 0 | 0 | 0 | 0 | 0 | 0 | 1 | 1 | 1 | 1 | 1 | 0 | 1 | 1 |
| **Cholesterol** | 0 | 0 | 0 | 0 | 0 | 0 | 0 | 0 | 0 | 0 | 0 | 0 | 1 | 1 | 0 | 0 | 0 |
| **PolyL** | 1 | 0 | 1 | 0 | 1 | 0 | 1 | 0 | 1 | 0 | 0 | 1 | 0 | 1 | 0 | 0 | 1 |
| **Nano Molecular Weight** | 4400000 | 4400000 | 4400000 | 4400000 | 4400000 | 4400000 | 4400000 | 4400000 | 4400000 | 4000000 | 58000 | 58000 | 58000 | 58000 | 4300000 | 4600000 | 4600000 |
| **Surface Area** | 7300 | 16000 | 16000 | 16000 | 16000 | 16000 | 16000 | 16000 | 16000 | 6900 | 80 | 80 | 80 | 80 | 16000 | 9800 | 9800 |
| **Volume** | 18000 | 15000 | 15000 | 15000 | 15000 | 15000 | 15000 | 15000 | 15000 | 17000 | 54 | 54 | 54 | 54 | 15000 | 9600 | 9600 |

### 10 Model Features and Importance Values

**Table S2.** All features used in XGBoost model (600+)

| Feature | Description | Group | Importance Presence | Importance Enriched |
|---|---|---|---|---|
| CK | Percent of lysine amino acids with predicted coil secondary structure in protein | Physiochemical | 0.04863844 | 0.05246249 |
| histone binding | GO:0042393 | Functional | 0.031012878 | 0.03200804 |
| IExpo | Percent of absolute solvent-accessible area in protein that is isoleucine absolute solvent-accessible area | Physiochemical | 0.022903422 | 0.012775368 |
| Sq8 | Percent of amino acids in protein with bend Q8 secondary structure prediction | Physiochemical | 0.021514 | 0.01068395 |
| ES | Percent of serine amino acids with predicted strand secondary structure in protein | Physiochemical | 0.0154957175 | 0.004306738 |

| | | | | |
|---|---|---|---|---|
| Q | Percent of amino acids in protein that are glutamine | Physiochemical | 0.013880412 | 0.0074803336 |
| HI | Percent of isoleucine amino acids with predicted helix secondary structure in protein | Physiochemical | 0.012711016 | 0.01107276 |
| Gq8 | Percent of amino acids in protein with $3_{10}$ helix Q8 secondary structure prediction | Physiochemical | 0.012531735 | 0.005741954 |
| K | Percent of amino acids in protein that are lysine | Physiochemical | 0.012482047 | 0.002750812 |
| psiMean | Mean ψ dihedral angle of amino acids in peptide | Physiochemical | 0.011956653 | 0.0069094133 |
| NExpo | Percent of absolute solvent-accessible area in protein that is asparagine absolute solvent-accessible area | Physiochemical | 0.011342108 | 0.007076583 |

| | | | | |
|---|---|---|---|---|
| HN | Percent of asparagine amino acids with predicted helix secondary structure in protein | Physiochemical | 0.01044615 | 0.0087263575 |
| Helix | Predicted fraction of peptide with helix secondary structure | Physiochemical | 0.0095185265 | 0.00808553 |
| KExpo | Percent of absolute solvent-accessible area in protein that is lysine absolute solvent-accessible area | Physiochemical | 0.009502832 | 0.0104895225 |
| MExpo | Percent of absolute solvent-accessible area in protein that is methionine absolute solvent-accessible area | Physiochemical | 0.0094225025 | 0.004207068 |
| HP | Percent of proline amino acids with predicted | Physiochemical | 0.008338282 | 0.014326307 |

| | | | | |
|---|---|---|---|---|
| | helix secondary structure in protein | | | |
| I | Percent of amino acids in protein that are isoleucine | Physiochemical | 0.008213813 | 0.012266831 |
| Cq8 | Percent of amino acids in protein with other Q8 secondary structure prediction | Physiochemical | 0.007888729 | 0.004797918 |
| calcium ion binding | GO:0005509 | Functional | 0.0076410547 | 0.00021841431 |
| EY | Percent of tyrosine amino acids with predicted strand secondary structure in protein | Physiochemical | 0.0075483657 | 0.0033847985 |
| HW | Percent of tryptophan amino acids with predicted helix secondary structure in protein | Physiochemical | 0.0075113866 | 0.012919128 |
| flexMed | Median flexibility calculated for 9 amino acid | Physiochemical | 0.0073936805 | 0.004015983 |

| | | | | |
|---|---|---|---|---|
| | windows in peptide | | | |
| CM | Percent of methionine amino acids with predicted coil secondary structure in protein | Physiochemical | 0.007382908 | 0.0046738 |
| YExpo | Percent of absolute solvent-accessible area in protein that is tyrosine absolute solvent-accessible area | Physiochemical | 0.007154082 | 0.007535076 |
| FExpo | Percent of absolute solvent-accessible area in protein that is phenylalanine absolute solvent-accessible area | Physiochemical | 0.007121844 | 0.007739568 |
| QExpo | Percent of absolute solvent-accessible area in protein that is glutamine | Physiochemical | 0.007101977 | 0.006185065 |

| | | | | |
|---|---|---|---|---|
| | absolute solvent-accessible area | | | |
| HC | Percent of cysteine amino acids with predicted helix secondary structure in protein | Physiochemical | 0.007087916 | 0.005808009 |
| HF | Percent of phenylalanine amino acids with predicted helix secondary structure in protein | Physiochemical | 0.0070673134 | 0.014266795 |
| CY | Percent of tyrosine amino acids with predicted coil secondary structure in protein | Physiochemical | 0.0070083593 | 0.008733654 |
| heparin binding | GO:0008201 | Functional | 0.0067929374 | 0.001117672 |
| TExpo | Percent of absolute solvent-accessible area in protein that is threonine absolute solvent- | Physiochemical | 0.0067129345 | 0.007986927 |

| | | | | |
|---|---|---|---|---|
| | accessible area | | | |
| A | Percent of amino acids in protein that are alanine | Physiochemical | 0.0066487417 | 0.0058263605 |
| EExpo | Percent of absolute solvent-accessible area in protein that is glutamic acid absolute solvent-accessible area | Physiochemical | 0.0064392528 | 0.003182299 |
| phiMean | Mean φ dihedral angle of amino acids in peptide | Physiochemical | 0.006375838 | 0.00709989 |
| F | Percent of amino acids in protein that are phenylalanine | Physiochemical | 0.006309852 | 0.004980066 |
| copper ion binding | GO:0005507 | Functional | 0.0062570167 | 0.0153401755 |
| W | Percent of amino acids in protein that are tryptophan | Physiochemical | 0.0060779406 | 0.007843743 |
| R | Percent of amino acids in protein that are arginine | Physiochemical | 0.0060498863 | 0.0027657521 |

| | | | | |
|---|---|---|---|---|
| P | Percent of amino acids in protein that are proline | Physiochemical | 0.0060089594 | 0.0047869226 |
| metal ion binding | GO:0046872 | Functional | 0.005885671 | 0.0029460369 |
| Eq8 | Percent of amino acids in protein with β-strand Q8 secondary structure prediction | Physiochemical | 0.005852295 | 0.015194813 |
| EN | Percent of asparagine amino acids with predicted strand secondary structure in protein | Physiochemical | 0.005838587 | 0.0069358936 |
| CA | Percent of alanine amino acids with predicted coil secondary structure in protein | Physiochemical | 0.0057569104 | 0.004113255 |
| Net Charge | Sum of amino acid charges in protein | Physiochemical | 0.0055615595 | 0.009622199 |
| Cholesterol | Presence or absence of cholesterol in DNA nanostructure | Nanostructure | 0.0054195034 | 0.0016413368 |
| L | Percent of amino acids | Physiochemical | 0.005250708 | 0.0036698366 |

| | | | | |
|---|---|---|---|---|
| | in protein that are leucine | | | |
| CE | Percent of glutamic acid amino acids with predicted coil secondary structure in protein | Physiochemical | 0.005236253 | 0.005656758 |
| CC | Percent of cysteine amino acids with predicted coil secondary structure in protein | Physiochemical | 0.0052308473 | 0.003072103 |
| V | Percent of amino acids in protein that are valine | Physiochemical | 0.00518711 | 0.0067176865 |
| structural molecule activity | GO:0005198 | Functional | 0.005160191 | 0.0005408098 |
| EL | Percent of leucine amino acids with predicted strand secondary structure in protein | Physiochemical | 0.0051426166 | 0.0038406798 |
| PExpo | Percent of absolute solvent-accessible area in protein that is proline absolute | Physiochemical | 0.005138945 | 0.003507204 |

| | | | | |
|---|---|---|---|---|
| | solvent-accessible area | | | |
| EP | Percent of proline amino acids with predicted strand secondary structure in protein | Physiochemical | 0.0051146746 | 0.004177352 |
| EE | Percent of glutamic acid amino acids with predicted strand secondary structure in protein | Physiochemical | 0.005107869 | 0.0040897774 |
| HExpo | Percent of absolute solvent-accessible area in protein that is histidine absolute solvent-accessible area | Physiochemical | 0.004991412 | 0.004982057 |
| Hydropathy | Mean hydropathy of amino acids in peptide | Physiochemical | 0.0049596876 | 0.0035770633 |
| T | Percent of amino acids in protein that are threonine | Physiochemical | 0.0048816865 | 0.0040888526 |

| | | | | |
|---|---|---|---|---|
| EC | Percent of cysteine amino acids with predicted strand secondary structure in protein | Physiochemical | 0.0048741046 | 0.0018868246 |
| VExpo | Percent of absolute solvent-accessible area in protein that is valine absolute solvent-accessible area | Physiochemical | 0.004840562 | 0.002865462 |
| LengthAminoAcids | Number of amino acids in peptide sequence | Physiochemical | 0.004830021 | 0.0044173226 |
| ED | Percent of aspartic acid amino acids with predicted strand secondary structure in protein | Physiochemical | 0.004807906 | 0.0023054387 |
| EK | Percent of lysine amino acids with predicted strand secondary structure in protein | Physiochemical | 0.004793387 | 0.003323209 |

| | | | | |
|---|---|---|---|---|
| Instability | Calculated instability index or protein | Physiochemical | 0.004777667 | 0.0031841216 |
| C | Percent of amino acids in protein that are cysteine | Physiochemical | 0.0047630165 | 0.0050633615 |
| HT | Percent of theronine amino acids with predicted helix secondary structure in protein | Physiochemical | 0.004730181 | 0.004387746 |
| ubiquitin protein ligase binding | GO:0031625 | Functional | 0.0047110817 | 0.0010748075 |
| DExpo | Percent of absolute solvent-accessible area in protein that is aspartic acid absolute solvent-accessible area | Physiochemical | 0.004598907 | 0.0037078927 |
| HA | Percent of alanine amino acids with predicted helix secondary structure in protein | Physiochemical | 0.004562256 | 0.0032787167 |
| SExpo | Percent of absolute | Physiochemical | 0.0045423526 | 0.0033802078 |

| | | | | |
|---|---|---|---|---|
| | solvent-accessible area in protein that is serine absolute solvent-accessible area | | | |
| CS | Percent of serine amino acids with predicted coil secondary structure in protein | Physiochemical | 0.004532428 | 0.0020374737 |
| flexMin | Minimum flexibility calculated for 9 amino acid windows in peptide | Physiochemical | 0.004424968 | 0.0045596007 |
| IsoelectricPoint | Calculated isoelectronic point of peptide | Physiochemical | 0.004421683 | 0.00599794 |
| E | Percent of amino acids in protein that are glutamic acid | Physiochemical | 0.004329267 | 0.0024576322 |
| LExpo | Percent of absolute solvent-accessible area in protein that is leucine absolute solvent- | Physiochemical | 0.0043197777 | 0.0077284477 |

| | | | | |
|---|---|---|---|---|
| | accessible area | | | |
| EW | Percent of tryptophan amino acids with predicted strand secondary structure in protein | Physiochemical | 0.0042427536 | 0.0025675804 |
| CExpo | Percent of absolute solvent-accessible area in protein that is cysteine absolute solvent-accessible area | Physiochemical | 0.004190235 | 0.0055555375 |
| HE | Percent of glutamic acid amino acids with predicted helix secondary structure in protein | Physiochemical | 0.0041807564 | 0.004397704 |
| HR | Percent of arginine amino acids with predicted helix secondary structure in protein | Physiochemical | 0.004146391 | 0.0027449953 |
| EA | Percent of alanine amino | Physiochemical | 0.0041345498 | 0.003537085 |

| | | | | |
|---|---|---|---|---|
| | acids with predicted strand secondary structure in protein | | | |
| Sheets | Predicted fraction of peptide with sheet secondary structure | Physiochemical | 0.004127359 | 0.004771958 |
| EV | Percent of valine amino acids with predicted strand secondary structure in protein | Physiochemical | 0.0040098606 | 0.0051077143 |
| ET | Percent of theronine amino acids with predicted strand secondary structure in protein | Physiochemical | 0.0040096007 | 0.0041241203 |
| CW | Percent of tryptophan amino acids with predicted coil secondary structure in protein | Physiochemical | 0.003929872 | 0.0041739736 |
| Y | Percent of amino acids | Physiochemical | 0.003915003 | 0.0032359073 |

| | | | | |
|---|---|---|---|---|
| | in protein that are tyrosine | | | |
| ATP binding | GO:0005524 | Functional | 0.0039109522 | 0.0012508489 |
| HQ | Percent of glutamine amino acids with predicted helix secondary structure in protein | Physiochemical | 0.0038139452 | 0.005136721 |
| CI | Percent of isoleucine amino acids with predicted coil secondary structure in protein | Physiochemical | 0.0037597474 | 0.004556303 |
| magnesium ion binding | GO:0000287 | Functional | 0.0037339032 | 0.0024344667 |
| CV | Percent of valine amino acids with predicted coil secondary structure in protein | Physiochemical | 0.0037236959 | 0.0025064582 |
| HH | Percent of histidine amino acids with predicted helix secondary structure in protein | Physiochemical | 0.0037198798 | 0.0070326347 |
| HY | Percent of tyrosine amino acids | Physiochemical | 0.003719863 | 0.0024474734 |

| | | | | |
|---|---|---|---|---|
| | with predicted helix secondary structure in protein | | | |
| Molecular Weight | Molecular weight of peptides | Physiochemical | 0.003705822 | 0.0030626669 |
| Bq8 | Percent of amino acids in protein with bridge Q8 secondary structure prediction | Physiochemical | 0.0036579077 | 0.0028045997 |
| HV | Percent of valine amino acids with predicted helix secondary structure in protein | Physiochemical | 0.0036504804 | 0.0029893762 |
| EG | Percent of glycine amino acids with predicted strand secondary structure in protein | Physiochemical | 0.003615857 | 0.0050780694 |
| RNA binding | GO:0003723 | Functional | 0.0035970677 | 0.0040025604 |
| H | Percent of amino acids in protein that are histidine | Physiochemical | 0.0035881614 | 0.0021263189 |
| CP | Percent of proline amino | Physiochemical | 0.0035748067 | 0.005206261 |

| | | | | |
|---|---|---|---|---|
| | acids with predicted coil secondary structure in protein | | | |
| WExpo | Percent of absolute solvent-accessible area in protein that is tryptophan absolute solvent-accessible area | Physiochemical | 0.003481327 | 0.0022490202 |
| GTPase binding | GO:0051020 | Functional | 0.0034726618 | 0.011554317 |
| GExpo | Percent of absolute solvent-accessible area in protein that is glycine absolute solvent-accessible area | Physiochemical | 0.0033750697 | 0.002746223 |
| CT | Percent of theronine amino acids with predicted coil secondary structure in protein | Physiochemical | 0.003330395 | 0.003598562 |
| PolyL | Presence or absence of polymer coating on | Nanostructure | 0.0032775742 | 0.0034173583 |

| | | | | |
|---|---|---|---|---|
| | DNA nanostructure | | | |
| actin filament binding | GO:0051015 | Functional | 0.0032676086 | 0.005472068 |
| M | Percent of amino acids in protein that are methionine | Physiochemical | 0.0032663608 | 0.004253273 |
| Turn | Predicted fraction of peptide with sheet secondary structure | Physiochemical | 0.0032624328 | 0.0031735497 |
| RExpo | Percent of absolute solvent-accessible area in protein that is arganine absolute solvent-accessible area | Physiochemical | 0.0032133479 | 0.0028540688 |
| identical protein binding | GO:0042802 | Functional | 0.0031653778 | 3.71E-05 |
| disMean | Mean disorder of amino acids in peptide | Physiochemical | 0.0031200403 | 0.0040420005 |
| ER | Percent of arginine amino acids with predicted strand secondary structure in protein | Physiochemical | 0.003119797 | 0.0028545223 |

| | | | | |
|---|---|---|---|---|
| EH | Percent of histidine amino acids with predicted strand secondary structure in protein | Physiochemical | 0.0030913164 | 0.0027681966 |
| CG | Percent of glycine amino acids with predicted coil secondary structure in protein | Physiochemical | 0.003077736 | 0.004392327 |
| HG | Percent of glycine amino acids with predicted helix secondary structure in protein | Physiochemical | 0.0030775182 | 0.005140609 |
| G | Percent of amino acids in protein that are glycine | Physiochemical | 0.0030686744 | 0.0017018807 |
| CL | Percent of leucine amino acids with predicted coil secondary structure in protein | Physiochemical | 0.003033426 | 0.007797996 |
| CD | Percent of aspartic acid amino acids with predicted coil | Physiochemical | 0.0030278647 | 0.0027521588 |

| | | | | |
|---|---|---|---|---|
| | secondary structure in protein | | | |
| D | Percent of amino acids in protein that are aspartic acid | Physiochemical | 0.003015637 | 0.0030553332 |
| AExpo | Percent of absolute solvent-accessible area in protein that is alanine absolute solvent-accessible area | Physiochemical | 0.0029496902 | 0.002806821 |
| CF | Percent of phenylalanine amino acids with predicted coil secondary structure in protein | Physiochemical | 0.002949603 | 0.0035011154 |
| EM | Percent of methionine amino acids with predicted strand secondary structure in protein | Physiochemical | 0.0029211408 | 0.0035284404 |
| CQ | Percent of glutamine amino acids with predicted coil | Physiochemical | 0.0028861326 | 0.0029729032 |

| | | | | |
|---|---|---|---|---|
| | secondary structure in protein | | | |
| N | Percent of amino acids in protein that are asparagine | Physiochemical | 0.002885919 | 0 |
| EF | Percent of phenylalanine amino acids with predicted strand secondary structure in protein | Physiochemical | 0.0028678286 | 0.0034034287 |
| EQ | Percent of glutamine amino acids with predicted strand secondary structure in protein | Physiochemical | 0.0028497777 | 0.0035517132 |
| HL | Percent of leucine amino acids with predicted helix secondary structure in protein | Physiochemical | 0.0027899044 | 0.0028721853 |
| CH | Percent of histidine amino acids with predicted coil secondary structure in protein | Physiochemical | 0.0027775997 | 0.004965474 |

| | | | | |
|---|---|---|---|---|
| EI | Percent of isoleucine amino acids with predicted strand secondary structure in protein | Physiochemical | 0.0027735587 | 0.0026387554 |
| Neutral Charge | Charge of protein at pH 7 | Physiochemical | 0.0027631475 | 0.002531418 |
| HM | Percent of methionine amino acids with predicted helix secondary structure in protein | Physiochemical | 0.0026881457 | 0.0027733883 |
| S | Percent of amino acids in protein that are serine | Physiochemical | 0.0026719181 | 0.002270447 |
| Tq8 | Percent of amino acids in protein with turn Q8 secondary structure prediction | Physiochemical | 0.002660281 | 0.0034040804 |
| Iq8 | Percent of amino acids in protein with π-helix Q8 secondary structure prediction | Physiochemical | 0.002650441 | 0.0024488708 |

| | | | | |
|---|---|---|---|---|
| Hq8 | Percent of amino acids in protein with α-helix Q8 secondary structure prediction | Physiochemical | 0.0026386296 | 0.0028556222 |
| HD | Percent of aspartic acid amino acids with predicted helix secondary structure in protein | Physiochemical | 0.0025976468 | 0.003042708 |
| CR | Percent of arginine amino acids with predicted coil secondary structure in protein | Physiochemical | 0.0025823761 | 0.003803373 |
| Surface Area | Estimated theoretical surface area of DNA nanostructure | Nanostructure | 0.0025509486 | 0.0018697886 |
| Aromaticity | Aromaticity of protein sequence | Physiochemical | 0.0025057846 | 0.004181103 |
| flexMax | Maximum flexibility calculated for 9 amino acid windows in peptide | Physiochemical | 0.0023994655 | 0.0045033926 |

| | | | | |
|---|---|---|---|---|
| CN | Percent of asparagine amino acids with predicted coil secondary structure in protein | Physiochemical | 0.002384828 | 0.0026873467 |
| lysine-acetylated histone binding | GO:0070577 | Functional | 0.0023365957 | 0.0010582681 |
| HK | Percent of lysine amino acids with predicted helix secondary structure in protein | Physiochemical | 0.0023337053 | 0.0033340827 |
| Vertices | Number of vertices in DNA nanostructure | Nanostructure | 0.002311988 | 0.0018900147 |
| Interactivity | Number of known binary protein-protein interactions | Functional | 0.0022950224 | 0.0024997261 |
| HS | Percent of serine amino acids with predicted helix secondary structure in protein | Physiochemical | 0.002294315 | 0.0031527379 |
| ZetaPotential | Measured or imputed zetapotential | Nanostructure | 0.0019191571 | 0.0014764096 |

| | | | | |
|---|---|---|---|---|
| | of DNA nanostructure | | | |
| Aspect Ratio | Estimated theoretical aspect ratio of DNA nanostructure | Nanostructure | 0.0018817135 | 0.0014926624 |
| phosphatidylserine binding | GO:0001786 | Functional | 0.0017924556 | 0.0015726469 |
| Aptamers | Presence or absence of aptamers strands in DNA nanostructure | Nanostructure | 0.0017034491 | 0.0012154155 |
| growth factor activity | GO:0008083 | Functional | 0.0016937145 | 0.0113259815 |
| ATP hydrolysis activity | GO:0016887 | Functional | 0.0015979435 | 0.00063684926 |
| 3D | Classification of nanostructure as 3-dimensional (1) or 2-dimensional (0) | Nanostructure | 0.001419479 | 0.0012264238 |
| tumor necrosis factor receptor binding | GO:0005164 | Functional | 0.001410404 | 0.00073657517 |
| cholesterol transfer activity | GO:0120020 | Functional | 0.0012966442 | 0 |
| Scaffold Length | Length in nucleotides of DNA scaffold. | Nanostructure | 0.0012578836 | 0.0015469233 |
| cysteine-type endopeptidase activity | GO:0004197 | Functional | 0.0012295041 | 0.0009149186 |
| Nano Molecular Weight | Estimated molecular weight of | Nanostructure | 0.0011459987 | 0.011681802 |

| | | | | |
|---|---|---|---|---|
| | DNA nanostructure | | | |
| antigen binding | GO:0003823 | Functional | 0.0010426592 | 0.00011268411 |
| Volume | Estimated theoretical Volume of DNA nanostructure | Nanostructure | 0.0010300435 | 0.001217449 |
| GDP binding | GO:0019003 | Functional | 0.0009774928 | 0.0006500999 |
| sarcosine dehydrogenase activity | GO:0008480 | Functional | 0.000966357 | 0.0008052063 |
| NADP binding | GO:0050661 | Functional | 0.00080285827 | 0 |
| structural constituent of chromatin | GO:0030527 | Functional | 0.00074735866 | 0 |
| endopeptidase inhibitor activity | GO:0004866 | Functional | 0.0007340728 | 0 |
| proteasome-activating activity | GO:0036402 | Functional | 0.00055007456 | 0 |
| protein homodimerization activity | GO:0042803 | Functional | 0.00043907773 | 0.0012528334 |
| misfolded protein binding | GO:0051787 | Functional | 0.00033630515 | 0 |
| virus receptor activity | GO:0001618 | Functional | 0.00016324261 | 0 |
| G protein activity | GO:0003925 | Functional | 4.10E-05 | 0.00029097393 |
| 1-alkyl-2-acetylglycerophosphocholine esterase activity | GO:0003847 | Functional | 0 | 0 |
| 1-phosphatidylinositol binding | GO:0005545 | Functional | 0 | 0 |
| 1-phosphatidylinositol-3-kinase activity | GO:0016303 | Functional | 0 | 0 |
| 1-phosphatidylinositol-4-phosphate 3-kinase activity | GO:0035005 | Functional | 0 | 0 |

| | | | | |
|---|---|---|---|---|
| 1-phosphatidylinositol-4,5-bisphosphate 3-kinase activity | GO:0046934 | Functional | 0 | 0 |
| 14-3-3 protein binding | GO:0071889 | Functional | 0 | 0 |
| 3 iron, 4 sulfur cluster binding | GO:0051538 | Functional | 0 | 0 |
| 4 iron, 4 sulfur cluster binding | GO:0051539 | Functional | 0 | 0 |
| 5'-nucleotidase activity | GO:0008253 | Functional | 0 | 0 |
| 5S rRNA binding | GO:0008097 | Functional | 0 | 0 |
| A1 adenosine receptor binding | GO:0031686 | Functional | 0 | 0 |
| ABC-type sterol transporter activity | GO:0034041 | Functional | 0 | 0 |
| ABC-type transporter activity | GO:0140359 | Functional | 0 | 0 |
| ABC-type xenobiotic transporter activity | GO:0008559 | Functional | 0 | 0 |
| acetoacetyl-CoA hydrolase activity | GO:0047603 | Functional | 0 | 0 |
| acetyl-CoA hydrolase activity | GO:0003986 | Functional | 0 | 0 |
| acetylcholine receptor activity | GO:0015464 | Functional | 0 | 0 |
| aconitate hydratase activity | GO:0003994 | Functional | 0 | 0 |
| acrosin binding | GO:0032190 | Functional | 0 | 0 |
| actin binding | GO:0003779 | Functional | 0 | 0 |
| actin monomer binding | GO:0003785 | Functional | 0 | 0 |
| acyl-CoA hydrolase activity | GO:0016289 | Functional | 0 | 0 |
| acyl-L-homoserine-lactone lactonohydrolase activity | GO:0102007 | Functional | 0 | 0 |
| adenosylhomocysteinase activity | GO:0004013 | Functional | 0 | 0 |
| adenyl-nucleotide exchange factor activity | GO:0000774 | Functional | 0 | 0 |
| adipokinetic hormone receptor activity | GO:0097003 | Functional | 0 | 0 |
| adiponectin binding | GO:0055100 | Functional | 0 | 0 |
| ADP binding | GO:0043531 | Functional | 0 | 0 |
| alpha-amylase activity | GO:0004556 | Functional | 0 | 0 |
| alpha-catenin binding | GO:0045294 | Functional | 0 | 0 |

| | | | | |
|---|---|---|---|---|
| alpha-tubulin binding | GO:0043014 | Functional | 0 | 0 |
| amino acid sensor activity | GO:0140785 | Functional | 0 | 0 |
| aminoacylase activity | GO:0004046 | Functional | 0 | 0 |
| aminopeptidase activity | GO:0004177 | Functional | 0 | 0 |
| amyloid-beta binding | GO:0001540 | Functional | 0 | 0 |
| angiostatin binding | GO:0043532 | Functional | 0 | 0 |
| ankyrin binding | GO:0030506 | Functional | 0 | 0 |
| ankyrin repeat binding | GO:0071532 | Functional | 0 | 0 |
| antioxidant activity | GO:0016209 | Functional | 0 | 0 |
| apolipoprotein A-I binding | GO:0034186 | Functional | 0 | 0 |
| apolipoprotein A-I receptor binding | GO:0034191 | Functional | 0 | 0 |
| apolipoprotein binding | GO:0034185 | Functional | 0 | 0 |
| apolipoprotein receptor binding | GO:0034190 | Functional | 0 | 0 |
| arachidonic acid binding | GO:0050544 | Functional | 0 | 0 |
| arginase activity | GO:0004053 | Functional | 0 | 0 |
| Arp2/3 complex binding | GO:0071933 | Functional | 0 | 0 |
| aryldialkylphosphatase activity | GO:0004063 | Functional | 0 | 0 |
| arylesterase activity | GO:0004064 | Functional | 0 | 0 |
| aspartic-type endopeptidase activity | GO:0004190 | Functional | 0 | 0 |
| aspartic-type peptidase activity | GO:0070001 | Functional | 0 | 0 |
| ATP-dependent activity, acting on RNA | GO:0008186 | Functional | 0 | 0 |
| ATP-dependent protein disaggregase activity | GO:0140545 | Functional | 0 | 0 |
| ATP-dependent protein folding chaperone | GO:0140662 | Functional | 0 | 0 |
| ATPase binding | GO:0051117 | Functional | 0 | 0 |
| ATPase-coupled intramembrane lipid transporter activity | GO:0140326 | Functional | 0 | 0 |

| | | | | |
|---|---|---|---|---|
| ATPase-coupled lipid transmembrane transporter activity | GO:0034040 | Functional | 0 | 0 |
| ATPase-coupled transmembrane transporter activity | GO:0042626 | Functional | 0 | 0 |
| axon guidance receptor activity | GO:0008046 | Functional | 0 | 0 |
| beta-catenin binding | GO:0008013 | Functional | 0 | 0 |
| bis(5'-adenosyl)-hexaphosphatase activity | GO:0034431 | Functional | 0 | 0 |
| bis(5'-adenosyl)-pentaphosphatase activity | GO:0034432 | Functional | 0 | 0 |
| bis(5'-nucleosyl)-tetraphosphatase (asymmetrical) activity | GO:0004081 | Functional | 0 | 0 |
| brain-derived neurotrophic factor binding | GO:0048403 | Functional | 0 | 0 |
| bubble DNA binding | GO:0000405 | Functional | 0 | 0 |
| C3HC4-type RING finger domain binding | GO:0055131 | Functional | 0 | 0 |
| C5L2 anaphylatoxin chemotactic receptor binding | GO:0031715 | Functional | 0 | 0 |
| cadherin binding | GO:0045296 | Functional | 0 | 0.00071312196 |
| calcidiol binding | GO:1902118 | Functional | 0 | 0 |
| calcium channel inhibitor activity | GO:0019855 | Functional | 0 | 0 |
| calcium oxalate binding | GO:0046904 | Functional | 0 | 0 |
| calcium-activated potassium channel activity | GO:0015269 | Functional | 0 | 0 |
| calcium-dependent cysteine-type endopeptidase activity | GO:0004198 | Functional | 0 | 0 |
| calcium-dependent phospholipid binding | GO:0005544 | Functional | 0 | 0 |
| calcium-dependent protein binding | GO:0048306 | Functional | 0 | 0 |

| | | | | |
|---|---|---|---|---|
| calmodulin binding | GO:0005516 | Functional | 0 | 0 |
| carbohydrate binding | GO:0030246 | Functional | 0 | 0 |
| carbohydrate derivative binding | GO:0097367 | Functional | 0 | 0 |
| carboxylic ester hydrolase activity | GO:0052689 | Functional | 0 | 0 |
| carboxypeptidase activity | GO:0004180 | Functional | 0 | 0 |
| catalase activity | GO:0004096 | Functional | 0 | 0 |
| Cavities | Number of cavities in DNA nanostructure | Nanostructure | 0 | 0 |
| cell adhesion molecule binding | GO:0050839 | Functional | 0 | 0 |
| cell adhesive protein binding involved in bundle of His cell-Purkinje myocyte communication | GO:0086083 | Functional | 0 | 0 |
| ceramide binding | GO:0097001 | Functional | 0 | 0 |
| ceramide transfer activity | GO:0120017 | Functional | 0 | 0 |
| cerebroside transfer activity | GO:0140340 | Functional | 0 | 0 |
| chemokine activity | GO:0008009 | Functional | 0 | 0 |
| chemorepellent activity | GO:0045499 | Functional | 0 | 0 |
| chloride channel activity | GO:0005254 | Functional | 0 | 0 |
| chloride ion binding | GO:0031404 | Functional | 0 | 0.003923789 |
| cholesterol binding | GO:0015485 | Functional | 0 | 0 |
| choline binding | GO:0033265 | Functional | 0 | 0 |
| choloyl-CoA hydrolase activity | GO:0033882 | Functional | 0 | 0 |
| chromatin binding | GO:0003682 | Functional | 0 | 0 |
| chromatin DNA binding | GO:0031490 | Functional | 0 | 0 |
| chromatin-protein adaptor activity | GO:0140463 | Functional | 0 | 0 |
| citrate (Si)-synthase activity | GO:0004108 | Functional | 0 | 0 |
| clathrin binding | GO:0030276 | Functional | 0 | 0 |
| clathrin heavy chain binding | GO:0032050 | Functional | 0 | 0 |

| | | | | |
|---|---|---|---|---|
| clathrin light chain binding | GO:0032051 | Functional | 0 | 0 |
| clathrin-uncoating ATPase activity | GO:1990833 | Functional | 0 | 0 |
| collagen binding | GO:0005518 | Functional | 0 | 0 |
| collagen V binding | GO:0070052 | Functional | 0 | 0 |
| complement binding | GO:0001848 | Functional | 0 | 0 |
| complement component C1q complex binding | GO:0001849 | Functional | 0 | 0 |
| complement component C3a binding | GO:0001850 | Functional | 0 | 0 |
| complement component C3b binding | GO:0001851 | Functional | 0 | 0 |
| coreceptor activity | GO:0015026 | Functional | 0 | 0 |
| CX | Percent of undertermined or atypical amino acids with predicted coil secondary structure in protein | Physiochemical | 0 | 0 |
| cyclosporin A binding | GO:0016018 | Functional | 0 | 0 |
| cysteine-type aminopeptidase activity | GO:0070005 | Functional | 0 | 0 |
| cysteine-type deubiquitinase activity | GO:0004843 | Functional | 0 | 0 |
| cysteine-type endopeptidase inhibitor activity | GO:0004869 | Functional | 0 | 0 |
| cysteine-type peptidase activity | GO:0008234 | Functional | 0 | 0 |
| cytoskeletal motor activity | GO:0003774 | Functional | 0 | 0 |
| cytoskeletal protein binding | GO:0008092 | Functional | 0 | 0 |
| cytoskeletal protein-membrane anchor activity | GO:0106006 | Functional | 0 | 0 |
| cytoskeleton-nuclear membrane anchor activity | GO:0140444 | Functional | 0 | 0 |

| | | | | |
|---|---|---|---|---|
| damaged DNA binding | GO:0003684 | Functional | 0 | 0 |
| death receptor agonist activity | GO:0038177 | Functional | 0 | 0 |
| death receptor binding | GO:0005123 | Functional | 0 | 0 |
| delta-catenin binding | GO:0070097 | Functional | 0 | 0 |
| denatured protein binding | GO:0031249 | Functional | 0 | 0 |
| deoxycytidyl transferase activity | GO:0017125 | Functional | 0 | 0 |
| diacylglyceride transfer activity | GO:0140337 | Functional | 0 | 0 |
| diacylglycerol binding | GO:0019992 | Functional | 0 | 0 |
| dinitrosyl-iron complex binding | GO:0035731 | Functional | 0 | 0 |
| dipeptidase activity | GO:0016805 | Functional | 0 | 0 |
| dipeptidyl-peptidase activity | GO:0008239 | Functional | 0 | 0 |
| diphosphoinositol-polyphosphate diphosphatase activity | GO:0008486 | Functional | 0 | 0 |
| disordered domain specific binding | GO:0097718 | Functional | 0 | 0 |
| disulfide oxidoreductase activity | GO:0015036 | Functional | 0 | 0 |
| DNA binding | GO:0003677 | Functional | 0 | 0 |
| DNA helicase activity | GO:0003678 | Functional | 0 | 0 |
| DNA replication origin binding | GO:0003688 | Functional | 0 | 0 |
| DNA-binding transcription activator activity, RNA polymerase II-specific | GO:0001228 | Functional | 0 | 0 |
| DNA-binding transcription factor activity | GO:0003700 | Functional | 0 | 0 |
| DNA-binding transcription factor activity, RNA polymerase II-specific | GO:0000981 | Functional | 0 | 0 |
| DNA-binding transcription factor binding | GO:0140297 | Functional | 0 | 0 |

| | | | | |
|---|---|---|---|---|
| DNA-binding transcription repressor activity, RNA polymerase II-specific | GO:0001227 | Functional | 0 | 0.0008062766 |
| DNA-directed DNA polymerase activity | GO:0003887 | Functional | 0 | 0 |
| double-stranded DNA binding | GO:0003690 | Functional | 0 | 0 |
| double-stranded methylated DNA binding | GO:0010385 | Functional | 0 | 0 |
| double-stranded RNA adenosine deaminase activity | GO:0003726 | Functional | 0 | 0 |
| double-stranded RNA binding | GO:0003725 | Functional | 0 | 0 |
| dynein complex binding | GO:0070840 | Functional | 0 | 0 |
| dynein intermediate chain binding | GO:0045505 | Functional | 0 | 0 |
| dynein light intermediate chain binding | GO:0051959 | Functional | 0 | 0 |
| efflux transmembrane transporter activity | GO:0015562 | Functional | 0 | 0 |
| endopeptidase activity | GO:0004175 | Functional | 0 | 0 |
| endopolyphosphatase activity | GO:0000298 | Functional | 0 | 0 |
| enterobactin binding | GO:1903981 | Functional | 0 | 0 |
| enzyme binding | GO:0019899 | Functional | 0 | 0 |
| enzyme inhibitor activity | GO:0004857 | Functional | 0 | 0 |
| enzyme regulator activity | GO:0030234 | Functional | 0 | 0 |
| ephrin receptor binding | GO:0046875 | Functional | 0 | 0 |
| EX | Percent of undertermined or atypical amino acids with predicted strand secondary structure in protein | Physiochemical | 0 | 0 |
| exogenous protein binding | GO:0140272 | Functional | 0 | 0 |

| | | | | |
|---|---|---|---|---|
| extracellular matrix binding | GO:0050840 | Functional | 0 | 0 |
| extracellular matrix structural constituent | GO:0005201 | Functional | 0 | 0 |
| extracellular matrix structural constituent conferring compression resistance | GO:0030021 | Functional | 0 | 0 |
| extracellular matrix structural constituent conferring tensile strength | GO:0030020 | Functional | 0 | 0 |
| FAD binding | GO:0071949 | Functional | 0 | 0 |
| fatty acid binding | GO:0005504 | Functional | 0 | 0 |
| fatty acyl-CoA hydrolase activity | GO:0047617 | Functional | 0 | 0 |
| Fc-gamma receptor I complex binding | GO:0034988 | Functional | 0 | 0 |
| ferric iron binding | GO:0008199 | Functional | 0 | 0 |
| ferrous iron binding | GO:0008198 | Functional | 0 | 0 |
| ferroxidase activity | GO:0004322 | Functional | 0 | 0 |
| fibrinogen binding | GO:0070051 | Functional | 0 | 0 |
| fibroblast growth factor binding | GO:0017134 | Functional | 0 | 0 |
| fibronectin binding | GO:0001968 | Functional | 0 | 0 |
| flavin-dependent sulfhydryl oxidase activity | GO:0016971 | Functional | 0 | 0 |
| four-way junction DNA binding | GO:0000400 | Functional | 0 | 0 |
| fructose binding | GO:0070061 | Functional | 0 | 0 |
| fructose-1-phosphate aldolase activity | GO:0061609 | Functional | 0 | 0 |
| fructose-bisphosphate aldolase activity | GO:0004332 | Functional | 0 | 0 |
| G protein-coupled receptor binding | GO:0001664 | Functional | 0 | 0 |
| gamma-catenin binding | GO:0045295 | Functional | 0 | 0 |

| | | | | |
|---|---|---|---|---|
| general transcription initiation factor binding | GO:0140296 | Functional | 0 | 0 |
| glucuronosyl-N-acetylgalactosaminyl-proteoglycan 4-beta-N-acetylgalactosaminyltransferase activity | GO:0047238 | Functional | 0 | 0 |
| glutamine synthetase activity | GO:0004356 | Functional | 0 | 0 |
| glutathione peroxidase activity | GO:0004602 | Functional | 0 | 0 |
| glutathione transferase activity | GO:0004364 | Functional | 0 | 0 |
| glyceraldehyde-3-phosphate dehydrogenase (NAD+) (phosphorylating) activity | GO:0004365 | Functional | 0 | 0 |
| glycine-tRNA ligase activity | GO:0004820 | Functional | 0 | 0 |
| glycosaminoglycan binding | GO:0005539 | Functional | 0 | 0 |
| Gravy | Calculated gravy score of peptide | Physiochemical | 0 | 0 |
| growth factor binding | GO:0019838 | Functional | 0 | 0 |
| GTP binding | GO:0005525 | Functional | 0 | 0.00089870155 |
| GTP-dependent protein binding | GO:0030742 | Functional | 0 | 0 |
| GTPase activating protein binding | GO:0032794 | Functional | 0 | 0.0027450728 |
| GTPase activator activity | GO:0005096 | Functional | 0 | 0 |
| GTPase activity | GO:0003924 | Functional | 0 | 0 |
| guanyl-nucleotide exchange factor activity | GO:0005085 | Functional | 0 | 0 |
| heat shock protein binding | GO:0031072 | Functional | 0 | 0 |
| helicase activity | GO:0004386 | Functional | 0 | 0 |
| heme binding | GO:0020037 | Functional | 0 | 0 |
| heme transmembrane transporter activity | GO:0015232 | Functional | 0 | 0 |
| hemoglobin alpha binding | GO:0031721 | Functional | 0 | 0 |

| | | | | |
|---|---|---|---|---|
| hemoglobin binding | GO:0030492 | Functional | 0 | 0 |
| heparan sulfate binding | GO:1904399 | Functional | 0 | 0 |
| heparan sulfate proteoglycan binding | GO:0043395 | Functional | 0 | 0 |
| high-density lipoprotein particle binding | GO:0008035 | Functional | 0 | 0 |
| high-density lipoprotein particle receptor binding | GO:0070653 | Functional | 0 | 0 |
| histone acetyltransferase activity | GO:0004402 | Functional | 0 | 0 |
| histone deacetylase binding | GO:0042826 | Functional | 0 | 0 |
| histone H3 acetyltransferase activity | GO:0010484 | Functional | 0 | 0 |
| histone H3K14 acetyltransferase activity | GO:0036408 | Functional | 0 | 0 |
| histone H3T11 kinase activity | GO:0035402 | Functional | 0 | 0 |
| histone pre-mRNA DCP binding | GO:0071208 | Functional | 0 | 0 |
| hormone activity | GO:0005179 | Functional | 0 | 0 |
| Hsp70 protein binding | GO:0030544 | Functional | 0 | 0 |
| hyaluronic acid binding | GO:0005540 | Functional | 0 | 0 |
| hydrolase activity | GO:0016787 | Functional | 0 | 0 |
| hydroxymethylglutaryl-CoA hydrolase activity | GO:0047994 | Functional | 0 | 0 |
| IgA binding | GO:0019862 | Functional | 0 | 0 |
| IgG binding | GO:0019864 | Functional | 0 | 0 |
| immunoglobulin binding | GO:0019865 | Functional | 0 | 0 |
| immunoglobulin receptor binding | GO:0034987 | Functional | 0 | 0.0020631605 |
| importin-alpha family protein binding | GO:0061676 | Functional | 0 | 0 |
| inositol-3,5-bisdiphosphate-2,3,4,6-tetrakisphosphate 5-diphosphatase activity | GO:0052848 | Functional | 0 | 0 |

| | | | | |
|---|---|---|---|---|
| inositol-5-diphosphate-1,2,3,4,6-pentakisphosphate diphosphatase activity | GO:0052845 | Functional | 0 | 0 |
| insulin-like growth factor binding | GO:0005520 | Functional | 0 | 0 |
| integrin binding | GO:0005178 | Functional | 0 | 0 |
| interleukin-1 binding | GO:0019966 | Functional | 0 | 0 |
| interleukin-2 receptor binding | GO:0005134 | Functional | 0 | 0 |
| interleukin-8 binding | GO:0019959 | Functional | 0 | 0 |
| intermediate filament binding | GO:0019215 | Functional | 0 | 0 |
| intraciliary transport particle A binding | GO:0120160 | Functional | 0 | 0 |
| ionotropic glutamate receptor binding | GO:0035255 | Functional | 0 | 0 |
| iron chaperone activity | GO:0034986 | Functional | 0 | 0 |
| iron ion binding | GO:0005506 | Functional | 0 | 0 |
| isocitrate dehydrogenase (NAD+) activity | GO:0004449 | Functional | 0 | 0 |
| isocitrate dehydrogenase (NADP+) activity | GO:0004450 | Functional | 0 | 0 |
| isomerase activity | GO:0016853 | Functional | 0 | 0 |
| JUN kinase binding | GO:0008432 | Functional | 0 | 0 |
| keratin filament binding | GO:1990254 | Functional | 0 | 0 |
| kinase activity | GO:0016301 | Functional | 0 | 0 |
| kinase binding | GO:0019900 | Functional | 0 | 0 |
| kinase inhibitor activity | GO:0019210 | Functional | 0 | 0 |
| kinase regulator activity | GO:0019207 | Functional | 0 | 0 |
| kinesin binding | GO:0019894 | Functional | 0 | 0 |
| kynurenine-oxoglutarate transaminase activity | GO:0016212 | Functional | 0 | 0 |
| L-aspartate:2-oxoglutarate aminotransferase activity | GO:0004069 | Functional | 0 | 0 |
| L-cysteine transaminase activity | GO:0047801 | Functional | 0 | 0 |

| | | | | |
|---|---|---|---|---|
| L-lactate dehydrogenase activity | GO:0004459 | Functional | 0 | 0 |
| lamin binding | GO:0005521 | Functional | 0 | 0 |
| laminin binding | GO:0043236 | Functional | 0 | 0 |
| laminin receptor activity | GO:0005055 | Functional | 0 | 0 |
| ligand-gated sodium channel activity | GO:0015280 | Functional | 0 | 0 |
| LIM domain binding | GO:0030274 | Functional | 0 | 0 |
| lipase binding | GO:0035473 | Functional | 0 | 0 |
| lipase inhibitor activity | GO:0055102 | Functional | 0 | 0 |
| lipid binding | GO:0008289 | Functional | 0 | 0 |
| lipid transporter activity | GO:0005319 | Functional | 0 | 0 |
| lipopeptide binding | GO:0071723 | Functional | 0 | 0 |
| lipopolysaccharide binding | GO:0001530 | Functional | 0 | 0.00019818774 |
| lipoprotein lipase activator activity | GO:0060230 | Functional | 0 | 0 |
| lipoprotein particle binding | GO:0071813 | Functional | 0 | 0 |
| lipoteichoic acid binding | GO:0070891 | Functional | 0 | 0 |
| lncRNA binding | GO:0106222 | Functional | 0 | 0 |
| long-chain fatty acyl-CoA hydrolase activity | GO:0052816 | Functional | 0 | 0 |
| low-density lipoprotein particle binding | GO:0030169 | Functional | 0 | 0 |
| low-density lipoprotein particle receptor binding | GO:0050750 | Functional | 0 | 0.009353189 |
| lysozyme activity | GO:0003796 | Functional | 0 | 0 |
| manganese ion binding | GO:0030145 | Functional | 0 | 0 |
| medium-chain fatty acyl-CoA hydrolase activity | GO:0052815 | Functional | 0 | 0 |
| metal chelating activity | GO:0046911 | Functional | 0 | 0 |
| metallocarboxypeptidase activity | GO:0004181 | Functional | 0 | 0 |
| metallodipeptidase activity | GO:0070573 | Functional | 0 | 0 |

| | | | | |
|---|---|---|---|---|
| metalloendopeptidase inhibitor activity | GO:0008191 | Functional | 0 | 0 |
| methionine adenosyltransferase activity | GO:0004478 | Functional | 0 | 0 |
| methylated histone binding | GO:0035064 | Functional | 0 | 0 |
| methylglyoxal synthase activity | GO:0008929 | Functional | 0 | 0 |
| MHC class I protein binding | GO:0042288 | Functional | 0 | 0 |
| MHC class II protein complex binding | GO:0023026 | Functional | 0 | 0 |
| microfilament motor activity | GO:0000146 | Functional | 0 | 0 |
| microtubule binding | GO:0008017 | Functional | 0 | 0 |
| microtubule motor activity | GO:0003777 | Functional | 0 | 0 |
| microtubule plus-end binding | GO:0051010 | Functional | 0 | 0 |
| minus-end-directed microtubule motor activity | GO:0008569 | Functional | 0 | 0 |
| miRNA binding | GO:0035198 | Functional | 0 | 0 |
| mitogen-activated protein kinase binding | GO:0051019 | Functional | 0 | 0 |
| molecular adaptor activity | GO:0060090 | Functional | 0 | 0 |
| molecular condensate scaffold activity | GO:0140693 | Functional | 0 | 0 |
| molecular function activator activity | GO:0140677 | Functional | 0 | 0 |
| molecular sequestering activity | GO:0140313 | Functional | 0 | 0 |
| monoatomic anion channel activity | GO:0005253 | Functional | 0 | 0.0011389724 |
| monoatomic cation channel activity | GO:0005261 | Functional | 0 | 0 |
| monoatomic ion channel activity | GO:0005216 | Functional | 0 | 0 |
| monosaccharide binding | GO:0048029 | Functional | 0 | 0 |
| mRNA 3'-UTR AU-rich region binding | GO:0035925 | Functional | 0 | 0 |
| mRNA 3'-UTR binding | GO:0003730 | Functional | 0 | 0 |

| | | | | |
|---|---|---|---|---|
| mRNA binding | GO:0003729 | Functional | 0 | 0 |
| myosin II binding | GO:0045159 | Functional | 0 | 0 |
| myosin phosphatase activity | GO:0017018 | Functional | 0 | 0 |
| myosin phosphatase regulator activity | GO:0017020 | Functional | 0 | 0 |
| N-acetylgalactosaminyl-proteoglycan 3-beta-glucuronosyltransferase activity | GO:0050510 | Functional | 0 | 0.006708251 |
| N-acetylmuramoyl-L-alanine amidase activity | GO:0008745 | Functional | 0 | 0 |
| N6-methyladenosine-containing RNA reader activity | GO:1990247 | Functional | 0 | 0 |
| NAD binding | GO:0051287 | Functional | 0 | 0 |
| NAD+ nucleosidase activity | GO:0003953 | Functional | 0 | 0 |
| NAD+ nucleotidase, cyclic ADP-ribose generating | GO:0061809 | Functional | 0 | 0 |
| nerve growth factor binding | GO:0048406 | Functional | 0 | 0 |
| neuropilin binding | GO:0038191 | Functional | 0 | 0 |
| NF-kappaB binding | GO:0051059 | Functional | 0 | 0 |
| nicotinate-nucleotide adenylyltransferase activity | GO:0004515 | Functional | 0 | 0 |
| nitric oxide binding | GO:0070026 | Functional | 0 | 0 |
| nitric-oxide synthase binding | GO:0050998 | Functional | 0 | 0 |
| non-membrane spanning protein tyrosine kinase activity | GO:0004715 | Functional | 0 | 0 |
| nuclear receptor binding | GO:0016922 | Functional | 0 | 0 |
| nucleosomal DNA binding | GO:0031492 | Functional | 0 | 0 |
| nucleotide binding | GO:0000166 | Functional | 0 | 0 |
| organic acid binding | GO:0043177 | Functional | 0 | 0 |
| oxidoreductase activity | GO:0016491 | Functional | 0 | 0 |
| oxidoreductase activity, acting on metal ions, oxygen as acceptor | GO:0016724 | Functional | 0 | 0 |

| | | | | |
|---|---|---|---|---|
| oxidoreductase activity, acting on peroxide as acceptor | GO:0016684 | Functional | 0 | 0 |
| oxygen binding | GO:0019825 | Functional | 0 | 0 |
| oxygen carrier activity | GO:0005344 | Functional | 0 | 0 |
| p53 binding | GO:0002039 | Functional | 0 | 0 |
| pattern recognition receptor activity | GO:0038187 | Functional | 0 | 0 |
| PDZ domain binding | GO:0030165 | Functional | 0 | 0 |
| peptidase activator activity | GO:0016504 | Functional | 0 | 0 |
| peptidase activator activity involved in apoptotic process | GO:0016505 | Functional | 0 | 0 |
| peptidase activity | GO:0008233 | Functional | 0 | 0 |
| peptide binding | GO:0042277 | Functional | 0 | 0 |
| peptide-lysine-N-acetyltransferase activity | GO:0061733 | Functional | 0 | 0 |
| peptidoglycan binding | GO:0042834 | Functional | 0 | 0 |
| peptidoglycan immune receptor activity | GO:0016019 | Functional | 0 | 0 |
| peptidyl-prolyl cis-trans isomerase activity | GO:0003755 | Functional | 0 | 0 |
| peroxidase activity | GO:0004601 | Functional | 0 | 0 |
| phosphatase binding | GO:0019902 | Functional | 0 | 0 |
| phosphate transmembrane transporter activity | GO:0005315 | Functional | 0 | 0 |
| phosphate:proton symporter activity | GO:0015317 | Functional | 0 | 0 |
| phosphatidic acid binding | GO:0070300 | Functional | 0 | 0 |
| phosphatidic acid transfer activity | GO:1990050 | Functional | 0 | 0 |
| phosphatidylcholine binding | GO:0031210 | Functional | 0 | 0 |
| phosphatidylcholine floppase activity | GO:0090554 | Functional | 0 | 0 |
| phosphatidylcholine transfer activity | GO:0120019 | Functional | 0 | 0 |

| | | | | |
|---|---|---|---|---|
| phosphatidylcholine-sterol O-acyltransferase activator activity | GO:0060228 | Functional | 0 | 0 |
| phosphatidylcholine-sterol O-acyltransferase activity | GO:0004607 | Functional | 0 | 0 |
| phosphatidylethanolamine binding | GO:0008429 | Functional | 0 | 0 |
| phosphatidylethanolamine transfer activity | GO:1904121 | Functional | 0 | 0 |
| phosphatidylglycerol binding | GO:1901611 | Functional | 0 | 0 |
| phosphatidylglycerol transfer activity | GO:0140339 | Functional | 0 | 0 |
| phosphatidylinositol 3-kinase catalytic subunit binding | GO:0036313 | Functional | 0 | 0 |
| phosphatidylinositol binding | GO:0035091 | Functional | 0 | 0 |
| phosphatidylinositol transfer activity | GO:0008526 | Functional | 0 | 0 |
| phosphatidylinositol-3-phosphate binding | GO:0032266 | Functional | 0 | 0 |
| phosphatidylinositol-3,4,5-trisphosphate binding | GO:0005547 | Functional | 0 | 0 |
| phosphatidylinositol-4,5-bisphosphate binding | GO:0005546 | Functional | 0 | 0 |
| phosphatidylserine decarboxylase activity | GO:0004609 | Functional | 0 | 0 |
| phosphoglycerate kinase activity | GO:0004618 | Functional | 0 | 0 |
| phospholipase activator activity | GO:0016004 | Functional | 0 | 0 |
| phospholipase binding | GO:0043274 | Functional | 0 | 0 |
| phospholipase C activity | GO:0004629 | Functional | 0 | 0 |
| phospholipase inhibitor activity | GO:0004859 | Functional | 0 | 0 |
| phospholipid binding | GO:0005543 | Functional | 0 | 0 |
| phospholipid transfer activity | GO:0120014 | Functional | 0 | 0 |

| | | | | |
|---|---|---|---|---|
| phospholipid transporter activity | GO:0005548 | Functional | 0 | 0 |
| phospholipid-hydroperoxide glutathione peroxidase activity | GO:0047066 | Functional | 0 | 0 |
| phosphopyruvate hydratase activity | GO:0004634 | Functional | 0 | 0 |
| phosphoserine residue binding | GO:0050815 | Functional | 0 | 0 |
| phosphotyrosine residue binding | GO:0001784 | Functional | 0 | 0 |
| platelet-activating factor acetyltransferase activity | GO:0047179 | Functional | 0 | 0 |
| poly(A) binding | GO:0008143 | Functional | 0 | 0 |
| poly(U) RNA binding | GO:0008266 | Functional | 0 | 0 |
| polysaccharide binding | GO:0030247 | Functional | 0 | 0 |
| potassium ion binding | GO:0030955 | Functional | 0 | 0 |
| pre-mRNA intronic binding | GO:0097157 | Functional | 0 | 0 |
| preribosome binding | GO:1990275 | Functional | 0 | 0 |
| primary miRNA binding | GO:0070878 | Functional | 0 | 0 |
| profilin binding | GO:0005522 | Functional | 0 | 0 |
| proline-rich region binding | GO:0070064 | Functional | 0 | 0 |
| propionyl-CoA carboxylase activity | GO:0004658 | Functional | 0 | 0 |
| prostaglandin binding | GO:1904593 | Functional | 0 | 0 |
| protease binding | GO:0002020 | Functional | 0 | 0 |
| protein carrier chaperone | GO:0140597 | Functional | 0 | 0 |
| protein dimerization activity | GO:0046983 | Functional | 0 | 0 |
| protein disulfide isomerase activity | GO:0003756 | Functional | 0 | 0 |
| protein domain specific binding | GO:0019904 | Functional | 0 | 0 |
| protein folding chaperone | GO:0044183 | Functional | 0 | 0 |
| protein heterodimerization activity | GO:0046982 | Functional | 0 | 0 |
| protein kinase A binding | GO:0051018 | Functional | 0 | 0 |

| | | | | |
|---|---|---|---|---|
| protein kinase A catalytic subunit binding | GO:0034236 | Functional | 0 | 0 |
| protein kinase A regulatory subunit binding | GO:0034237 | Functional | 0 | 0 |
| protein kinase activator activity | GO:0030295 | Functional | 0 | 0 |
| protein kinase activity | GO:0004672 | Functional | 0 | 0 |
| protein kinase binding | GO:0019901 | Functional | 0 | 0.024595255 |
| protein kinase C binding | GO:0005080 | Functional | 0 | 0 |
| protein kinase C inhibitor activity | GO:0008426 | Functional | 0 | 0 |
| protein phosphatase binding | GO:0019903 | Functional | 0 | 0 |
| protein phosphatase regulator activity | GO:0019888 | Functional | 0 | 0 |
| protein sequestering activity | GO:0140311 | Functional | 0 | 0 |
| protein serine kinase activity | GO:0106310 | Functional | 0 | 0 |
| protein serine/threonine kinase activity | GO:0004674 | Functional | 0 | 0 |
| protein serine/threonine phosphatase activity | GO:0004722 | Functional | 0 | 0 |
| protein tag activity | GO:0031386 | Functional | 0 | 0 |
| protein transmembrane transporter activity | GO:0008320 | Functional | 0 | 0 |
| protein tyrosine kinase activity | GO:0004713 | Functional | 0 | 0 |
| protein tyrosine phosphatase activity | GO:0004725 | Functional | 0 | 0 |
| protein-containing complex binding | GO:0044877 | Functional | 0 | 0 |
| protein-cysteine S-palmitoyltransferase activity | GO:0019706 | Functional | 0 | 0 |
| protein-disulfide reductase (NAD(P)H) activity | GO:0047134 | Functional | 0 | 0 |
| protein-disulfide reductase activity | GO:0015035 | Functional | 0 | 0 |
| protein-folding chaperone binding | GO:0051087 | Functional | 0 | 0 |

| | | | | |
|---|---|---|---|---|
| protein-glutamine gamma-glutamyltransferase activity | GO:0003810 | Functional | 0 | 0 |
| protein-macromolecule adaptor activity | GO:0030674 | Functional | 0 | 0 |
| protein-membrane adaptor activity | GO:0043495 | Functional | 0 | 0 |
| protein-RNA adaptor activity | GO:0140517 | Functional | 0 | 0 |
| protein-RNA sequence-specific adaptor activity | GO:0160134 | Functional | 0 | 0 |
| proteinase activated receptor binding | GO:0031871 | Functional | 0 | 0 |
| proteoglycan binding | GO:0043394 | Functional | 0 | 0 |
| proton transmembrane transporter activity | GO:0015078 | Functional | 0 | 0 |
| proton-transporting ATP synthase activity, rotational mechanism | GO:0046933 | Functional | 0 | 0 |
| proton-transporting ATPase activity, rotational mechanism | GO:0046961 | Functional | 0 | 0 |
| purine ribonucleoside triphosphate binding | GO:0035639 | Functional | 0 | 0 |
| pyridoxal phosphate binding | GO:0030170 | Functional | 0 | 0 |
| pyruvate kinase activity | GO:0004743 | Functional | 0 | 0 |
| RAGE receptor binding | GO:0050786 | Functional | 0 | 0 |
| receptor ligand activity | GO:0048018 | Functional | 0 | 0 |
| retinal binding | GO:0016918 | Functional | 0 | 0 |
| retinoic acid binding | GO:0001972 | Functional | 0 | 0 |
| retinol binding | GO:0019841 | Functional | 0 | 0 |
| retinol transmembrane transporter activity | GO:0034632 | Functional | 0 | 0 |
| retromer complex binding | GO:1905394 | Functional | 0 | 0 |
| ribosome binding | GO:0043022 | Functional | 0 | 0 |
| RNA cap binding | GO:0000339 | Functional | 0 | 0 |
| RNA helicase activity | GO:0003724 | Functional | 0 | 0 |

| | | | | |
|---|---|---|---|---|
| RNA nuclease activity | GO:0004540 | Functional | 0 | 0 |
| RNA polymerase II cis-regulatory region sequence-specific DNA binding | GO:0000978 | Functional | 0 | 0 |
| RNA polymerase II core promoter sequence-specific DNA binding | GO:0000979 | Functional | 0 | 0 |
| RNA polymerase II transcription regulatory region sequence-specific DNA binding | GO:0000977 | Functional | 0 | 0 |
| RNA polymerase II-specific DNA-binding transcription factor binding | GO:0061629 | Functional | 0 | 0 |
| S-nitrosoglutathione binding | GO:0035730 | Functional | 0 | 0 |
| S100 protein binding | GO:0044548 | Functional | 0 | 0 |
| scaffold protein binding | GO:0097110 | Functional | 0 | 0 |
| scavenger receptor activity | GO:0005044 | Functional | 0 | 0 |
| selenium binding | GO:0008430 | Functional | 0 | 0 |
| sequence-specific double-stranded DNA binding | GO:1990837 | Functional | 0 | 0 |
| serine-type endopeptidase activity | GO:0004252 | Functional | 0 | 0 |
| serine-type endopeptidase inhibitor activity | GO:0004867 | Functional | 0 | 0 |
| serine-type peptidase activity | GO:0008236 | Functional | 0 | 0 |
| SH2 domain binding | GO:0042169 | Functional | 0 | 0 |
| SH3 domain binding | GO:0017124 | Functional | 0 | 0.0004519035 |
| signaling receptor activity | GO:0038023 | Functional | 0 | 0 |
| signaling receptor binding | GO:0005102 | Functional | 0 | 0.0002532568 |
| single-stranded DNA binding | GO:0003697 | Functional | 0 | 0 |
| small GTPase binding | GO:0031267 | Functional | 0 | 0 |
| small molecule binding | GO:0036094 | Functional | 0 | 0 |
| SNARE binding | GO:0000149 | Functional | 0 | 0 |
| snoRNA binding | GO:0030515 | Functional | 0 | 0 |

| | | | | |
|---|---|---|---|---|
| somatostatin receptor binding | GO:0031877 | Functional | 0 | 0 |
| sphingomyelin transfer activity | GO:0140338 | Functional | 0 | 0 |
| STAT family protein binding | GO:0097677 | Functional | 0 | 0 |
| steroid binding | GO:0005496 | Functional | 0 | 0 |
| sterol esterase activity | GO:0004771 | Functional | 0 | 0 |
| structural constituent of cytoskeleton | GO:0005200 | Functional | 0 | 0 |
| structural constituent of eye lens | GO:0005212 | Functional | 0 | 0 |
| structural constituent of muscle | GO:0008307 | Functional | 0 | 0 |
| structural constituent of nuclear pore | GO:0017056 | Functional | 0 | 0 |
| structural constituent of postsynaptic actin cytoskeleton | GO:0098973 | Functional | 0 | 0 |
| structural constituent of postsynaptic density | GO:0098919 | Functional | 0 | 0 |
| structural constituent of ribosome | GO:0003735 | Functional | 0 | 0 |
| structural constituent of skin epidermis | GO:0030280 | Functional | 0 | 0 |
| succinyl-CoA hydrolase activity | GO:0004778 | Functional | 0 | 0 |
| synaptic receptor adaptor activity | GO:0030160 | Functional | 0 | 0 |
| syntaxin binding | GO:0019905 | Functional | 0 | 0 |
| syntaxin-1 binding | GO:0017075 | Functional | 0 | 0 |
| Tat protein binding | GO:0030957 | Functional | 0 | 0 |
| tau protein binding | GO:0048156 | Functional | 0 | 0 |
| telomerase RNA binding | GO:0070034 | Functional | 0 | 0 |
| thiamine pyrophosphate binding | GO:0030976 | Functional | 0 | 0 |
| thioredoxin peroxidase activity | GO:0008379 | Functional | 0 | 0 |
| thrombospondin receptor activity | GO:0070053 | Functional | 0 | 0 |

| | | | | |
|---|---|---|---|---|
| thyroid hormone binding | GO:0070324 | Functional | 0 | 0 |
| thyrotropin-releasing hormone receptor binding | GO:0031531 | Functional | 0 | 0 |
| Toll-like receptor 4 binding | GO:0035662 | Functional | 0 | 0 |
| toxic substance binding | GO:0015643 | Functional | 0 | 0 |
| transcription coactivator activity | GO:0003713 | Functional | 0 | 0 |
| transcription coregulator activity | GO:0003712 | Functional | 0 | 0 |
| transcription corepressor activity | GO:0003714 | Functional | 0 | 0 |
| transcription corepressor binding | GO:0001222 | Functional | 0 | 0 |
| transcription factor binding | GO:0008134 | Functional | 0 | 0 |
| transcription regulator inhibitor activity | GO:0140416 | Functional | 0 | 0 |
| transferase activity | GO:0016740 | Functional | 0 | 0 |
| transferrin receptor activity | GO:0004998 | Functional | 0 | 0 |
| transferrin receptor binding | GO:1990459 | Functional | 0 | 0 |
| transforming growth factor beta binding | GO:0050431 | Functional | 0 | 0 |
| transition metal ion binding | GO:0046914 | Functional | 0 | 0 |
| transketolase activity | GO:0004802 | Functional | 0 | 0 |
| translation activator activity | GO:0008494 | Functional | 0 | 0 |
| translation elongation factor activity | GO:0003746 | Functional | 0 | 0 |
| translation factor activity, RNA binding | GO:0008135 | Functional | 0 | 0 |
| translation initiation factor activity | GO:0003743 | Functional | 0 | 0 |
| transmembrane transporter binding | GO:0044325 | Functional | 0 | 0 |
| triglyceride binding | GO:0017129 | Functional | 0 | 0 |
| triglyceride lipase activity | GO:0004806 | Functional | 0 | 0 |

| | | | | |
|---|---|---|---|---|
| triose-phosphate isomerase activity | GO:0004807 | Functional | 0 | 0 |
| tRNA 2'-phosphotransferase activity | GO:0000215 | Functional | 0 | 0 |
| tRNA-specific adenosine deaminase activity | GO:0008251 | Functional | 0 | 0 |
| tryptophan-tRNA ligase activity | GO:0004830 | Functional | 0 | 0 |
| tubulin binding | GO:0015631 | Functional | 0 | 0 |
| tumor necrosis factor binding | GO:0043120 | Functional | 0 | 0 |
| type 1 angiotensin receptor binding | GO:0031702 | Functional | 0 | 0 |
| type 2 angiotensin receptor binding | GO:0031703 | Functional | 0 | 0 |
| type I transforming growth factor beta receptor binding | GO:0034713 | Functional | 0 | 0 |
| type II transforming growth factor beta receptor binding | GO:0005114 | Functional | 0 | 0 |
| U1 snRNP binding | GO:1990446 | Functional | 0 | 0 |
| U7 snRNA binding | GO:0071209 | Functional | 0 | 0 |
| ubiquitin binding | GO:0043130 | Functional | 0 | 0 |
| ubiquitin ligase inhibitor activity | GO:1990948 | Functional | 0 | 0 |
| ubiquitin protein ligase activity | GO:0061630 | Functional | 0 | 0 |
| ubiquitin-like ligase-substrate adaptor activity | GO:1990756 | Functional | 0 | 0 |
| ubiquitin-like protein ligase binding | GO:0044389 | Functional | 0 | 0 |
| unfolded protein binding | GO:0051082 | Functional | 0 | 0 |
| very-low-density lipoprotein particle binding | GO:0034189 | Functional | 0 | 0 |
| very-low-density lipoprotein particle receptor binding | GO:0070326 | Functional | 0 | 0 |
| vinculin binding | GO:0017166 | Functional | 0 | 0 |
| virion binding | GO:0046790 | Functional | 0 | 0 |

| | | | | |
|---|---|---|---|---|
| vitamin D binding | GO:0005499 | Functional | 0 | 0 |
| vitamin E binding | GO:0008431 | Functional | 0 | 0 |
| vitamin transmembrane transporter activity | GO:0090482 | Functional | 0 | 0 |
| X | Percent of amino acids in protein that are undetermined or atypical | Physiochemical | 0 | 0 |
| XExpo | Percent of absolute solvent-accessible area in protein that is undertermined or atypical amino acid absolute solvent-accessible area | Physiochemical | 0 | 0 |
| XMP 5'-nucleosidase activity | GO:0106411 | Functional | 0 | 0 |
| zinc ion binding | GO:0008270 | Functional | 0 | 0 |
| zinc ion sequestering activity | GO:0140486 | Functional | 0 | 0 |
| zymogen binding | GO:0035375 | Functional | 0 | 0 |

### 11 DNA Sequences

**Table S3. DNA sequences**

| Sequence | Structure(s) |
|---|---|
| CCAGGCAGTTGAGACGAACATTCCTAAGTCTGAAATTTATCACCCGCCA TAGTAGACGTATCA | Th, Th-Ch |
| CTTGCTACACGATTCAGACTTAGGAATGTTCGACATGCGAGGGTCCAAT ACCGACGATTACAG | Th, Th-Ch |
| GGTGATAAAACGTGTAGCAAGCTGTAATCGACGGGAAGAGCATGCCCA TCCACTACTATGGCG | Th, Th-Ch |
| GGTGATAAAACGTGTAGCAAGCTGTAATCGACGGGAAGAGCATGCCCA TCCACTACTATGGCG-CHOL | Th-Ch |
| CCTCGCATGACTCAACTGCCTGGTGATACGAGGATGGGCATGCTCTTCC CGACGGTATTGGAC | Th, Th-Ch |
| /5BIO/CCTCGCATGACTCAACTGCCTGGTGATACGAGGATGGGCATGCTC TTCCCGACGGTATTGGAC | Th, Th-Ch |
| AGTCAGGACATAGGCTGGCTGACCTTTGAAAG | Sq1 |
| CCATGTACCGTAACACTGTAGCATTCCACAGATTCCAGAC | Sq1 |
| TTTCATTGAGTAGATTTAGTTTCTATATTT | Sq1 |
| GTCAGGAAGAGGTCATTTTTGCTCTGGAAG | Sq1 |
| AACAGTTAGGTCTTTACCCTGATCCAACAG | Sq1 |
| ATACATACAACACTATCATAACATGCTTTA | Sq1 |
| GTGAATATAGTAAATTGGGCTTTAATGCAG | Sq1 |
| ACAACGGAAATCCGCGACCTGCCTCATTCA | Sq1 |
| CTCAGCAGGCTACAGAGGCTTTAACAAAGT | Sq1 |
| CAAAAGGTTCGAGGTGAATTTCTCGTCACC | Sq1 |
| GTTAGTAACTTTCAACAGTTTCAAAGGCTC | Sq1 |
| CTTGATACTGAAAATCTCCAAAAAAGCGGAGT | Sq1 |
| AAGACTTTGGCCGCTTTTGCGGGATTAAACAG | Sq1 |
| ACTTAGCCATTATACCAAGCGCGAGAGGACTA | Sq1 |
| TTAATTTCCAACGTAACAAAGCTGTCCATGTT | Sq1 |
| ACCAGACGGAATACCACATTCAACGAGATGGT | Sq1 |
| GTCAGAAGATTGAATCCCCCTCAACCTCGTTT | Sq1 |
| TAGAGCTTCAGACCGGAAGCAAACCTATTATA | Sq1 |
| AAATATTCCAAAGCGGATTGCATCGAGCTTCA | Sq1 |
| AGATTTAGACGATAAAAACCAAAAATCGTCAT | Sq1 |

| | |
|---|---|
| ACCCAAATAACTTTAATCATTGTGATCAGTTG | Sq1 |
| CCCCAGCGGGAACGAGGCGCAGACTATTCATT | Sq1 |
| GAGTTAAATTCATGAGGAAGTTTCTCTTTGAC | Sq1 |
| TTTCACGTCGATAGTTGCGCCGACCTTGCAGG | Sq1 |
| CGGGTAAAATTCGGTCGCTGAGGAATGACA | Sq1 |
| CATAAGGGACACTAAAACACTCACATTAAA | Sq1 |
| TTATGCGATTGACAAGAACCGGAGGTCAAT | Sq1 |
| AGGCTTTTCAGGTAGAAAGATTCAATTACC | Sq1 |
| TTAAGAGGGTCCAATACTGCGGATAGCGAG | Sq1 |
| GAGTAATCTTTTAAGAACTGGCTCCGGAACAA | Sq1 |
| AAAAGAATAACCGAACTGACCAACTTCATCAA | Sq1 |
| CATTATTAGCAAAAGAAGTTTTGC | Sq1 |
| CAACTAAAGTACGGTGGGATGGCT | Sq1 |
| TCATTTGCTAATAGTAGTAGCATT | Sq1 |
| GCCATTTGCAAACGTAGAAAATACCTGGCATG | Sq1 |
| AGCTAATGCAGAACGCGAGAAAAATAATATCCTGTCTTTC | Sq1 |
| GTACCAGGTATAGCCCGGAATAGAACCGCC | Sq1 |
| CAGTGCCCCCCCTGCCTATTTCTTTGCTCA | Sq1 |
| GCCAGCAGCCTTGATATTCACAAACGGGGT | Sq1 |
| AGTTTGCGCATTTTCGGTCATAGAGCCGCC | Sq1 |
| TAGAAAAGGCGACATTCAACCGCAGAATCA | Sq1 |
| ATGAAATGAAAAGTAAGCAGATACAATCAA | Sq1 |
| ATCCCAAAAAAATGAAAATAGCAAGAAACA | Sq1 |
| TAAGAACGGAGGTTTTGAAGCCTATTATTT | Sq1 |
| CTTATCACTCATCGAGAACAAGCGGTATTC | Sq1 |
| AGATTAGTATATAGAAGGCTTATCCAAGCCGT | Sq1 |
| CAGAGAGAACAAAATAAACAGCCATTAAATCA | Sq1 |
| AAAGTTACGCCCAATAATAAGAGCAGCCTTTA | Sq1 |
| AGGGAAGGATAAGTTTATTTTGTCAGCCGAAC | Sq1 |
| ATTAGCGTCCGTAATCAGTAGCGAATTGAGGG | Sq1 |
| AATCCTCAACCAGAACCACCACCAGCCCCCTT | Sq1 |
| TTATTCTGACTGGTAATAAGTTTTAACAAATA | Sq1 |
| GAGCCGCCTTAAAGCCAGAATGGAGATGATAC | Sq1 |

| | |
|---|---|
| TAGCAGCATTGCCATCTTTTCATACACCCTCA | Sq1 |
| ACCACGGATAAATATTGACGGAAAACCATCGA | Sq1 |
| TGAGTTAACAGAAGGAAACCGAGGGCAAAGAC | Sq1 |
| TGCCAGTTATAACATAAAAACAGGACAAGAAT | Sq1 |
| CAAATCAGTGCTATTTTGCACCCAGCCTAATT | Sq1 |
| ATTAGACGGAGCGTCTTTCCAGAGCTACAA | Sq1 |
| ATAATAACTCAGAGAGATAACCCGAAGCGC | Sq1 |
| TTAAAGGTACATATAAAAGAAACAAACGCA | Sq1 |
| TCACCGGAAACGTCACCAATGAATTATTCA | Sq1 |
| GTCTCTGACACCCTCAGAGCCACATCAAAA | Sq1 |
| AGGTGGCAGAATTATCACCGTCACCATTAGCA | Sq1 |
| CGCTAATAGGAATACCCAAAAGAAATACATAA | Sq1 |
| AGGCCGGAACCAGAGCCACCACCG | Sq1 |
| TTAGGATTAGCGGGGTGGAACCTA | Sq1 |
| ACCCTCATTCAGGGATAGCAAGCC | Sq1 |
| GTAGATTTGTTATTAATTTTAAAAAACAATTC | Sq1 |
| CTAAACAGGAGGCCGATAATCCTGAGAAGTGTCACGCAAA | Sq1 |
| CCAGTATGAATCGCCATATTTAGTAATAAG | Sq1 |
| ATTTCATGACCGTGTGATAAATAATTCTTA | Sq1 |
| GCTTAGAATCAAAATCATAGGTTTTAGTTA | Sq1 |
| GCGAATTATGAAACAAACATCATAGCGATA | Sq1 |
| ATTATCAGTTTGGATTATACTTGCGCAGAG | Sq1 |
| ACAGTTGTTAGGAGCACTAACATATTCCTG | Sq1 |
| ATGCGCGTACCGAACGAACCACGCAAATCA | Sq1 |
| AATACCTATTTACATTGGCAGAAGTCTTTA | Sq1 |
| TTAACCGTCACTTGCCTGAGTACTCATGGA | Sq1 |
| TCACACGATGCAACAGGAAAAACGGAAGAACT | Sq1 |
| GATAAAACTTTTTGAATGGCTATTTTCACCAG | Sq1 |
| ATTAGAGCAATATCTGGTCAGTTGCAGCAGAA | Sq1 |
| TGGAAGGGAGCGGAATTATCATCAACTAATAG | Sq1 |
| AAATTAATACCAAGTTACAAAATCCTGAATAA | Sq1 |
| CTACCTTTAGAATCCTTGAAAACAAGAAAACA | Sq1 |
| AAATAAGAACTTTTTCAAATATATCTGAGAGA | Sq1 |

| | |
|---|---|
| TTTCCCTTTTAACCTCCGGCTTAGCAAAGAAC | Sq1 |
| CTTTGAATTACATTTAACAATTTCTAATTAAT | Sq1 |
| CCAGAAGGTTAGAACCTACCATATCCTGATTG | Sq1 |
| CCTCAATCCGTCAATAGATAATACAGAAACCA | Sq1 |
| AGACAATAAGAGGTGAGGCGGTCATATCAAAC | Sq1 |
| CCAGCCATCCAGTAATAAAAGGGACGTGGCAC | Sq1 |
| CACCGCCTGAAAGCGTAAGAATACATTCTG | Sq1 |
| GATTTAGATTGCTGAACCTCAAAGTATTAA | Sq1 |
| ATTTGCACCATTTTGCGGAACAAATTTGAG | Sq1 |
| TTACCTTTACAATAACGGATTCGCAAAATT | Sq1 |
| TATATAACGTAAATCGTCGCTATATTTGAA | Sq1 |
| AACATTATGTAAAACAGAAATAAATTTTACAT | Sq1 |
| GCATCACCAGTATTAGACTTTACAGTTTGAGT | Sq1 |
| CGGGAGAATTTAATGGAAACAGTA | Sq1 |
| TCATATGCGTTATACAAAGGCGTT | Sq1 |
| AGAATATCAGACGACGACAATAAA | Sq1 |
| GAATTCGTGCCATTCGCCATTCAGTTCCGGCA | Sq1 |
| AATCATACAGGCAAGGCAGAGCATAAAGCTAAGGGAGAAG | Sq1 |
| GGAAGGGGGCAAGTGTAGCGGTGCTACAGG | Sq1 |
| GGCGATGTTTTTGGGGTCGAGGCGAGAAA | Sq1 |
| CTGGTTTGTTCCGAAATCGGCATCTATCAG | Sq1 |
| CTAACTCCCAGTCGGGAAACCTGGTCCACG | Sq1 |
| GTGCTGCCCCAGTCACGACGTTTGAGTGAG | Sq1 |
| ATTGACCCGCATCGTAACCGTGAGGGGGAT | Sq1 |
| CAGGAAGTAATATTTTGTTAAAAACGGCGG | Sq1 |
| ACCGTTCATTTTTGAGAGATCTCCCAAAAA | Sq1 |
| CCTTTATCATATATTTTAAATGGATATTCA | Sq1 |
| TATCAGGTAAATCACCATCAATATCAATGCCT | Sq1 |
| TAAATTTTTGATAATCAGAAAAGCACAAAGGC | Sq1 |
| AGTTTGAGATTCTCCGTGGGAACAATTCGCAT | Sq1 |
| ACGGCCAGTACGCCAGCTGGCGAACATCTGCC | Sq1 |
| AGCTGCATAGCCTGGGGTGCCTAAGTAAAACG | Sq1 |
| TATAAATCGAGAGTTGCAGCAAGCGTCGTGCC | Sq1 |

| | |
|---|---|
| AGCACTAAAAAGGGCGAAAAACCGAAATCCCT | Sq1 |
| GGCCCTGAAAAAGAATAGCCCGAGCGTGGACT | Sq1 |
| AAGTGTAATAATGAATCGGCCAACCACCGCCT | Sq1 |
| TTCGCTATTGCCAAGCTTGCATGCGAAGCATA | Sq1 |
| CCCGTCGGGGGACGACGACAGTATCGGGCCTC | Sq1 |
| ACCCCGGTTGTTAAATCAGCTCATAGTAACAA | Sq1 |
| AGACAGTCCATTGCCTGAGAGTCTTCATATGT | Sq1 |
| CCAATAGGAAACTAGCATGTCAAGGAGCAA | Sq1 |
| AGGAAGATCATTAAATGTGAGCGTTTTTAA | Sq1 |
| TCGACTCTGAAGGGCGATCGGTGCGGCCTC | Sq1 |
| GAGAGGCGACAACATACGAGCCGCTGCAGG | Sq1 |
| TGAGTGTTCAGCTGATTGCCCTTGCGCGGG | Sq1 |
| ACTGTTGGAGAGGATCCCCGGGTACCGCTCAC | Sq1 |
| TTCATCAACGCACTCCAGCCAGCTGCTGCGCA | Sq1 |
| AATTCCACGTTTGCGTATTGGGCG | Sq1 |
| AAAGCCGGCGAACGTGTGCCGTAA | Sq1 |
| GCGCGTACTTTCCTCGTTAGAATC | Sq1 |
| ACTTTTGCATCGGTTGTACTTTTTTTAACCTGTTTAGGACCATTA | Sq1 |
| GATACATTTCGCTTTTTTGACCCTGTAAT | Sq1 |
| GAGTAATGTGTAGGTAAAGATTTTTTGTTTTAAATATG | Sq1 |
| AAGCGAACAATTGCTGAATATAATGCTGTATTTTTTTGTGAGAAAGGCC GG | Sq1 |
| ACAAGAGTTTTTTTCGCGTTTTAATTCAAAAAGA | Sq1 |
| TGGATAGCAAGCCCGATTTTTAATCGTAAACGCCAT | Sq1 |
| CAAAAATAATTTTTTTTGTTTAGAC | Sq1 |
| CAGAGGGGGTTTTGCCTTCCTGTAGCCAGCT | Sq1 |
| CCGCTTCTGGTTTTTTCGTTAATAAAACGAACTAAATTATACC | Sq1 |
| GGAATTAGAGCTTTTTTTTCAGACCAGGCGCGTTGGGAAGATTTTTTTTC CAGGCAAAGC | Sq1 |
| TGTCGTCTCAGCCCTCATATTTTTTTCGCCACCCTCAGGTGTATC | Sq1 |
| ACCGTACTCAGGTTTTTGATCTAAAGTTT | Sq1 |
| GAGAATAGAAAGGAACAACTATTTCTCAAGAGAAGGA | Sq1 |
| AGGAGTGTAAACATGAAAGTATTAAGAGGCTTTTTTTGCGAATAATAAT TT | Sq1 |

| | |
|---|---|
| ACAACCATTTTTTCATACATGGCTTTTAAGCGCA | Sq1 |
| AGAACCGCATTTACCGTTTTACCGATATATACGTAA | Sq1 |
| TGCCACTACTTTTTTTGCCACCCTC | Sq1 |
| GAACCGCCTCTTTACCTAAAACGAAAGAGGC | Sq1 |
| AGGACAGATGATTTTTTCACCAGTAGCACCATTACCGACTTGA | Sq1 |
| TAATCGGCCATCCTAATTTTTTTTTTTTTCGAGCCAACAACGCC | Sq1 |
| AACATGTAATTTTTTTTGAAACCAATCAA | Sq1 |
| TTTTATTTTCATCGTAGGAATTTTTAGCCTGTTTAGTA | Sq1 |
| GCGAGAAAATAAACACCGGAATCATAATTATTTTTTTCGCCCAATAGCA AG | Sq1 |
| TTTTATCTTTTTTATCCAATCGCAAGAGTTGGGT | Sq1 |
| TTGCTTCTTATATGTATTTTACGCTAACGGAGAATT | Sq1 |
| AACTGAACATTTTTTTTGAATAACC | Sq1 |
| CATAAATCAATTTAGTCAGAGGGTAATTGAG | Sq1 |
| ATTAAGACTCCTTTTTAATATACAGTAACAGTACCGAAATTGC | Sq1 |
| AATCATGGTCATTTTTTTTTTTGCCCGAACTCAGGTTTAACTTTTTTTTCA GTATGTTAG | Sq1 |
| CTGTCCATTTTTATAATCATTTTTTTCTTAATGCGCCCACGCTGC | Sq1 |
| GCGTAACCACCATTTTTGAGTAAAAGAGT | Sq1 |
| CAAACTATCGGCCTTGCTGGTTTTTGAGCTTGACGGGG | Sq1 |
| CCAACGTCATCGGAACCCTAAAGGGAGCCCTTTTTTTGAACAATATTAC CG | Sq1 |
| GCCAACATTTTTTCCACTATTAAAGAAATAGGGT | Sq1 |
| ACGGGCAAGTTCCAGTTTTTTCTGACCTGCAACAGT | Sq1 |
| GCCACGCTGTTTTTTTACCAGTGAG | Sq1 |
| CCAGGGTGGTTTTGCAAATGAAAAATCTAAA | Sq1 |
| GACAACTCGTATTTTTTCCTGTGTGAAATTGTTATCCGAGCTC | Sq1 |
| GTGTCGTAGACACTCCCAATTCTGCGAACCCATATAACAGTTGATGTGT CGTAGACAC | Sq1 |
| GTGTCGTAGACACCCATAAATCAAAAATCCAGAAAACGAGAATGAGTG TCGTAGACAC | Sq1 |
| GTGTCGTAGACACGAACGAGGGTAGCAACGCGAAAGACAGCATCGGTG TCGTAGACAC | Sq1 |
| GTGTCGTAGACACGGGATTTTGCTAAACAAATGAATTTTCTGTATGTGT CGTAGACAC | Sq1 |

| | |
|---|---|
| GTGTCGTAGACACGAGAGGGTTGATATAAGCGGATAAGTGCCGTCGTGTCGTAGACAC | Sq1 |
| GTGTCGTAGACACGCAGGTCAGACGATTGTTGACAGGAGGTTGAGGTGTCGTAGACAC | Sq1 |
| GTGTCGTAGACACGCGCCAAAGACAAAAGTTCATATGGTTTACCAGTGTCGTAGACAC | Sq1 |
| GTGTCGTAGACACTTTTTTGTTTAACGTCTCCAAATAAGAAACGAGTGTCGTAGACAC | Sq1 |
| GTGTCGTAGACACTAAACCAAGTACCGCATTCCAAGAACGGGTATGTGTCGTAGACAC | Sq1 |
| GTGTCGTAGACACAGTAGGGCTTAATTGAAAAGCCAACGCTCAACGTGTCGTAGACAC | Sq1 |
| GTGTCGTAGACACAGTCAATAGTGAATTTTTAAGACGCTGAGAAGGTGTCGTAGACAC | Sq1 |
| GTGTCGTAGACACCAATATAATCCTGATTGATGATGGCAATTCATGTGTCGTAGACAC | Sq1 |
| GTGTCGTAGACACACATCGCCATTAAAAAAACTGATAGCCCTAAAGTGTCGTAGACAC | Sq1 |
| GTGTCGTAGACACTTGATTAGTAATAACATTGTAGCAATACTTCTGTGTCGTAGACAC | Sq1 |
| GTGTCGTAGACACCGGGCGCTAGGGCGCTAAGAAAGCGAAAGGAGGTGTCGTAGACAC | Sq1 |
| GTGTCGTAGACACATCCTGTTTGATGGTGGCCCCAGCAGGCGAAAGTGTCGTAGACAC | Sq1 |
| GTGTCGTAGACACGTAACGCCAGGGTTTTAAGGCGATTAAGTTGGGTGTCGTAGACAC | Sq1 |
| GTGTCGTAGACACTTTAAATTGTAAACGTATTGTATAAGCAAATAGTGTCGTAGACAC | Sq1 |
| GTGTCGTAGACACAAATTTTTAGAACCCTTTCAACGCAAGGATAAGTGTCGTAGACAC | Sq1 |
| CGTACTGACCGTTTCGCTGCTTTTTGAAACACCAGAACGAGAGGCTTGCCCTGACGA | Sq1 |
| GCAGCGAAACGGTCAGTACGTTTTTTTTTTTTTTTTTTTT/3BIO/ | Sq1, Tu, Rd, Bx, Sq-Apt1, Sq3, Sq-Apt2, Sq2 |
| ATCAAATCACCGGAACCAGACACCCTCA | Tu |
| GAACCGCCTACATGGCTTTTGATGGGGTCAGTGCCT | Tu |
| CAGAACCGAATTTTCTGTATGGGAAGTGAGAATAGA | Tu |
| TGAGAGGTGTATCACCGTACGCCACCCT | Tu |

| | |
|---|---|
| CCAGAACGTAGGAATACCACATTCTACGAGGCATAG | Tu |
| GTGTAATCAACGTAACAAAGGAGAAACA | Tu |
| AAGGGATATATTCGGTCGCTGGGATCGT | Tu |
| CACCCTCAGCGATTATACCAAGCGGCCTGATAAATT | Tu |
| TAAGCAAATTTCCGGCACCGCTTCCATTCAGGCTGC | Tu |
| AAAACAATCATATGTACCCCAGATTGTA | Tu |
| TAAGATTCGAGCTTCAAAGCGGATTAGA | Tu |
| GAGTACCTCAAAGAATTAGCAAAAATCGGTTGTACC | Tu |
| CGTGCCAGGAAAAACCGTCTATCAAATCAAGTTTTTTGGG | Tu |
| GCAACTAACTCACATTAATTAAACCTGT | Tu |
| CAAAGGGCCTGCATTAATGAATCGGTGCCTAA | Tu |
| TGAGTGAGCTGTTGGGAAGGGCGATCGCACTC | Tu |
| ATAATCAACCAGTAGCACCATTACATTGACGG | Tu |
| AAGCGTCAACCCTCAGAGCCACCAATCTTTTC | Tu |
| AGTAAATGCCACCCTCAGAGCCACAAGTATAG | Tu |
| CCCGGAATTAACAGTGCCCGTATAGTTCCAGT | Tu |
| TTGAGATTAGTAGTAAATTGGGCTGGATATTC | Tu |
| ATTACCCACGAAATCCGCGACCTGTCATCTTT | Tu |
| GCATAACCAACAACTAAAGGAATTCAGACGTT | Tu |
| GACCCCCAGCAGCGAAAGACAGCATCGCCCAC | Tu |
| CAGCCAGCTATTTAAATTGTAAACCGTAAAAC | Tu |
| TAGCATGTACATTATGACCCTGTAAATCATAC | Tu |
| GCGTTTTAAGCAACACTATCATAATTCATCAG | Tu |
| AGGCAAGGTTAATTGCTCCTTTTGCAAATATC | Tu |
| GCCGTAAATCCAACGT | Tu |
| AGGCCGGACTTATTAGCGTTTGCCCCCTCAGA | Tu |
| GCCGCCACAGTCTCTGAATTTACCAACAGTTA | Tu |
| TTTTCAGGAGTTTTGTCGTCTTTCGCGAATAA | Tu |
| ATGCCCCCTCGAGAGGGTTGATATCACCCTCA | Tu |
| TTTAATTTATTACAGGTAGAAAGACCCTCGTT | Tu |
| TACTTAGCAATCTTGACAAGAACCTGAGATGG | Tu |
| TAATTTTTACAATGACAACAACCATCGGAACG | Tu |
| AGGGTAGCGAATACACTAAAACACCTCCATGT | Tu |

| | |
|---|---|
| TTTGTTAAATCGGCCTCAGGAAGATCGGTGCG | Tu |
| GCGGGAGAATCGATGAACGGTAATGTTAATAT | Tu |
| TACCAGACGAAGCCCGAAAGACTTATAAGAGG | Tu |
| TCATTTTTGCATTAACATCCAATAATACTTTT | Tu |
| GCGGGGAGATTAAAGAACGTGGACGCACTAAATCGGAACC | Tu |
| GGCCTCTTAAGTGTAAAGCCTGGGGCCAACGC | Tu |
| CACTAGGCGGTTTGCGTATCGAGCCG | Tu |
| GAAGCATACGCTATTACGCCAGCGGGGACG | Tu |
| ATAGCCCCAACGTCACCAATGAACATTCAA | Tu |
| GAAAGCGCCAGAACCACCACCAGTTCGGTC | Tu |
| CGATCTAAGATAGCAAGCCCAATGGCGGAT | Tu |
| AAGTGCCGTGCCTATTTCGGAACCAGAATG | Tu |
| ACAACATTCAACTTTAATCATTGACCTTCA | Tu |
| TCAAGAGTCGGAACGAGGCGCAGACGAAAG | Tu |
| TTGCGCCGTCACGTTGAAAATCTAGCGTAA | Tu |
| AGGCAAAAAACGGCTACAGAGGCCCGATAG | Tu |
| ACGACAGTAATTCGCATTAAATTGGAGCAA | Tu |
| ACAAGAGAAGCCTTTATTTCAACCTACTAA | Tu |
| ATTAAGAGGACGATAAAAACCAATAACGGA | Tu |
| TAGTAGTAGCGGATGGCTTAGAGCAAAAAG | Tu |
| AGCCCCCGAACAAGAGTC | Tu |
| ATAGCCGAATCAGAGAGATAACCCAATAAGAA | Tu |
| ACGATTTTCTACAATTTTATCCTGTCATTACC | Tu |
| GTAATTCTGTAGGGCTTAATTGAGATTTAATG | Tu |
| GCGCCCAAAATAATCGGCTGTCTTAAGGTAAA | Tu |
| AGTTTGAGATTAGAGCCGTCAATAAATCGTCA | Tu |
| CTTTTACATGGAAGGGTTAGAACCATTTTAAA | Tu |
| GTTTGAAATATAACTATATGTAAATCGCTATT | Tu |
| AATTAATTAAATTAATTACATTTAAACAGTAC | Tu |
| ATGTGAGCCCGTCGGATTCTCCGTGTGCCAAG | Tu |
| AGACAGTCCAATATGATATTCAACAACATTAA | Tu |
| TAAATATTCCCTCAAATGCTTTAAACAGTTGA | Tu |
| TTCCCAATAGTAGATTTAGTTTGAAAGGCCGG | Tu |

| | |
|---|---|
| TCCACGCTAGCAGGCGAAAATCCTGTCACGCTG | Tu |
| CTTGCATGGACTCTAGAGGATCCCGCAAGCGG | Tu |
| CCCCGGTTTGAGCTAAACACATCACGC | Tu |
| AAAGGTCCCTGCATTAACCGTTCCAGCCAT | Tu |
| CGCTAATACAAAGTTACCAGAAATACATAAAGGTGG | Tu |
| CACCCAGTTGTTTAACGTCAAATAATTGAG | Tu |
| CTCAACAGTCCAGACGACGACAATGTAGAA | Tu |
| ACCAATCTAGCAAGCAAATCAGCTATTTTG | Tu |
| CTAATAGTAACATTATCATTTTTATACTTC | Tu |
| TGAATAATCGGGAGAAACAATAAACATCAA | Tu |
| TGGGTTATACCGACCGTGTGATAGCCAACG | Tu |
| GAAAACATTCCCTTAGAATCCTGCTTAGGT | Tu |
| TGACAACGAGTACAACAGGAAAGACATTCT | Tu |
| GGACCATAAATCCCAACAGAGACATCGCCA | Tu |
| ATAATCCCATTGCACCTTGCTGACTAACAA | Tu |
| TTGAACGTCTGCAAAAATACCGTCTAAAGC | Tu |
| CACCACACCCGCCGATCAGAGCGGG | Tu |
| CTTTAAGGTCAGACGGCCAGTTACAAAGAAACAATTTTGC | Tu |
| CCTTGCTGAGACTCCTCATCGAGAAAGCCTAATTTATTGG | Tu |
| AGAAGCACCAGTACAGCATTTTCGAAGTACCGCACTCAAG | Tu |
| AAAATCTTTGAAAGAATTGCGTAGAAACAGTACATCGGGT | Tu |
| GATGACATTATACCAAATTCGACAAAAATAAAGAAGGACA | Tu |
| CAACGTGTATCGGTTCTTTTTCAAATTAGGCAGAGAACTA | Tu |
| GCTTGTTCCATTAAAAAATCAATATCGCGAGAAAATATCA | Tu |
| TTCATTTTTAAATGCAATGCCTGAGCTGTTTAGCTATATT | Tu |
| GATCTGAACGCCATCAAAAATAATTTAGCTATTTTTGAGA | Tu |
| GACGTCCAGAGGGGGTAATAGTAAACTTTACAAACGTCAG | Tu |
| CCCTGATGTTTTAAATATGCAACTAAAAATCAGGTCTTTA | Tu |
| GTGAAGAGACGGGCAACAGCTGATTATAGCTGTTTCCTGT | Tu |
| AGATGAGTTGGGTAACGCCAGGGTTAGGTCACGTTGGTGT | Tu |
| AGCACGTTTTCATCGGCATTAGCCGCCG | Tu |
| CCAGCATTAAATCCTCATTAAAGCCTATTATTCTGA | Tu |
| ATGTACCGGACAGCCCTCATAGTTCCAAAAAAAAGG | Tu |

| | |
|---|---|
| AACAGTTTTGCTCAGTACCAAGGAACCC | Tu |
| CTTATGCGCGTTAATAAAACGAACAATAGCGAGAGG | Tu |
| AAGGGCGCATAGGCTGGCTGTGAATTAC | Tu |
| CTCCCTTAAACAGCTTGATATTTGAGGA | Tu |
| CTAAAGACAAGGCACCAACCTAAAACGGTCAATCAT | Tu |
| ATCAGCTCGCATCTGCCAGTTTGATGGCGAAAGGGG | Tu |
| ATTTCATTGCCTGAGAGTCTTTTGTTAA | Tu |
| CTTTGCAAAGCGGATTGCATCTTAATTG | Tu |
| CTGAATATAAAGGTGGCATCAATTGCAAGGATAAAA | Tu |
| AGGGTGGTGTGTTGTTCCAGTTTGGATTTAGAGCTTGACG | Tu |
| GATGAATTCCACACAACATATGGGCGCC | Tu |
| AGGGTTGATTTTCTTTTCACCAGTATTGTTAT | Tu |
| CCGCTCACTGCTGCAAGGCGATTAGGCGCATC | Tu |
| CTGTAGCGCCGTAATCAGTAGCGAATGGTTTA | Tu |
| AAACAAATGACAGGAGGTTGAGGCGCGTCAGA | Tu |
| ATTCCACATAACACTGAGTTTCGTGATTAGGA | Tu |
| TTAGCGGGTGAAAGTATTAAGAGGATATTCAC | Tu |
| CGTACTGACCGTTTCGCTGCTTTTTAAAATCTAATTTTAAGAACTGGCTA<br>CGGTGTA | Tu |
| CAGACCAGGAACCGAACTGACCAAACGTAATG | Tu |
| GTGAATTTAAAAGGAGCCTTTAATCCTGTAGC | Tu |
| CCACTACGTTTTTCATGAGGAAGTCTTTCGAG | Tu |
| GTAACCGTATTTTTTAACCAATAGACAAAGGC | Tu |
| TATCAGGTTTAGAACCCTCATATATTGGGGCG | Tu |
| AGTCAGAATGCAAAAGAAGTTTTGTGGGAAGA | Tu |
| CGAGCTGAAATGCTGTAGCTCAACACTATTAT | Tu |
| CGGCGAACCCCGAGAT | Tu |
| CGAAGCCCAATAATAAGAGCAAATAAAC | Tu |
| AGCCATATACGAGCGTCTTTCCAGCAAGCAAGCCGT | Tu |
| TAAGAGAAAACGCCAACATGTAATTATATTTTAGTT | Tu |
| TTTTAACGGGTATTAAACCAGCCAGTAA | Tu |
| AAATCCTTATTTAGAAGTATTAGAATGTTTAGACTG | Tu |
| ACGTTTTGCACGTAAAACAGCTCGTATT | Tu |

| | |
|---|---|
| AATTATCGCAAGACAAAGAAATGTGAGT | Tu |
| GAATAACCTACCTTTTTTAATGGATTTTCAGGTTTA | Tu |
| GCCTTCCTTTGACCGTAATGGGATTTCCCAGTCACG | Tu |
| AAAGTTAATGCCGGAGAGGGCGCGTCTG | Tu |
| GATAGAATGACCATAAATCAAAGTACGG | Tu |
| TGTCTGGAGCAAATGGTCAATAACTAATGTGTAGGT | Tu |
| CCGCCTGGGAAATCGGCAAAATCCGAGCGGGCGCTAGGGC | Tu |
| ACGTATTCGTAATCATGGTCGCCCTTCA | Tu |
| GTGGTTCCCCCTGAGAGAGTTGCACGGGTACC | Tu |
| GAGCTCGATGTAAAACGACGGCCAGGGAACAA | Tu |
| TGAGTTAAGCCCTTTTTAAGAAAAGCAAAGAC | Tu |
| CAACGCTATATTTATCCCAATCCAACAAGAAT | Tu |
| TATTTAACTATAAAGTACCGACAATCCTTATC | Tu |
| ATTCCAAGATTTTCATCGTAGGAAAATCTTAC | Tu |
| ATTTGAGGTGCCCGAACGTTATTATACCATAT | Tu |
| CAAAATTACAGATGAATATACAGTACAATTTC | Tu |
| CAAATCCATCATCTTCTGACCTAAAATCGCCA | Tu |
| ATTTGAATTTGCTTCTGTAAATCGTGCTGATG | Tu |
| ACGGCGGAGTAGCCAGCTTTCATCCGTTCTAG | Tu |
| CTGATAAAATTCAAAAGGGTGAGACCATTAGA | Tu |
| GAAAACGAGCGTCCAATACTGCGGGATAATAC | Tu |
| TACATTTCAGTTTCATTCCATATAACAGTTCA | Tu |
| GTGTAGCGGTTTGATG | Tu |
| AACGGAACAAAGTCAGAGGGAATGAAAA | Tu |
| TAGCAGCCAAATCAAGATTAGTTGATATAGAAGGCT | Tu |
| CATGTTCAATTCTTACCAGTATAAAAATAAGGCGTT | Tu |
| TATCATCCTAATTTACGAGCATAAACAA | Tu |
| AAGAAACCAAATATCTTTAGGAGCAACCTCAAATAT | Tu |
| GATTTCCTGATTGTTTGGATGCGGAACA | Tu |
| AAATACCTTTTTAACCTCCGTGAAAACA | Tu |
| TAGCGATAAGAAGATGATGAAACAACGGATTCGCCT | Tu |
| TGGAAATACAGAACAATATTACCGGTAGCAATACTT | Tu |
| GACCGACCAGTAATAAAAGGAACGCTCA | Tu |

| | |
|---|---|
| CAAAAGCAGCAAATGAAAAAAACGAACC | Tu |
| ACCAGCAGACTGATAGCCCTAAAATAGAACCCTTCT | Tu |
| ATTAAAGGTGCTTTCCTCGTTAGACGCTTAATGCGCCGCT | Tu |
| CTTTAGTAAAAGAGTCTGTCGGAGGCCG | Tu |
| GTATAACGGATTTTAGACAGGAACATCAGTGA | Tu |
| GGCCACCGGATTAGTAATAACATCCCTTGCTG | Tu |
| AACACCCTCAATAATAACGGAATATTACGCAG | Tu |
| GAAGCCTTTTTACAGAGAGAATAAATTAACTG | Tu |
| TTATACAAGCTAATGCAGAACGCGAGAAAAAT | Tu |
| AATATCCCCGGTATTCTAAGAACGGAGGTTTT | Tu |
| GTTATCTAACCAGAAGGAGCGGAACAATTCAT | Tu |
| CAATATAAGCTTTGAATACCAAGTCAATTACC | Tu |
| GAGAGACTAAGAATAAACACCGGACATATGCG | Tu |
| TGAGCAAAGCTTAGATTAAGACGCATAGGTCT | Tu |
| GTAATATCCCTACATTTTGACGCTGATTCACC | Tu |
| AGTCACACTGAAAGCGTAAGAATAAGTCTTTA | Tu |
| TGAGAGCCCCCTCAATCAATATCTTGAGGAAG | Tu |
| ATGCGCGAAAGATAAAACAGAGGTTGCCACGC | Tu |
| CGTACTATACGAGCAC | Tu |
| CTAAAGGGCCGATTGAGGGAGGGAAGGTAAATCATTAGCA | Tu |
| CTGGCGAGAAATTCATCAGAATCAAGGAAATAGCAAACAATCAATAGA | Tu |
| CGCGTAACCAACATATAAAAGAAACGTAAGCAG | Tu |
| AAAGGAAGGGAACTTATAAATCAAAAGAATAG | Tu |
| GTCGAGGTAAATTATTCATTAAAGGTGAATTAAGAGCCAGCAAA | Tu |
| GGGAAAGCCCAGCGCCAAAGACAAAAGGGCGAACCATCGATAGC | Tu |
| GCTGGCAAACCACGGAATAAGTTTATTTTGTCTAGCTATCTTAC | Tu |
| ACAGGGCGTATGTTAGCAAACGTAGAAAATACGGAAACCGAGGA | Tu |
| TGGCAGATGAGTAAAAAATCGCCATATTTAACTGTAATTTAGGACAAC | Rd |
| AGGCCACCTCACCAGTTCAACAGTGGCGTTTT | Rd |
| AAAGGGAACCGTCTATCATTATAATCAGTG | Rd |
| CGTCAAAGGGCGAAAACATTCTGGCCATCCAC | Rd |
| ACCAAAAGTACCCGACTTGAGCCACAACCATCAACCGATAGACTCCAA | Rd |
| TATTAAAGAACCGGTCGCAAGGTGTATTCGGT | Rd |

| | |
|---|---|
| AGAATGCGCAGCGCAGTACTTATAGCTCACACATTCAACTTCATAACC | Rd |
| GAGCCGCCAGTTGAGAAAAACGAACTGTGGTGCTGCGGCC | Rd |
| TTACCTGCCGCGCCTGTGCTGTTCTGGTGACTCTAACGGA | Rd |
| CCTTACACAGCAAATCGTTTGGGTGGTAAAAC | Rd |
| TCTTAGCCTCCTGTTGCTCGTCATAAACATC | Rd |
| AATTCGCGTCTGGCCTAGCTTTCACAGGTCAGTACCTTTA | Rd |
| AATCATAATTACAACAAACGCCTAGCCAACGCCACACGACGCTCAATC | Rd |
| AATGTTGATTAAGCAAGCAAATTCCCGACTATTTTGACCAGTAATA | Rd |
| CACGCTGGTTAAACGGGTAAACAATTTGGGAAGGCTTGCACATCGGAA | Rd |
| ACGTTAATTTTAGGAATGTCACTGAGCCAGCGGTGCCGGTGTGGTGCC | Rd |
| GAATGGCTTAGAGCTTGCGGCTAAAGGTT | Rd |
| ACCCAGCCATTGCTGGATTATGAACGCGAAGGGCTTAGAACAAAG | Rd |
| GTCTGAAAAACAGGAAGAAGGCTTCGGGTAGGAATCATTACCGCGCCC | Rd |
| ACCTTCTACCCTACTGCGGGATCTTACCAGTATAAAGAAAAAGC | Rd |
| AAGAGACAGAGATAGAGACCTGAAAAATCAAGCTATTTTG | Rd |
| GAGTGTTGTTCTCCGAGTGGTCAGTTTGGAAC | Rd |
| TTTTCACGGGCACCAAAGTGGCGAAAATCCT | Rd |
| TTTTCACCCCTAAAACAAAGAATAAGCACCATTACAGCGTCAGACTGT | Rd |
| AGTTTTAAGACGATAATCTGGTCACAACCAGCTTACGGCTATGCCGGG | Rd |
| GGCATCAGGGAGGTGTCGAGGCATATAGCGAGAGGCTTAT | Rd |
| GCGGATTACCAGCCGGGTCACTGTTGAGTAAGAGCGCCCTAAGAGAG | Rd |
| AGCGAACCCAGATATAAAACGCTCTTTTGAATGGCCAGAA | Rd |
| AAAAACAGCTTGATACCGATACTTAGCGGGTT | Rd |
| ATCCCACGGCAGCAACCGCAAGAAATGACTTGTAGAACGT | Rd |
| CCCTCGGCCAACGCGCAACTAAAGTAATAATT | Rd |
| CAGCGTGGTGCTGCAGGTCATTGGAAACCAAAAGTAAGAG | Rd |
| TATGAGCATCAGCGGGGCGCTTTCTAACCGTGCATCTGCC | Rd |
| GCGCATCGGCACTCAATCCGCCGGGCAACGGGAACAGCGGTTGCGG | Rd |
| ATATCTATTATCTGGTCAGTTGGCTTATCTAATCTTTCCTTACCGCAC | Rd |
| CACCCGTGTAACCTTGCTTCTCCTAAAACATAATACCGTCCTGAA | Rd |
| TTTCCAGTAATGAGTGAGCTAACTGAGCCGGAAGCATAAA | Rd |
| TTTCCCGTTCAACTTTAATCATTTTATGCGATTGTAAA | Rd |
| ATAGCTGGAAATTGGAGGTTTCCCTCAGAACAGTATATATACGC | Rd |

| | |
|---|---|
| GAGGGGGTGTATCACCTACCAGACCGGAACGTGCCGGGTC | Rd |
| TTGCTCAGCTGGATAGTACAAAGGATTGCCTGAGAGTCTTAGATGG | Rd |
| ATAGGTCACGTTGGTGGGAGCAAAGAGCGGAATCGTCAT | Rd |
| TCTTACGTTTTTATTTTCATCCTGAATAACCTCAAATATC | Rd |
| TACATAAATCAATTAGTTATCAGCATCAATAG | Rd |
| AGGAACCCCACCCTCATATGGGATCAACATACCACATTAATTGTGTGT | Rd |
| CTCTTACCGTGAAGTTGTACTCAGTTACCAGAGCACATCC | Rd |
| TCAGAGGGGACGACGATTTTGCCATAGTAAAA | Rd |
| AAATTTGCAAAAGAAGCAGAGGTCCTATCTAT | Rd |
| CAGAAAACCAAGAGAAGTAATCGTAAATTTTGGCTATACTTAACGGGG | Rd |
| TGCTGATCTTTAGGAGTAGATAATCAGAGGGTTTTGAACC | Rd |
| GTGCCACGTTTGCACGAGCCTAATTTGCCCTGAACAAGCACATCACCT | Rd |
| TGACCTTTAATTAATTCATATGGTCGGCTTAGATAACTATATGGAATT | Rd |
| AAATGTCGTCTTTCCCCGAAGAGTCAATAG | Rd |
| GGTTGGCCGTTCCGGCATTCCACATTTCGCCAAGTACGCT | Rd |
| GCTCACAAAACGCGGTCCGTTTAAGGGTAA | Rd |
| CGGCCTCAGGAAGCGCTGGCAGCCTCCGGTCC | Rd |
| GAGGGTAAGAGATCCGTCCAATACTGAAT | Rd |
| TCATCGAGAACAAAGTACAGCAAATGAAAAAT | Rd |
| CCGCCAGCACCCTCATGAAACAGCAAAAAAATCCCGTAAAATTTGTAC | Rd |
| TTTCAGCGGTAAATGAATTTTCTGGAGCCACCAGTTGGGC | Rd |
| ACCAGAGCCGCCATGTGGCCTTTAGTGGGAAAGTGCCATGTTTCGTCT | Rd |
| CGTATCGCACTCCAGCGGATAAGTAGCTCAAA | Rd |
| TGTTTAGATACCAGGCCAGAATTAATGCCGGA | Rd |
| GACCATAAATAAGTTTAGCATGTCGACCCTGTAATACTTT | Rd |
| GAGGATTTAGAAGTATTTAAATCCAATTGAGCTGAGTTAA | Rd |
| AACTGAACAATGGAAGTACCATATCAAAATTACTGAGAGCCAGCATTT | Rd |
| ACCTTTTTAACCAAGAAACATCTCTTAAAACGAAAAGCCAGCGCCAAA | Rd |
| GACAAGCCTCTGTTATGTTGGCACGGAAAAATCGGTCTGAGAGACT | Rd |
| ATCGACATGGATCAAACTTAAATTGAGACGCATTTGTAAC | Rd |
| AGTTAAACGATGCTGAAAAGCCGAGAACCGCATGTACCGTAACGGA | Rd |
| CGCTTCTGCCAGGCAAGCCGTCGAGAACCGCCTCCCTCAG | Rd |
| TGCAACCGTTCTAGCTGATACTTTCCGGCAC | Rd |

| | |
|---|---|
| ATTCATTTCAACATATCAAAGACACCACGGTCTTTCCAGTAACAAA | Rd |
| CACCCTCCACAGGCTTACCAGTCCCGGAA | Rd |
| TCACCATCAATATGATATTCGGGTCAGGTT | Rd |
| GGTTAGCCCGAACGTTATTTTGCGTAATAAGATTAGAGAG | Rd |
| CTGTTGCGAGAAAAATACCAGTTACAAAATAA | Rd |
| AGAAATAATAGATTTTATATTATTTATCCAGCGCATTAGA | Rd |
| ACAACATAAAGGTGGCAATTACCTGAGAAC | Rd |
| TGGTGAAGAGACGGTCTGTAGCATGACAACGTCACCAATGGTACAACG | Rd |
| AACTGACCTTTGTGAGAGATAGACTTTCTCCG | Rd |
| GTAGCTGCTTCAGCAGCACCACCGGAGGGTTGAGCCCGGAATAGGTAA | Rd |
| ATTCGCCAGCAACTGTCGCCACCCCACCCTCAGAGCCCAT | Rd |
| AAAAAAGGGTGAGAATAGGATTAGCGGGTG | Rd |
| CCTTGAGTAACAGGCTTAATCAACGCAAGGAT | Rd |
| GCCGAATTCGGGACAAGAATTGGATTATACTT | Rd |
| TAGAAAATACATGCCCAGGTTTAACGTAAA | Rd |
| ATCGCGCAGAGGCTAAAACATGTTGCAGTCGATCACCGTC | Rd |
| ACAAAGGGCGACATTCATGCTGATGCAAAAAC | Rd |
| TTAGCCAGGGATAGCAACAACGCCAATCATAACGACCTGC | Rd |
| TGTGTACAACGGTGTCGAAATCCGGGGAACCG | Rd |
| AACTTTGATGAGTTTCCACCGTAACAGAATACCGGATATTCACGG | Rd |
| AATCAAAATCACCACAAGAATCGGCGAAAC | Rd |
| CTCAGAACTGGGAAGGCGGGCCTCTTCGCTATGGCGAAAG | Rd |
| GAAAGTATTGTCGGTGGCGATGTAGGTAAAGATTCAAAT | Rd |
| AATAACAGAGGCATTTAATAAGAGAAAAACAGTAATCCTGATTGTCAA | Rd |
| GCAATTCATCATTTTAGTACCATAGGACAATCCAAATAAG | Rd |
| CAGTAACACATCGGGATAAATAAGGCGCCAGT | Rd |
| TCACCCTATACCGACAAGACTCTACCAGATGAATATA | Rd |
| CAATTGAATACCAAGTCTTATTACAGCAAACG | Rd |
| GTGAATTATTGAGGGAGGGAAGGTCGGTCCAATCGCAAGA | Rd |
| AAACTTTTTCAAATAACTTAGCAAATATTTCCACAG | Rd |
| TCCATGTTTATTTGTATCATCGCCTGATAAAT | Rd |
| AAACAGGACAGATGAGACCAGGCGCATCCA | Rd |
| CAATGTGCTGCAAGGCGATTTCAGAGGTGGAGTGCCATCTCTCACCGG | Rd |

| | |
|---|---|
| CGGGAACGGATCAGCTTACGCAACTTTGCCACTCAGACAT | Rd |
| TTTTAAATGCAATGCCTGAGTAATAAGAGGCTGAGTAACTATTTC | Rd |
| TTACCAGAATGAAAATAGCAGCCTAATAACATAATATAAAAGATGATG | Rd |
| AAACGATTCGTGTGAGAAACAATAACGGATTCGCCTGA | Rd |
| TTGCTAGAAATTTAATGGTTTGAACAGCAGCGAAAGACAGGGGAGTTA | Rd |
| CGAGGGTATTCATCTTCTGACCTAACGCGAGA | Rd |
| GGGGATAACCTGTTTAGCTATATTTTCATTTATTAGAT | Rd |
| GAGATTTTAGTTAATGCAACGGAATTATTAGCAAAA | Rd |
| CCGGAGGACTAATACCAAGCGCGAAACAATCTAGAGTAAAAAACCATC | Rd |
| CTTCATCATGACAAGACAAGTTTGCCTTTCATTAGCAAGG | Rd |
| AGCTAGCGATCAGGTTCCGAGGCTGGCTGAC | Rd |
| CAGAACCAGTTGGGTAACGCCTATAACAGTTGCAAATGGT | Rd |
| ACATTTCGATTCCCAATTCTGCGGTACGGTGTCTGGAAATTCTGTA | Rd |
| TCGAGTTACGTCAAAAAGGAAACCGAGGAAACGCAATAAGGAACCGG | Rd |
| ATGATTTTTTGTTTAAAATAAGAATAAACTCG | Rd |
| CCAGTGCCAAGCTTAACCATAGCCGGTCACCA | Rd |
| GGAACCTAGGTTGAGGCAGGTCAGACGATTGCAACTAAAAACGAGTA | Rd |
| CTGTTTAGTATCATATTTTGCGTAACGGAAAACTGGC | Rd |
| AAGGCCGCTGCCGCATGCCCAGTTATACAAATGGTTTTGAAGCGTTGC | Rd |
| CGCCTAAAGAGGATGATTAGAGCCCATTAAAG | Rd |
| TCTGCTCAAAGCTTTGACCCCCAGCGATTCAG | Rd |
| AGCGCGTTTTCATCGCGATTACCCAAATCAACGTAACATTCAGTGA | Rd |
| TTTCCCAGTCACGACGTTGTAAAAGCATTTTCCCCTTATT | Rd |
| CAACACTAAATGCAGATACATAACGATTCATCGCCAGCATCCAAGGGT | Rd |
| ACAACATTGTTTCATTTGACAGGATTATTCTGAAAGCCAC | Rd |
| GCTCAACATGTTTTAAATGCAAACGGAAATGGCCTTGATGAATGGAA | Rd |
| CCAGTCAGGAGCTTGCCCTGACGAGAAGGCAGAAAGAAC | Rd |
| GATTATTGCTGAATATAATGACAGGTAGAAAGCCAAAAG | Rd |
| GCCGACAATGAATACGTAATGCCACTACGAATTGAAAATCTCCAAA | Rd |
| TCATTGAATCCCCCTTAAGAGGTCATTTTT | Rd |
| AGCGAATAAGTTTATTTTGTCACATTGCTTTC | Rd |
| GATAGCAGGTCACCAGTACAAACTAGCCCAAT | Rd |
| ATTAATTGTATCGGTATTAAGAC | Rd |

| | |
|---|---|
| TCCCTTATAAATTAGAATCCTTGAGTCG | Rd |
| AAATTAAACGCCACCCTCAATCAATAGTCTTTAATGCGCGAACTG | Rd |
| CTAGTCGAAGAGCCCAACGCTAACG | Rd |
| TGAGGCAGTATTAACACCTACATTTAATGCCTGCAACA | Rd |
| TACTGGTAATCAAAAACCCGAACGTCGATAA | Rd |
| CGTACTGACCGTTTCGCTGCTTTTTGTTTGATGAATCGGCAAAATTTGCG TATTGG | Rd |
| ATTTACCGTTCCAGTAATAGCGGGGGAAGTTGGGTAAATACACTAATCA A | Bx |
| CGAGTTAGCGTTTACAAATTCTTAATGCCACAGACTTTGGTAGCAACCC | Bx |
| CACAAAGCCAAAGCCTGAACTATATGTAAATTGAATTT | Bx |
| ATAGCAAGCCCCAGAACCTTTCGGACCTTGAGACCAGAGGCCA | Bx |
| GTAATTTCAGCAGGCTCCAGTGAATTGACAAGAACCGGATTCA | Bx |
| AGGCAGATAGAAACGCAATAATAAAAGGGCTAAA | Bx |
| AAGCTAATTTGCTAACACAATGAAATAGCAATTAA | Bx |
| TGAATTAATAAACCCACAAATTT | Bx |
| AGCATTAGTTTAAGAGCATGTTAGCAAACGTCGGA | Bx |
| AAATGGAATTGCGCCTTAAATC | Bx |
| TATACGTAACAAAGCTGAATTGGGTTAGACT | Bx |
| TCAAACTAAAAAACAGCCAAAGCTCTTACCGAAAGGAACCGTTGAAAA TCTCCAAA | Bx |
| ATTAAACAATCATTACCGCTTTAACAGTCAAGTTA | Bx |
| TTGAAGAGTAAACCAACTTTGAAAGATAA | Bx |
| TCACAGGGAGCGCATAAAGTTGCGCGAGGTGGCCTTTAATTG | Bx |
| ATCCCATCATTCCAAGAATTAATCGCGAACATCA | Bx |
| GCTGAACCGAATTGTGTCGAAATCCTATC | Bx |
| CAATAAACTGAACAAGAAATTAAACAAAACATAAA | Bx |
| ATCGGCCTGATCACTCATCTTTGACAAAAGAAATAC | Bx |
| ACAGTAGACAACGCCTGTAGCCCGGAATTTAGGATGCGT | Bx |
| GGCTTCTGTCCAGACTTATAT | Bx |
| ACATGTAACCCATGTACCGTACACCCTCAACATGATAAGTTTGACA | Bx |
| TCGCCATGTTTCGTCACCAGTACTCAGGTGAGACTGATACAGAGAC | Bx |
| GTACCAGTATAGACAGCCCTCATAGAGGGTTTCAGTACCTGA | Bx |
| TCAGTGTGATAAATAAGAACGCCA | Bx |

| | |
|---|---|
| ATCGAACGAGTTCATGAGTTTTGCGATATAAGTATAGCATTC | Bx |
| TTACAAAAACGATTTCAAC | Bx |
| AGAAGGAAGGTGTATCACCGTACAAACTGGCT | Bx |
| TCATACCGATCCGATATTCGGCATATCTTTTCATAATCAATAAATAAGA | Bx |
| TCGGTCGTAGGCAAGTACCCAATGATCAGAGCCACCACCCTCAG | Bx |
| TACACCTTTTGTCAGATACGTAAACATATTCTTTTGCGGAACAAACCAA | Bx |
| AAGGCTTGCCCTATACTTCTGAATAATGCCTG | Bx |
| GTATTAATTTATCCGAATAGAAGCCCTTACCCAAAAGAACTGGGTTTAC | Bx |
| CTTTGTTTGGATTGACGAGAAACACACAACTC | Bx |
| TTATCATACAGAAAGCGTTTTGCAAATCACGCAAACATTACCATTAGCA | Bx |
| TCAGATGTCAAAATCTTGCGGGAGGTTTTGAAGAATAATAATTTTT | Bx |
| CAAAAAGTTTGAGTAACCCAG | Bx |
| ATATAATGAAGGGTTAGAACCTACCATAATGG | Bx |
| ATTGCCCGAACGTTATTACA | Bx |
| TAATGCAGATACTTACAACATAACTGGCACAACTAAT | Bx |
| AATCAGCAGAAGATAAATGTAAACATCAGTGTAGGATGTTTACATTTTT GTAGATTTTCAGGTTTAACACA | Bx |
| ACAATGCTGTCCTTTAAGTCACGTTGTGAGCGAGTAACTTGA | Bx |
| ACCAATTTTATAACGCCAAAATTAGAGACAAATATTCAG | Bx |
| ATACGTGCCCTAAAACATCTTGAGA | Bx |
| ATCGTTTTTAATAGCGAGAGGCGAA | Bx |
| GCGAAAAGAAGTTTACCCGAGGCAATACCACATTCAAC | Bx |
| ACGACCAGACCTGAAAGCGTTAAAGGGCAAGTTTTCAAA | Bx |
| GGTTGCTAAGGGAAGAAAGCGGGGCGCG | Bx |
| ACGCAAACTATTCATAAATATTCAATCAAAATAGG | Bx |
| TTATAATGCGCCGCTACCGGGCGC | Bx |
| ATCCCGTTAGAATCAGAGTTAGAGCGGTGCCGCCAAATCGAAAAACAC GT | Bx |
| CAGCATTTAACGGGGTCAGTGACCTATTTTGACGA | Bx |
| GCAGGTCGAGTGTACTGGTAAAAGTATT | Bx |
| GATACCACCGGTTTGCCCCGACTTGAGCCATTTGGGAAAGCGACA | Bx |
| GATATTCACATGGCTTTTGATCCTCAAG | Bx |
| TAATTGAATCTGAGTAGAAGAACTCAAA | Bx |

| | |
|---|---|
| GCGGTTTAGCTATATTTCCCTGTAATGCCTGCTGG | Bx |
| CAACCGCGCTACCGTTGTAGCAATACTTGCCCCCCTCAGATA | Bx |
| AGCGAGAATGACTATTAATTAAGATAATTCGCCAACAGTTTTGATAAGA | Bx |
| ACGTTCGCAAATGGTCAGAAGCCTCCCTCATATGAACGACCC | Bx |
| CGTGGCGCATCAATTCTACTATCAGAGCAGGCCGGTTTGAGAGTAA | Bx |
| AGAAGGTCTTAAATGCAATACTTTTGCGGGAATAACCTCTGG | Bx |
| TAGCTCCAACTTTGTTAAGAGGGTAGCTATTAGACAGT | Bx |
| AGCTCAACATGTCCCTGAGAGAGTTGCAGCCC | Bx |
| CGAAGAATAGGTATCGGAGGAACGCCATCAATTAATAT | Bx |
| CTGGTTTTGAGACGTCGTGCCAGCTGTTTCCTGTGTGAA | Bx |
| GTTGAACATTTTCCCAGTCACGACCAACTGT | Bx |
| CCCACCGCCTGGTTTAAATATGCAATTGC | Bx |
| TTCGCTTTCCAGTATCAAAACGACGGCTGATTGCAAGCGGTCCACG | Bx |
| TGGTACGGTGGAATATAGGTAGAAAGATTCAATTACCTTCAATAGAT | Bx |
| AAGCATATAACTAGAGCCGTATGCGATTTTAAGTATT | Bx |
| CATAGCTGCAAGTTGAACTAAAGCATCACCTAACAGTGCAGG | Bx |
| ATTGTTCGGGGTTATCTTGGGAAGAACAAATGAAAAATAGGAATTGA | Bx |
| CTTGGGATAGTTGCTCCGTCAGGAGGAATTAAGACGACGATAAAA | Bx |
| TCACATCTGCCAGTTTGTCGCGTC | Bx |
| TGGGAAGGATGGGCGCATCGTCCAGCTT | Bx |
| CGTTATTTTTTTTTTTTTTTTTTTTTTTTTTTTTTAACCTAAAACGAGACTAC | Bx |
| TAACGCGTTTGGAAGCCCTTTTGCTCCAATACTGCGGA | Bx |
| CGGAACCCGTCCGTAATCGCTATTGGCGAGTAAAGCCTGGG | Bx |
| TCGAACCAATCCTCAGGCTTCTGGTCGACCGTAATCATGGT | Bx |
| ACGAAATAATAGGGGACCAAAGCGCCAAGCTTGCATGCTCAC | Bx |
| ACCAATATATTTTAGTTTTCC | Bx |
| TCATCAAAAAACAGGAAGATTCATTGCC | Bx |
| TCATAAGCAATAAAGCCATAGTAGGGAAAGCCTTTCCTTGAGAAGTG | Bx |
| AGCAGCCCCACATTAAATGGTGTAGGCGATCCAGGGTACGAGCCGG | Bx |
| AAAGGCTCAAAAGGGTGAGAAATAAAGC | Bx |
| TGAGAGTAGTAATGTGTAGGTCAAAAAC | Bx |
| GCACCGTCCACCCTCAGAACCCCGCCGCTTAATAAT | Bx |
| GAATCAAGAACCGCCTCCCTCGGTTGAG | Bx |

| | |
|---|---|
| TCAGACTCCGGAACCAGAGCCTGGCCTT | Bx |
| ATTACTAGAAAACGCTCATTTAAACTCCAT | Bx |
| GACAACGGAATTTTAAATAACATAAAAACAGGGTATTAATC | Bx |
| TTCGGAAATTATTCATTGTTTTCAATTC | Bx |
| ATATTGCCCAATAGCAAAGCGAACTGCACCCTAACGAGCGGTCAGCCCAA | Bx |
| AGTTTATCAGCAAAATCACCAGATAGCATTCGGGT | Bx |
| CAGCGCCAATTATCACCGTCATTTAGCG | Bx |
| ATTATGATCATTTGAAAGGAGAGGGCGCGTCTGTCCATC | Bx |
| TAAGCTATTTCTCCCGATATTTGCGAATATA | Bx |
| AAAAGGAGTGACAATCCAAAGAGAGAAAA | Bx |
| CATACAAACAAAAATCAGTAGCGCAAAGGTGAAA | Bx |
| ATTAAAGCCAGTTTGCCTTTCGGTTATTGACAACCGATAACCGAGGCCGAATA | Bx |
| GGGAGGCTTGGCAGCGACCTC | Bx |
| AGTAGTACTCATTCAAAAGGAAATTTCTGACA | Bx |
| GGAAGAGCCGAATCAGTTTAGAGCTTTGTCAACGC | Bx |
| GGAGGTTGAGAATCAAAAAATCCTGGTGGTTATCGGCCAAAATTTCTAGA | Bx |
| AGCTGAAGTTGTACAAAGATTATCAGGTGTATAAGCCTGTAGAACC | Bx |
| AAATGAAAACAAAGTTATTGCTAAACAACTTTTTTGTT | Bx |
| ATGATTTCTTTCTGCAGGTGCCGGAAACCAGGGACGACA | Bx |
| AAAGGTAACGCGGTGCGGGCCT | Bx |
| GATAGCTAGCATTAGACGGGAG | Bx |
| GCAGCCTTTACAGATAAGACAATTCGCAGAACGTTTTTTTTAGATTCACATTAACT | Bx |
| GCGCCTTAGAAACCAAGCGGAATTTTTTTAATATGGGGAGAGGCGG | Bx |
| CGATTGACGGTAGCATTAACATCCGCAAAATCCATCAAATGCCGGAAAT | Bx |
| AGTAAGAATGAACACCCTGAACAAATCTTTATTATCACTGATTATTTTTTTTTCAACTTA | Bx |
| AATCATAGGGTTATATTTTAGTATCATATG | Bx |
| CGAGCTCGCGCGCCTGGTCACCTCAAATATCAAAGTCA | Bx |
| AACAATAATTTTTTTTTTTTTTTTTTTTTTTTTTTTTAATCAATAATCGATCG | Bx |
| AGTAATAAGATACCGACAATGCA | Bx |

| | |
|---|---|
| TATACGCGAGTAAAGAAATTGCTTTTTTTTTTTTTTTTTTTTTTTTTTTTCAAGC | Bx |
| CGCAGCTACAGAATTGAACATACATAAAG | Bx |
| TTGCTTTCCGACAATGACTTTTTTTTTTTTTTTTTTTTTTTTTTTTGAAC | Bx |
| CGGTTTTTTTTTTTTTTTTTTTTTTTTTTTTTCCAGGCGCATAGGCTGATGAACG | Bx |
| TCGCCCATTAAAGGGCAGACGGTCAATCAGGACAGGCT | Bx |
| ACAAGTTTTTTTTTTTTTTTTTTTTTTTTTTTTAGCCGCCACCAGTTACGAGACA | Bx |
| AGACGTTTTGAAGGCTTATCCTTTTTTTTTTTTTTTTTTTTTTTTTTTTTGGAAA | Bx |
| GATCGTCACGGCTAACAA | Bx |
| AGAAACCTTTTTTTTTTTTTTTTTTTTTTTTTTTTAGTTAATGCCCCTTCAGCTAAA | Bx |
| TTATCCTAATAACCACCTAACAGTGCCCGTATAAAC | Bx |
| CTAATACGAAGGCACCTTTTTTTTTTTTTTTTTTTTTTTTTTTTTGCCGT | Bx |
| GAACGCTTTTTTTTTTTTTTTTTTTTTTTTTTTTCCCTCATTTTCAAGCC | Bx |
| ATAGACAATTTCATTTTTTTTTTTTTTTTTTTTTTTTTTTTTTTTAAAGGAATA | Bx |
| GTCCAACATGCTGCCTAGCCACCCTCAGAGCCACCA | Bx |
| ATTCTGAAGAACCGCCACCCTAATAGGAATTTAGGACCG | Bx |
| CGGGATTATACCAAGCGCGTTTTTTTTTTTTTTTTTTTTTTTTTTTTTTGAGGA | Bx |
| GAATTACCTTTTTTAATGGATAACCTAAAGTAAATTTTCGGGG | Bx |
| TCAATTATTTTTTTTTTTTTTTTTTTTTTTTTTTTGCCTGTTTATCACATGT | Bx |
| ATTGTGATCAGTTTTTTTTTTTTTTTTTTTTTTTTTTTTTTAGACCGGAA | Bx |
| TTAGCGAATGGAAATTTTTTTTTTTTTTTTTTTTTTTTTTTTACCA | Bx |
| GTAGGTGGTTCCGAATTTTTTTGCGT | Bx |
| AGCATTTTTTTTTTTTTTTTTTTTTTTTTTTTGGATTGCAT | Bx |
| CAGAGGGGGTTTTTTTTTTTTTTTTTTTTTTTTTTTTTAGTTCAGAAAACGGTC | Bx |
| TAAACTTTTTTTTTTTTTTTTTTTTTTTTTTTTAACCACCACACCCGGTGT | Bx |
| ATTCTAAATCGGAACTTTTTTTTTTTTTTTTTTTTTTTTTTTTCAGATTTGAATCTTT | Bx |
| ACAGGAGGCCGATTTTTTTTTTTTTTTTTTTTTTTTTTTTTATCGTCCGCT | Bx |
| TATCCCTTCTGTAATAAAAGGGACGATT | Bx |

| | |
|---|---|
| AAGCTTTTTTTTTTTTTTTTTTTTTTTTTTTTCATTAGATACATCTGC | Bx |
| ACTAGCTTCACATATGTGTAATCGTAAAACTAG | Bx |
| TGGAACTTTTTTTTTTTTTTTTTTTTTTTTTTTTAGGTGAGGCGCCCT | Bx |
| GTTCTGATAGGCACAGACAATATTGATA | Bx |
| AACCTTTTTTTTTTTTTTTTTTTTTTTTTTTTATAAAAATTAAGCA | Bx |
| GGATGGCTTAGAGCTTAATTGCTTCTGGAATGAGTGAGCAAGTTTAAGT | Bx |
| TTGCTTTTTTTTTTTTTTTTTTTTTTTTTTTTCATGTCAATAAGCG | Bx |
| ATTGGTCCCCGGTTTTTTTTTTTTTTTTTTTTTTTTTTTTATCGGCAAAATC CACT | Bx |
| TGTTCCATTTTTTTTTTTTTTTTTTTTTTTTTTTTTACAAACGGCCATTT | Bx |
| GGAGCGCCAGGTTTGATTTAACACACCGAACGAACCACGCGCGAACCA GTT | Bx |
| CATATAAAAGAAAGATATAATTTTCAGAGA | Bx |
| CCAGCTTTTTTTTTTTTTTTTTTTTTTTTTTTTAAGAGTCCACTATTTT | Bx |
| CAGCTTTCCGGCACCGAAGATCGCCTTATATGTT | Bx |
| TGTTATTTTTTTTTTTTTTTTTTTTTTTTTTTTCCACTACGTGAATTCA | Bx |
| AATCAGCTCATTTTTTCATTAAATTAAAGACGTC | Bx |
| ACCGTTTTTTTTTTTTTTTTTTTTTTTTTTTTCCTAAAGGGAGCCATAC | Bx |
| TCTAGCTGATAAATTATATGATACCATCACTAAAGCATACA | Bx |
| AAGGCAAAGAATTAAATAAATCCCCGATCGGGAGCAGGAAC | Bx |
| AGGATGAGGGAGGTTTTTTTTTTTTTTTTTTTTTTTTTTTTGGATT | Bx |
| GTGGCAATTTTTTTTTTTTTTTTTTTTTTTTTTTTGGTATTCTAAGA | Bx |
| CGTCACGCACTCGCTGTCTTCATT | Bx |
| GGTACGCCAACGCTCGCAGATTCACCATTGGGT | Bx |
| TAACGGAATCCAAATCCAATCGCAGAC | Bx |
| AGAAAACCATAGCGATAGCTTTTTCAACATCAGTCTGATAAGCTA | Bx |
| ATGAGAATCCTTGAAAATTTT | Bx |
| TAGATTAAAGACAAAATAAGAATAAACACTTGAGAA | Bx |
| GCCATTAAAAATCGCCTGCTGCTGAAGTTGGCAAA | Bx |
| AAGAGGCAGGTTTAGTACCGCACACTGAATTTAACGCGTTAAGAACGC G | Bx |
| TCGGCCTTGCTGGTTAAAAGAGTACTAT | Bx |
| CAGACTGATGTTGACAGTGAGGCCACCGAGAATATCCAGAACAATATT ACCATAAT | Bx |

| | |
|---|---|
| TTTTTGCCAGCCATTGCAACAGGAAAAGA | Bx |
| TAAATCGAAGGTGGAGAAAGGTTGACGAGCACGTATAACGTGCGGCGA A | Bx |
| CTTTGAGTGAAAACAGTATTAATTTGAAACACAGAGGCTGAA | Bx |
| GCTGAGAAGAGTCAATAGGCTGATGATA | Bx |
| AAAAAACAAGAGAATCGATATTTTTACCCTGACCA | Bx |
| TGTCAAATACCCCACCTGAGCAAAAGAAGATGAACATTTAATAA | Bx |
| TCGTCGCTATTAATTAATTAATTTCATCTTCTGTTTCAGCTGCTTTTGGGA TTCCGTTG | Bx |
| TCCCAAAAGCAGCTGAAGGGATTTTAGACTAA | Bx |
| GTAATAACATCACTTCTTTCAGCCGCTGTCACACGCACAG | Bx |
| GTGACAGCGGCTGAAACCTCCGGCTTAGGTTGGTCTGAGAAAGAGGCC CC | Bx |
| ACGCTGTCGGTGAGTGTAAAATGTTTAGACTGAATGCTT | Bx |
| ATACCAGTCAACGAACTAACGGA | Bx |
| GTGGGCTGCGGTTGTCGCTCACAATTCCACACCGCTCAGCACTATCATT | Bx |
| CAGGCCCCAGCCACGCTGAGAGCCAGCAGAAATCTACG | Bx |
| GGTGGATTGACGGATTCTCCGTGGTTGCGAACG | Bx |
| TAAAGTTTTTTTTCTCAGGCGGATATTTTTTTTTTTTTTTTTTTTTTTTTTT TTGCGG | Bx |
| CGTTAGTTTTTTTAAACATAGCCCCTTTTTTTTTTTTTTTTTTTTTTTTTTTT CAGC | Bx |
| GAACAGGGCGATGGCTTTTTTTTTTTTTTTTTTTTTTTTTTTTTTAGT | Bx |
| GGATTCGCCTGATTGCTTGAATTATTTCC | Bx |
| GAGGCCCGCTTTTTTTTTTTTTTTTTTTTTTTTTTTTAAACAAAGTCAGA | Bx |
| TAACCCTCGTTTTGCCAAAAAGTAGTCAGTTTAGAATTATTTCAACGCA AGG | Bx |
| ATGCTCCTAATCCTACACTGATGTTTA | Bx |
| CGTACTGACCGTTTCGCTGCCTTGTCTTAATAAAGGACGTAAAATATCTT TAGGACTG | Bx |
| AACTTCGTATAATCGTGATTTAGCATT | Bx |
| AATCAGTTAATTTTATTCATCATTTTTTTTGGAAGAAACCTGGGCAACAG CCAGTGCCATTCGCCAT | Bx |
| TAACGTCAGAAGTACTTGAGATTTTTTTTAATACTGCCTAAGTTTCATGC TGCAAACGCCAGCTGGCG | Bx |
| AGGTTGCTTCTGTAAATTTTTTTTTTTTTTTTTTTTTTTTTTTTAAATTTAA TGGTTTGAAATCAGA | Bx |

| | |
|---|---|
| TTGATGGAAATACCTACATTTTGATGAAATGATTCTTTTTTGGGGTATTTGACAAACTGACA | Bx |
| CAGCAGATTTGGCGACCTGCTCCATGTTACTTAGCTTTTACTCACCGACAGCGTTGAATGTT | Bx |
| GACCTATTCATTACCCAATTTTTTAATGCTAAATCACGATTAGCATTAATTTAGGATAGTAAGGCAA | Bx |
| GCGCGTACTTTCCTCGTTAGAATC | Sq3 |
| AAAGCCGGCGAACGTGTGCCGTAA | Sq3 |
| AATTCCACGTTTGCGTATTGGGCG | Sq3 |
| TGAGTGTTCAGCTGATTGCCCTTGCGCGGG | Sq3 |
| GAGAGGCGACAACATACGAGCCGCTGCAGG | Sq3 |
| TCGACTCTGAAGGGCGATCGGTGCGGCCTC | Sq3 |
| AGGAAGATCATTAAATGTGAGCGTTTTTAA | Sq3 |
| CCAATAGGAAACTAGCATGTCAAGGAGCAA | Sq3 |
| AGACAGTCCATTGCCTGAGAGTCTTCATATGT | Sq3 |
| ACCCCGGTTGTTAAATCAGCTCATAGTAACAA | Sq3 |
| CCCGTCGGGGGACGACGACAGTATCGGGCCTC | Sq3 |
| TTCGCTATTGCCAAGCTTGCATGCGAAGCATA | Sq3 |
| AAGTGTAATAATGAATCGGCCAACCACCGCCT | Sq3 |
| GGCCCTGAAAAAGAATAGCCCGAGCGTGGACT | Sq3 |
| AGCACTAAAAAGGGCGAAAAACCGAAATCCCT | Sq3 |
| TATAAATCGAGAGTTGCAGCAAGCGTCGTGCC | Sq3 |
| AGCTGCATAGCCTGGGGTGCCTAAGTAAAACG | Sq3 |
| ACGGCCAGTACGCCAGCTGGCGAACATCTGCC | Sq3 |
| AGTTTGAGATTCTCCGTGGGAACAATTCGCAT | Sq3 |
| TAAATTTTTGATAATCAGAAAAGCACAAAGGC | Sq3 |
| TATCAGGTAAATCACCATCAATATCAATGCCT | Sq3 |
| CCTTTATCATATATTTTAAATGGATATTCA | Sq3 |
| ACCGTTCATTTTTGAGAGATCTCCCAAAAA | Sq3 |
| CAGGAAGTAATATTTTGTTAAAAACGGCGG | Sq3 |
| ATTGACCCGCATCGTAACCGTGAGGGGGAT | Sq3 |
| GTGCTGCCCCAGTCACGACGTTTGAGTGAG | Sq3 |
| CTAACTCCCAGTCGGGAAACCTGGTCCACG | Sq3 |
| CTGGTTTGTTCCGAAATCGGCATCTATCAG | Sq3 |

| | |
|---|---|
| GGCGATGTTTTTGGGGTCGAGGGCGAGAAA | Sq3 |
| GGAAGGGGGCAAGTGTAGCGGTGCTACAGG | Sq3 |
| AATCATACAGGCAAGGCAGAGCATAAAGCTAAGGGAGAAG | Sq3 |
| GAATTCGTGCCATTCGCCATTCAGTTCCGGCA | Sq3 |
| AGAATATCAGACGACGACAATAAA | Sq3 |
| TCATATGCGTTATACAAAGGCGTT | Sq3 |
| CGGGAGAATTTAATGGAAACAGTA | Sq3 |
| TATATAACGTAAATCGTCGCTATATTTGAA | Sq3 |
| TTACCTTTACAATAACGGATTCGCAAAATT | Sq3 |
| ATTTGCACCATTTTGCGGAACAAATTTGAG | Sq3 |
| GATTTAGATTGCTGAACCTCAAAGTATTAA | Sq3 |
| CACCGCCTGAAAGCGTAAGAATACATTCTG | Sq3 |
| CCAGCCATCCAGTAATAAAAGGGACGTGGCAC | Sq3 |
| AGACAATAAGAGGTGAGGCGGTCATATCAAAC | Sq3 |
| CCTCAATCCGTCAATAGATAATACAGAAACCA | Sq3 |
| CCAGAAGGTTAGAACCTACCATATCCTGATTG | Sq3 |
| CTTTGAATTACATTTAACAATTTCTAATTAAT | Sq3 |
| TTTCCCTTTTAACCTCCGGCTTAGCAAAGAAC | Sq3 |
| AAATAAGAACTTTTTCAAATATATCTGAGAGA | Sq3 |
| CTACCTTTAGAATCCTTGAAAACAAGAAAACA | Sq3 |
| AAATTAATACCAAGTTACAAAATCCTGAATAA | Sq3 |
| TGGAAGGGAGCGGAATTATCATCAACTAATAG | Sq3 |
| ATTAGAGCAATATCTGGTCAGTTGCAGCAGAA | Sq3 |
| GATAAAACTTTTTGAATGGCTATTTTCACCAG | Sq3 |
| TCACACGATGCAACAGGAAAAACGGAAGAACT | Sq3 |
| TTAACCGTCACTTGCCTGAGTACTCATGGA | Sq3 |
| AATACCTATTTACATTGGCAGAAGTCTTTA | Sq3 |
| ATGCGCGTACCGAACGAACCACGCAAATCA | Sq3 |
| ACAGTTGTTAGGAGCACTAACATATTCCTG | Sq3 |
| ATTATCAGTTTGGATTATACTTGCGCAGAG | Sq3 |
| GCGAATTATGAAACAAACATCATAGCGATA | Sq3 |
| GCTTAGAATCAAAATCATAGGTTTTAGTTA | Sq3 |
| ATTTCATGACCGTGTGATAAATAATTCTTA | Sq3 |

| | |
|---|---|
| CCAGTATGAATCGCCATATTTAGTAATAAG | Sq3 |
| CTAAACAGGAGGCCGATAATCCTGAGAAGTGTCACGCAAA | Sq3 |
| GTAGATTTGTTATTAATTTTAAAAAACAATTC | Sq3 |
| ACCCTCATTCAGGGATAGCAAGCC | Sq3 |
| TTAGGATTAGCGGGGTGGAACCTA | Sq3 |
| AGGCCGGAACCAGAGCCACCACCG | Sq3 |
| GTCTCTGACACCCTCAGAGCCACATCAAAA | Sq3 |
| TCACCGGAAACGTCACCAATGAATTATTCA | Sq3 |
| TTAAAGGTACATATAAAAGAAACAAACGCA | Sq3 |
| ATAATAACTCAGAGAGATAACCCGAAGCGC | Sq3 |
| ATTAGACGGAGCGTCTTTCCAGAGCTACAA | Sq3 |
| CAAATCAGTGCTATTTTGCACCCAGCCTAATT | Sq3 |
| TGCCAGTTATAACATAAAAACAGGACAAGAAT | Sq3 |
| TGAGTTAACAGAAGGAAACCGAGGGCAAAGAC | Sq3 |
| ACCACGGATAAATATTGACGGAAAACCATCGA | Sq3 |
| TAGCAGCATTGCCATCTTTTCATACACCCTCA | Sq3 |
| GAGCCGCCTTAAAGCCAGAATGGAGATGATAC | Sq3 |
| TTATTCTGACTGGTAATAAGTTTTAACAAATA | Sq3 |
| AATCCTCAACCAGAACCACCACCAGCCCCCTT | Sq3 |
| ATTAGCGTCCGTAATCAGTAGCGAATTGAGGG | Sq3 |
| AGGGAAGGATAAGTTTATTTTGTCAGCCGAAC | Sq3 |
| AAAGTTACGCCCAATAATAAGAGCAGCCTTTA | Sq3 |
| CAGAGAGAACAAAATAAACAGCCATTAAATCA | Sq3 |
| AGATTAGTATATAGAAGGCTTATCCAAGCCGT | Sq3 |
| CTTATCACTCATCGAGAACAAGCGGTATTC | Sq3 |
| TAAGAACGGAGGTTTTGAAGCCTATTATTT | Sq3 |
| ATCCCAAAAAAATGAAAATAGCAAGAAACA | Sq3 |
| ATGAAATGAAAAGTAAGCAGATACAATCAA | Sq3 |
| TAGAAAAGGCGACATTCAACCGCAGAATCA | Sq3 |
| AGTTTGCGCATTTTCGGTCATAGAGCCGCC | Sq3 |
| GCCAGCAGCCTTGATATTCACAAACGGGGT | Sq3 |
| CAGTGCCCCCCCTGCCTATTTCTTTGCTCA | Sq3 |
| GTACCAGGTATAGCCCGGAATAGAACCGCC | Sq3 |

| | |
|---|---|
| AGCTAATGCAGAACGCGAGAAAAATAATATCCTGTCTTTC | Sq3 |
| GCCATTTGCAAACGTAGAAAATACCTGGCATG | Sq3 |
| TCATTTGCTAATAGTAGTAGCATT | Sq3 |
| CAACTAAAGTACGGTGGGATGGCT | Sq3 |
| CATTATTAGCAAAAGAAGTTTTGC | Sq3 |
| TTAAGAGGGTCCAATACTGCGGATAGCGAG | Sq3 |
| AGGCTTTTCAGGTAGAAAGATTCAATTACC | Sq3 |
| TTATGCGATTGACAAGAACCGGAGGTCAAT | Sq3 |
| CATAAGGGACACTAAAACACTCACATTAAA | Sq3 |
| CGGGTAAAATTCGGTCGCTGAGGAATGACA | Sq3 |
| TTTCACGTCGATAGTTGCGCCGACCTTGCAGG | Sq3 |
| GAGTTAAATTCATGAGGAAGTTTCTCTTTGAC | Sq3 |
| CCCCAGCGGGAACGAGGCGCAGACTATTCATT | Sq3 |
| ACCCAAATAACTTTAATCATTGTGATCAGTTG | Sq3 |
| AGATTTAGACGATAAAAACCAAAAATCGTCAT | Sq3 |
| AAATATTCCAAAGCGGATTGCATCGAGCTTCA | Sq3 |
| TAGAGCTTCAGACCGGAAGCAAACCTATTATA | Sq3 |
| GTCAGAAGATTGAATCCCCCTCAACCTCGTTT | Sq3 |
| ACCAGACGGAATACCACATTCAACGAGATGGT | Sq3 |
| TTAATTTCCAACGTAACAAAGCTGTCCATGTT | Sq3 |
| ACTTAGCCATTATACCAAGCGCGAGAGGACTA | Sq3 |
| AAGACTTTGGCCGCTTTTGCGGGATTAAACAG | Sq3 |
| CTTGATACTGAAAATCTCCAAAAAAGCGGAGT | Sq3 |
| GTTAGTAACTTTCAACAGTTTCAAAGGCTC | Sq3 |
| CAAAAGGTTCGAGGTGAATTTCTCGTCACC | Sq3 |
| CTCAGCAGGCTACAGAGGCTTTAACAAAGT | Sq3 |
| ACAACGGAAATCCGCGACCTGCCTCATTCA | Sq3 |
| GTGAATATAGTAAATTGGGCTTTAATGCAG | Sq3 |
| ATACATACAACACTATCATAACATGCTTTA | Sq3 |
| AACAGTTAGGTCTTTACCCTGATCCAACAG | Sq3 |
| GTCAGGAAGAGGTCATTTTTGCTCTGGAAG | Sq3 |
| TTTCATTGAGTAGATTTAGTTTCTATATTT | Sq3 |
| CCATGTACCGTAACACTGTAGCATTCCACAGATTCCAGAC | Sq3 |

| | |
|---|---|
| AGTCAGGACATAGGCTGGCTGACCTTTGAAAG | Sq3 |
| GACAACTCGTATTTTTTCCTGTGTGAAATTGTTATCCGAGCTC | Sq3 |
| CCAGGGTGGTTTTGCAAATGAAAAATCTAAA | Sq3 |
| GCCACGCTGTTTTTTTACCAGTGAG | Sq3 |
| ACGGGCAAGTTCCAGTTTTTTCTGACCTGCAACAGT | Sq3 |
| GCCAACATTTTTTCCACTATTAAAGAAATAGGGT | Sq3 |
| CCAACGTCATCGGAACCCTAAAGGGAGCCCTTTTTTTGAACAATATTACCG | Sq3 |
| CAAACTATCGGCCTTGCTGGTTTTTGAGCTTGACGGGG | Sq3 |
| GCGTAACCACCATTTTTGAGTAAAAGAGT | Sq3 |
| CTGTCCATTTTTATAATCATTTTTTTCTTAATGCGCCCACGCTGC | Sq3 |
| AATCATGGTCATTTTTTTTTTTGCCCGAACTCAGGTTTAACTTTTTTTTCAGTATGTTAG | Sq3 |
| ATTAAGACTCCTTTTTAATATACAGTAACAGTACCGAAATTGC | Sq3 |
| CATAAATCAATTTAGTCAGAGGGTAATTGAG | Sq3 |
| AACTGAACATTTTTTTTGAATAACC | Sq3 |
| TTGCTTCTTATATGTATTTTACGCTAACGGAGAATT | Sq3 |
| TTTTATCTTTTTTATCCAATCGCAAGAGTTGGGT | Sq3 |
| GCGAGAAAATAAACACCGGAATCATAATTATTTTTTTCGCCCAATAGCAAG | Sq3 |
| TTTTATTTTCATCGTAGGAATTTTTAGCCTGTTTAGTA | Sq3 |
| AACATGTAATTTTTTTTGAAACCAATCAA | Sq3 |
| TAATCGGCCATCCTAATTTTTTTTTTTTTCGAGCCAACAACGCC | Sq3 |
| AGGACAGATGATTTTTTCACCAGTAGCACCATTACCGACTTGA | Sq3 |
| GAACCGCCTCTTTACCTAAAACGAAAGAGGC | Sq3 |
| TGCCACTACTTTTTTTGCCACCCTC | Sq3 |
| AGAACCGCATTTACCGTTTTACCGATATATACGTAA | Sq3 |
| ACAACCATTTTTTCATACATGGCTTTTAAGCGCA | Sq3 |
| AGGAGTGTAAACATGAAAGTATTAAGAGGCTTTTTTTGCGAATAATAATTT | Sq3 |
| GAGAATAGAAAGGAACAACTATTTTCTCAAGAGAAGGA | Sq3 |
| ACCGTACTCAGGTTTTTGATCTAAAGTTT | Sq3 |
| TGTCGTCTCAGCCCTCATATTTTTTTCGCCACCCTCAGGTGTATC | Sq3 |
| GGAATTAGAGCTTTTTTTTCAGACCAGGCGCGTTGGGAAGATTTTTTTTCCAGGCAAAGC | Sq3 |

| | |
|---|---|
| CCGCTTCTGGTTTTTTCGTTAATAAAACGAACTAAATTATACC | Sq3 |
| CAGAGGGGGTTTTGCCTTCCTGTAGCCAGCT | Sq3 |
| CAAAAATAATTTTTTTTGTTTAGAC | Sq3 |
| TGGATAGCAAGCCCGATTTTTAATCGTAAACGCCAT | Sq3 |
| ACAAGAGTTTTTTTCGCGTTTTAATTCAAAAAGA | Sq3 |
| AAGCGAACAATTGCTGAATATAATGCTGTATTTTTTTGTGAGAAAGGCC GG | Sq3 |
| GAGTAATGTGTAGGTAAAGATTTTTTGTTTTAAATATG | Sq3 |
| GATACATTTCGCTTTTTTGACCCTGTAAT | Sq3 |
| ACTTTTGCATCGGTTGTACTTTTTTTAACCTGTTTAGGACCATTA | Sq3 |
| GTGTCGTAGACACTCCCAATTCTGCGAACCCATATAACAGTTGATGTGT CGTAGACAC | Sq3 |
| GTGTCGTAGACACCCATAAATCAAAAATCCAGAAAACGAGAATGAGTG TCGTAGACAC | Sq3 |
| GTGTCGTAGACACGAAACACCAGAACGAGAGGCTTGCCCTGACGAGTG TCGTAGACAC | Sq3 |
| GTGTCGTAGACACGAACGAGGGTAGCAACGCGAAAGACAGCATCGGTG TCGTAGACAC | Sq3 |
| GTGTCGTAGACACGGGATTTTGCTAAACAAATGAATTTTCTGTATGTGT CGTAGACAC | Sq3 |
| GTGTCGTAGACACGAGAGGGTTGATATAAGCGGATAAGTGCCGTCGTGT CGTAGACAC | Sq3 |
| GTGTCGTAGACACGCAGGTCAGACGATTGTTGACAGGAGGTTGAGGTGT CGTAGACAC | Sq3 |
| GTGTCGTAGACACGCGCCAAAGACAAAAGTTCATATGGTTTACCAGTGT CGTAGACAC | Sq3 |
| GTGTCGTAGACACTTTTTTGTTTAACGTCTCCAAATAAGAAACGAGTGT CGTAGACAC | Sq3 |
| GTGTCGTAGACACTAAACCAAGTACCGCATTCCAAGAACGGGTATGTGT CGTAGACAC | Sq3 |
| GTGTCGTAGACACAGTAGGGCTTAATTGAAAAGCCAACGCTCAACGTGT CGTAGACAC | Sq3 |
| GTGTCGTAGACACAGTCAATAGTGAATTTTTAAGACGCTGAGAAGGTGT CGTAGACAC | Sq3 |
| GTGTCGTAGACACCAATATAATCCTGATTGATGATGGCAATTCATGTGT CGTAGACAC | Sq3 |
| GTGTCGTAGACACACATCGCCATTAAAAAAACTGATAGCCCTAAAGTGT CGTAGACAC | Sq3 |

| | |
|---|---|
| GTGTCGTAGACACTTGATTAGTAATAACATTGTAGCAATACTTCTGTGTCGTAGACAC | Sq3 |
| GTGTCGTAGACACCGGGCGCTAGGGCGCTAAGAAAGCGAAAGGAGGTGTCGTAGACAC | Sq3 |
| GTGTCGTAGACACATCCTGTTTGATGGTGGCCCCAGCAGGCGAAAGTGTCGTAGACAC | Sq3 |
| GTGTCGTAGACACGTAACGCCAGGGTTTTAAGGCGATTAAGTTGGGTGTCGTAGACAC | Sq3 |
| GTGTCGTAGACACTTTAAATTGTAAACGTATTGTATAAGCAAATAGTGTCGTAGACAC | Sq3 |
| GTGTCGTAGACACAAATTTTTAGAACCCTTTCAACGCAAGGATAAGTGTCGTAGACAC | Sq3 |
| CGTACTGACCGTTTCGCTGCAATCTTTTAAGAACTGGCTCCGGAACAA | Sq3, Sq-Apt1 |
| CGTACTGACCGTTTCGCTGCGGCAGAATTATCACCGTCACCATTAGCA | Sq3, Sq-Apt1 |
| CGTACTGACCGTTTCGCTGCTTATGTAAAACAGAAATAAATTTTACAT | Sq3, Sq-Apt1 |
| CGTACTGACCGTTTCGCTGCTTGGAGAGGATCCCCGGGTACCGCTCAC | Sq3, Sq-Apt1 |
| AAAGCCGGCGAACGTGTGCCGTAA | Sq-Apt2 |
| AATTCCACGTTTGCGTATTGGGCG | Sq-Apt2 |
| TTCATCAACGCACTCCAGCCAGCTGCTGCGCA | Sq-Apt2 |
| ACTGTTGGAGAGGATCCCCGGGTACCGCTCAC | Sq-Apt2 |
| TGAGTGTTCAGCTGATTGCCCTTGCGCGGG | Sq-Apt2 |
| AGACAGTCCATTGCCTGAGAGTCTTCATATGT | Sq-Apt2 |
| AATCATACAGGCAAGGCAGAGCATAAAGCTAAGGGAGAAG | Sq-Apt2 |
| GAATTCGTGCCATTCGCCATTCAGTTCCGGCA | Sq-Apt2 |
| TCATATGCGTTATACAAAGGCGTT | Sq-Apt2 |
| CGGGAGAATTTAATGGAAACAGTA | Sq-Apt2 |
| GCATCACCAGTATTAGACTTTACAGTTTGAGT | Sq-Apt2 |
| AACATTATGTAAAACAGAAATAAATTTTACAT | Sq-Apt2 |
| TATATAACGTAAATCGTCGCTATATTTGAA | Sq-Apt2 |
| CCAGCCATCCAGTAATAAAAGGGACGTGGCAC | Sq-Apt2 |
| CTAAACAGGAGGCCGATAATCCTGAGAAGTGTCACGCAAA | Sq-Apt2 |
| GTAGATTTGTTATTAATTTTAAAAAACAATTC | Sq-Apt2 |
| TTAGGATTAGCGGGGTGGAACCTA | Sq-Apt2 |
| AGGCCGGAACCAGAGCCACCACCG | Sq-Apt2 |

| | |
|---|---|
| CGCTAATAGGAATACCCAAAAGAAATACATAA | Sq-Apt2 |
| AGGTGGCAGAATTATCACCGTCACCATTAGCA | Sq-Apt2 |
| GTCTCTGACACCCTCAGAGCCACATCAAAA | Sq-Apt2 |
| CAAATCAGTGCTATTTTGCACCCAGCCTAATT | Sq-Apt2 |
| AGCTAATGCAGAACGCGAGAAAAATAATATCCTGTCTTTC | Sq-Apt2 |
| GCCATTTGCAAACGTAGAAAATACCTGGCATG | Sq-Apt2 |
| CAACTAAAGTACGGTGGGATGGCT | Sq-Apt2 |
| CATTATTAGCAAAAGAAGTTTTGC | Sq-Apt2 |
| AAAAGAATAACCGAACTGACCAACTTCATCAA | Sq-Apt2 |
| GAGTAATCTTTTAAGAACTGGCTCCGGAACAA | Sq-Apt2 |
| TTAAGAGGGTCCAATACTGCGGATAGCGAG | Sq-Apt2 |
| TTTCACGTCGATAGTTGCGCCGACCTTGCAGG | Sq-Apt2 |
| CCATGTACCGTAACACTGTAGCATTCCACAGATTCCAGAC | Sq-Apt2 |
| AGTCAGGACATAGGCTGGCTGACCTTTGAAAG | Sq-Apt2 |
| GACAACTCGTATTTTTTCCTGTGTGAAATTGTTATCCGAGCTC | Sq-Apt2 |
| CCAGGGTGGTTTTGCAAATGAAAAATCTAAA | Sq-Apt2 |
| GCCACGCTGTTTTTTTACCAGTGAG | Sq-Apt2 |
| ACGGGCAAGTTCCAGTTTTTTCTGACCTGCAACAGT | Sq-Apt2 |
| GCCAACATTTTTTCCACTATTAAAGAAATAGGGT | Sq-Apt2 |
| CCAACGTCATCGGAACCCTAAAGGGAGCCCTTTTTTTGAACAATATTAC<br>CG | Sq-Apt2 |
| CAAACTATCGGCCTTGCTGGTTTTTGAGCTTGACGGGG | Sq-Apt2 |
| GCGTAACCACCATTTTTGAGTAAAAGAGT | Sq-Apt2 |
| CTGTCCATTTTTATAATCATTTTTTTCTTAATGCGCCCACGCTGC | Sq-Apt2 |
| AATCATGGTCATTTTTTTTTTTGCCCGAACTCAGGTTTAACTTTTTTTTCA<br>GTATGTTAG | Sq-Apt2 |
| ATTAAGACTCCTTTTTAATATACAGTAACAGTACCGAAATTGC | Sq-Apt2 |
| CATAAATCAATTTAGTCAGAGGGTAATTGAG | Sq-Apt2 |
| AACTGAACATTTTTTTTGAATAACC | Sq-Apt2 |
| TTGCTTCTTATATGTATTTTACGCTAACGGAGAATT | Sq-Apt2 |
| TTTTATCTTTTTTATCCAATCGCAAGAGTTGGGT | Sq-Apt2 |
| GCGAGAAAATAAACACCGGAATCATAATTATTTTTTTCGCCCAATAGCA<br>AG | Sq-Apt2 |
| TTTTATTTTCATCGTAGGAATTTTTAGCCTGTTTAGTA | Sq-Apt2 |

| | |
|---|---|
| AACATGTAATTTTTTTTGAAACCAATCAA | Sq-Apt2 |
| TAATCGGCCATCCTAATTTTTTTTTTTTTTCGAGCCAACAACGCC | Sq-Apt2 |
| AGGACAGATGATTTTTTCACCAGTAGCACCATTACCGACTTGA | Sq-Apt2 |
| GAACCGCCTCTTTACCTAAAACGAAAGAGGC | Sq-Apt2 |
| TGCCACTACTTTTTTTGCCACCCTC | Sq-Apt2 |
| AGAACCGCATTTACCGTTTTACCGATATATACGTAA | Sq-Apt2 |
| ACAACCATTTTTTCATACATGGCTTTTAAGCGCA | Sq-Apt2 |
| AGGAGTGTAAACATGAAAGTATTAAGAGGCTTTTTTTGCGAATAATAAT<br>TT | Sq-Apt2 |
| GAGAATAGAAAGGAACAACTATTTTCTCAAGAGAAGGA | Sq-Apt2 |
| ACCGTACTCAGGTTTTTGATCTAAAGTTT | Sq-Apt2 |
| TGTCGTCTCAGCCCTCATATTTTTTTCGCCACCCTCAGGTGTATC | Sq-Apt2 |
| GGAATTAGAGCTTTTTTTTCAGACCAGGCGCGTTGGGAAGATTTTTTTTC<br>CAGGCAAAGC | Sq-Apt2 |
| CCGCTTCTGGTTTTTTCGTTAATAAAACGAACTAAATTATACC | Sq-Apt2 |
| CAGAGGGGGTTTTGCCTTCCTGTAGCCAGCT | Sq-Apt2 |
| CAAAAATAATTTTTTTTGTTTAGAC | Sq-Apt2 |
| TGGATAGCAAGCCCGATTTTTAATCGTAAACGCCAT | Sq-Apt2 |
| ACAAGAGTTTTTTTCGCGTTTTAATTCAAAAAGA | Sq-Apt2 |
| AAGCGAACAATTGCTGAATATAATGCTGTATTTTTTTGTGAGAAAGGCC<br>GG | Sq-Apt2 |
| GAGTAATGTGTAGGTAAAGATTTTTTGTTTTAAATATG | Sq-Apt2 |
| GATACATTTCGCTTTTTTGACCCTGTAAT | Sq-Apt2 |
| ACTTTTGCATCGGTTGTACTTTTTTTAACCTGTTTAGGACCATTA | Sq-Apt2 |
| GTGTCGTAGACACTCCCAATTCTGCGAACCCATATAACAGTTGATGTGT<br>CGTAGACAC | Sq-Apt2 |
| GTGTCGTAGACACCCATAAATCAAAAATCCAGAAAACGAGAATGAGTG<br>TCGTAGACAC | Sq-Apt2 |
| GTGTCGTAGACACGAAACACCAGAACGAGAGGCTTGCCCTGACGAGTG<br>TCGTAGACAC | Sq-Apt2 |
| GTGTCGTAGACACGAACGAGGGTAGCAACGCGAAAGACAGCATCGGTG<br>TCGTAGACAC | Sq-Apt2 |
| GTGTCGTAGACACGGGATTTTGCTAAACAAATGAATTTTCTGTATGTGT<br>CGTAGACAC | Sq-Apt2 |
| GTGTCGTAGACACGAGAGGGTTGATATAAGCGGATAAGTGCCGTCGTGT<br>CGTAGACAC | Sq-Apt2 |

| | |
|---|---|
| GTGTCGTAGACACGCAGGTCAGACGATTGTTGACAGGAGGTTGAGGTGTCGTAGACAC | Sq-Apt2 |
| GTGTCGTAGACACGCGCCAAAGACAAAAGTTCATATGGTTTACCAGTGTCGTAGACAC | Sq-Apt2 |
| GTGTCGTAGACACTTTTTTGTTTAACGTCTCCAAATAAGAAACGAGTGTCGTAGACAC | Sq-Apt2 |
| GTGTCGTAGACACTAAACCAAGTACCGCATTCCAAGAACGGGTATGTGTCGTAGACAC | Sq-Apt2 |
| GTGTCGTAGACACAGTAGGGCTTAATTGAAAAGCCAACGCTCAACGTGTCGTAGACAC | Sq-Apt2 |
| GTGTCGTAGACACAGTCAATAGTGAATTTTTAAGACGCTGAGAAGGTGTCGTAGACAC | Sq-Apt2 |
| GTGTCGTAGACACCAATATAATCCTGATTGATGATGGCAATTCATGTGTCGTAGACAC | Sq-Apt2 |
| GTGTCGTAGACACACATCGCCATTAAAAAAACTGATAGCCCTAAAGTGTCGTAGACAC | Sq-Apt2 |
| GTGTCGTAGACACTTGATTAGTAATAACATTGTAGCAATACTTCTGTGTCGTAGACAC | Sq-Apt2 |
| GTGTCGTAGACACCGGGCGCTAGGGCGCTAAGAAAGCGAAAGGAGGTGTCGTAGACAC | Sq-Apt2 |
| GTGTCGTAGACACATCCTGTTTGATGGTGGCCCCAGCAGGCGAAAGTGTCGTAGACAC | Sq-Apt2 |
| GTGTCGTAGACACGTAACGCCAGGGTTTTAAGGCGATTAAGTTGGGTGTCGTAGACAC | Sq-Apt2 |
| GTGTCGTAGACACTTTAAATTGTAAACGTATTGTATAAGCAAATAGTGTCGTAGACAC | Sq-Apt2 |
| GTGTCGTAGACACAAATTTTTAGAACCCTTTCAACGCAAGGATAAGTGTCGTAGACAC | Sq-Apt2 |
| GGAGAGAAGAGGGAAGGAAACAGCACCGACCTTGTGCTTTGGGAGTGCTGGTCCAAGGGCGTTAATGGACA | Sq-Apt1, Sq-Apt2 |
| CGTACTGACCGTTTCGCTGCGGTGGCATCAATTCTAGGGCGCGAGCTGAAAA | Sq-Apt2, Sq2 |
| CGTACTGACCGTTTCGCTGCGAAACACCAGAACGAGAGGCTTGCCCTGACGA | Sq-Apt2, Sq2 |
| CGTACTGACCGTTTCGCTGCACAAACTACAACGCCTGAGTTTCGTCACCAGT | Sq-Apt2, Sq2 |
| TTTCCTTCCCTCTTCTCTCCTTTTTTCATTGAGTAGATTTAGTTTCTATATTT | Sq-Apt2 |
| TTTCCTTCCCTCTTCTCTCCTTTGTCAGGAAGAGGTCATTTTTGCTCTGGAAG | Sq-Apt2 |

| | |
|---|---|
| TTTCCTTCCCTCTTCTCTCCTTTAACAGTTAGGTCTTTACCCTGATCCAACAG | Sq-Apt2 |
| TTTCCTTCCCTCTTCTCTCCTTTATACATACAACACTATCATAACATGCTTTA | Sq-Apt2 |
| TTTCCTTCCCTCTTCTCTCCTTTGTGAATATAGTAAATTGGGCTTTAATGCAG | Sq-Apt2 |
| TTTCCTTCCCTCTTCTCTCCTTTACAACGGAAATCCGCGACCTGCCTCATTCA | Sq-Apt2 |
| TTTCCTTCCCTCTTCTCTCCTTTCTCAGCAGGCTACAGAGGCTTTAACAAAGT | Sq-Apt2 |
| TTTCCTTCCCTCTTCTCTCCTTTCAAAAGGTTCGAGGTGAATTTCTCGTCACC | Sq-Apt2 |
| TTTCCTTCCCTCTTCTCTCCTTTGTTAGTAACTTTCAACAGTTTCAAAGGCTC | Sq-Apt2 |
| TTTCCTTCCCTCTTCTCTCCTTTCTTGATACTGAAAATCTCCAAAAAAGCGGAGT | Sq-Apt2 |
| TTTCCTTCCCTCTTCTCTCCTTTAAGACTTTGGCCGCTTTTGCGGGATTAAACAG | Sq-Apt2 |
| TTTCCTTCCCTCTTCTCTCCTTTACTTAGCCATTATACCAAGCGCGAGAGGACTA | Sq-Apt2 |
| TTTCCTTCCCTCTTCTCTCCTTTTTAATTTCCAACGTAACAAAGCTGTCCATGTT | Sq-Apt2 |
| TTTCCTTCCCTCTTCTCTCCTTTACCAGACGGAATACCACATTCAACGAGATGGT | Sq-Apt2 |
| TTTCCTTCCCTCTTCTCTCCTTTGTCAGAAGATTGAATCCCCCTCAACCTCGTTT | Sq-Apt2 |
| TTTCCTTCCCTCTTCTCTCCTTTTAGAGCTTCAGACCGGAAGCAAACCTATTATA | Sq-Apt2 |
| TTTCCTTCCCTCTTCTCTCCTTTAAATATTCCAAAGCGGATTGCATCGAGCTTCA | Sq-Apt2 |
| TTTCCTTCCCTCTTCTCTCCTTTAGATTTAGACGATAAAAACCAAAAATCGTCAT | Sq-Apt2 |
| TTTCCTTCCCTCTTCTCTCCTTTACCCAAATAACTTTAATCATTGTGATCAGTTG | Sq-Apt2 |
| TTTCCTTCCCTCTTCTCTCCTTTCCCCAGCGGGAACGAGGCGCAGACTATTCATT | Sq-Apt2 |
| TTTCCTTCCCTCTTCTCTCCTTTGAGTTAAATTCATGAGGAAGTTTCTCTTTGAC | Sq-Apt2 |
| TTTCCTTCCCTCTTCTCTCCTTTCGGGTAAAATTCGGTCGCTGAGGAATGACA | Sq-Apt2 |

| | |
|---|---|
| TTTCCTTCCCTCTTCTCTCCTTTCATAAGGGACACTAAAACACTCACATT AAA | Sq-Apt2 |
| TTTCCTTCCCTCTTCTCTCCTTTTTATGCGATTGACAAGAACCGGAGGTC AAT | Sq-Apt2 |
| TTTCCTTCCCTCTTCTCTCCTTTAGGCTTTTCAGGTAGAAAGATTCAATTA CC | Sq-Apt2 |
| TTTCCTTCCCTCTTCTCTCCTTTTCATTTGCTAATAGTAGTAGCATT | Sq-Apt2 |
| TTTCCTTCCCTCTTCTCTCCTTTGTACCAGGTATAGCCCGGAATAGAACC GCC | Sq-Apt2 |
| TTTCCTTCCCTCTTCTCTCCTTTCAGTGCCCCCCCTGCCTATTTCTTTGCT CA | Sq-Apt2 |
| TTTCCTTCCCTCTTCTCTCCTTTGCCAGCAGCCTTGATATTCACAAACGG GGT | Sq-Apt2 |
| TTTCCTTCCCTCTTCTCTCCTTTAGTTTGCGCATTTTCGGTCATAGAGCCG CC | Sq-Apt2 |
| TTTCCTTCCCTCTTCTCTCCTTTTAGAAAAGGCGACATTCAACCGCAGAA TCA | Sq-Apt2 |
| TTTCCTTCCCTCTTCTCTCCTTTATGAAATGAAAAGTAAGCAGATACAAT CAA | Sq-Apt2 |
| TTTCCTTCCCTCTTCTCTCCTTTATCCCAAAAAAATGAAAATAGCAAGAA ACA | Sq-Apt2 |
| TTTCCTTCCCTCTTCTCTCCTTTTAAGAACGGAGGTTTTGAAGCCTATTAT TT | Sq-Apt2 |
| TTTCCTTCCCTCTTCTCTCCTTTCTTATCACTCATCGAGAACAAGCGGTAT TC | Sq-Apt2 |
| TTTCCTTCCCTCTTCTCTCCTTTAGATTAGTATATAGAAGGCTTATCCAA GCCGT | Sq-Apt2 |
| TTTCCTTCCCTCTTCTCTCCTTTCAGAGAGAACAAAATAAACAGCCATTA AATCA | Sq-Apt2 |
| TTTCCTTCCCTCTTCTCTCCTTTAAAGTTACGCCCAATAATAAGAGCAGC CTTTA | Sq-Apt2 |
| TTTCCTTCCCTCTTCTCTCCTTTAGGGAAGGATAAGTTTATTTTGTCAGCC GAAC | Sq-Apt2 |
| TTTCCTTCCCTCTTCTCTCCTTTATTAGCGTCCGTAATCAGTAGCGAATTG AGGG | Sq-Apt2 |
| TTTCCTTCCCTCTTCTCTCCTTTAATCCTCAACCAGAACCACCACCAGCC CCCTT | Sq-Apt2 |
| TTTCCTTCCCTCTTCTCTCCTTTTTATTCTGACTGGTAATAAGTTTTAACA AATA | Sq-Apt2 |
| TTTCCTTCCCTCTTCTCTCCTTTGAGCCGCCTTAAAGCCAGAATGGAGAT GATAC | Sq-Apt2 |

| | |
|---|---|
| TTTCCTTCCCTCTTCTCTCCTTTTAGCAGCATTGCCATCTTTTCATACACCCTCA | Sq-Apt2 |
| TTTCCTTCCCTCTTCTCTCCTTTACCACGGATAAATATTGACGGAAAACCATCGA | Sq-Apt2 |
| TTTCCTTCCCTCTTCTCTCCTTTTGAGTTAACAGAAGGAAACCGAGGGCAAAGAC | Sq-Apt2 |
| TTTCCTTCCCTCTTCTCTCCTTTTGCCAGTTATAACATAAAAACAGGACAAGAAT | Sq-Apt2 |
| TTTCCTTCCCTCTTCTCTCCTTTATTAGACGGAGCGTCTTTCCAGAGCTACAA | Sq-Apt2 |
| TTTCCTTCCCTCTTCTCTCCTTTATAATAACTCAGAGAGATAACCCGAAGCGC | Sq-Apt2 |
| TTTCCTTCCCTCTTCTCTCCTTTTTAAAGGTACATATAAAAGAAACAAACGCA | Sq-Apt2 |
| TTTCCTTCCCTCTTCTCTCCTTTTCACCGGAAACGTCACCAATGAATTATTCA | Sq-Apt2 |
| TTTCCTTCCCTCTTCTCTCCTTTACCCTCATTCAGGGATAGCAAGCC | Sq-Apt2 |
| TTTCCTTCCCTCTTCTCTCCTTTCCAGTATGAATCGCCATATTTAGTAATAAG | Sq-Apt2 |
| TTTCCTTCCCTCTTCTCTCCTTTATTTCATGACCGTGTGATAAATAATTCTTA | Sq-Apt2 |
| TTTCCTTCCCTCTTCTCTCCTTTGCTTAGAATCAAAATCATAGGTTTTAGTTA | Sq-Apt2 |
| TTTCCTTCCCTCTTCTCTCCTTTGCGAATTATGAAACAAACATCATAGCGATA | Sq-Apt2 |
| TTTCCTTCCCTCTTCTCTCCTTTATTATCAGTTTGGATTATACTTGCGCAGAG | Sq-Apt2 |
| TTTCCTTCCCTCTTCTCTCCTTTACAGTTGTTAGGAGCACTAACATATTCCTG | Sq-Apt2 |
| TTTCCTTCCCTCTTCTCTCCTTTATGCGCGTACCGAACGAACCACGCAAATCA | Sq-Apt2 |
| TTTCCTTCCCTCTTCTCTCCTTTAATACCTATTTACATTGGCAGAAGTCTTTA | Sq-Apt2 |
| TTTCCTTCCCTCTTCTCTCCTTTTTAACCGTCACTTGCCTGAGTACTCATGGA | Sq-Apt2 |
| TTTCCTTCCCTCTTCTCTCCTTTTCACACGATGCAACAGGAAAAACGGAAGAACT | Sq-Apt2 |
| TTTCCTTCCCTCTTCTCTCCTTTGATAAAACTTTTTGAATGGCTATTTTCACCAG | Sq-Apt2 |
| TTTCCTTCCCTCTTCTCTCCTTTATTAGAGCAATATCTGGTCAGTTGCAGCAGAA | Sq-Apt2 |

| | |
|---|---|
| TTTCCTTCCCTCTTCTCTCCTTTTGGAAGGGAGCGGAATTATCATCAACT AATAG | Sq-Apt2 |
| TTTCCTTCCCTCTTCTCTCCTTTAAATTAATACCAAGTTACAAAATCCTG AATAA | Sq-Apt2 |
| TTTCCTTCCCTCTTCTCTCCTTTCTACCTTTAGAATCCTTGAAAACAAGA AAACA | Sq-Apt2 |
| TTTCCTTCCCTCTTCTCTCCTTTAAATAAGAACTTTTTCAAATATATCTGA GAGA | Sq-Apt2 |
| TTTCCTTCCCTCTTCTCTCCTTTTTTCCCTTTTAACCTCCGGCTTAGCAAA GAAC | Sq-Apt2 |
| TTTCCTTCCCTCTTCTCTCCTTTCTTTGAATTACATTTAACAATTTCTAAT TAAT | Sq-Apt2 |
| TTTCCTTCCCTCTTCTCTCCTTTCCAGAAGGTTAGAACCTACCATATCCT GATTG | Sq-Apt2 |
| TTTCCTTCCCTCTTCTCTCCTTTCCTCAATCCGTCAATAGATAATACAGA AACCA | Sq-Apt2 |
| TTTCCTTCCCTCTTCTCTCCTTTAGACAATAAGAGGTGAGGCGGTCATAT CAAAC | Sq-Apt2 |
| TTTCCTTCCCTCTTCTCTCCTTTCACCGCCTGAAAGCGTAAGAATACATT CTG | Sq-Apt2 |
| TTTCCTTCCCTCTTCTCTCCTTTGATTTAGATTGCTGAACCTCAAAGTATT AA | Sq-Apt2 |
| TTTCCTTCCCTCTTCTCTCCTTTATTTGCACCATTTTGCGGAACAAATTTG AG | Sq-Apt2 |
| TTTCCTTCCCTCTTCTCTCCTTTTTACCTTTACAATAACGGATTCGCAAAA TT | Sq-Apt2 |
| TTTCCTTCCCTCTTCTCTCCTTTAGAATATCAGACGACGACAATAAA | Sq-Apt2 |
| TTTCCTTCCCTCTTCTCTCCTTTGGAAGGGGGCAAGTGTAGCGGTGCTAC AGG | Sq-Apt2 |
| TTTCCTTCCCTCTTCTCTCCTTTGGCGATGTTTTTGGGGTCGAGGGCGAG AAA | Sq-Apt2 |
| TTTCCTTCCCTCTTCTCTCCTTTCTGGTTTGTTCCGAAATCGGCATCTATC AG | Sq-Apt2 |
| TTTCCTTCCCTCTTCTCTCCTTTCTAACTCCCAGTCGGGAAACCTGGTCC ACG | Sq-Apt2 |
| TTTCCTTCCCTCTTCTCTCCTTTGTGCTGCCCCAGTCACGACGTTTGAGTG AG | Sq-Apt2 |
| TTTCCTTCCCTCTTCTCTCCTTTATTGACCCGCATCGTAACCGTGAGGGG GAT | Sq-Apt2 |
| TTTCCTTCCCTCTTCTCTCCTTTCAGGAAGTAATATTTTGTTAAAAACGG CGG | Sq-Apt2 |

| | |
|---|---|
| TTTCCTTCCCTCTTCTCTCCTTTACCGTTCATTTTTGAGAGATCTCCCAAAAA | Sq-Apt2 |
| TTTCCTTCCCTCTTCTCTCCTTTCCTTTATCATATATTTTAAATGGATATTCA | Sq-Apt2 |
| TTTCCTTCCCTCTTCTCTCCTTTTATCAGGTAAATCACCATCAATATCAATGCCT | Sq-Apt2 |
| TTTCCTTCCCTCTTCTCTCCTTTTAAATTTTTGATAATCAGAAAAGCACAAAGGC | Sq-Apt2 |
| TTTCCTTCCCTCTTCTCTCCTTTAGTTTGAGATTCTCCGTGGGAACAATTCGCAT | Sq-Apt2 |
| TTTCCTTCCCTCTTCTCTCCTTTACGGCCAGTACGCCAGCTGGCGAACATCTGCC | Sq-Apt2 |
| TTTCCTTCCCTCTTCTCTCCTTTAGCTGCATAGCCTGGGGTGCCTAAGTAAAACG | Sq-Apt2 |
| TTTCCTTCCCTCTTCTCTCCTTTTATAAATCGAGAGTTGCAGCAAGCGTCGTGCC | Sq-Apt2 |
| TTTCCTTCCCTCTTCTCTCCTTTAGCACTAAAAAGGGCGAAAAACCGAAATCCCT | Sq-Apt2 |
| TTTCCTTCCCTCTTCTCTCCTTTGGCCCTGAAAAAGAATAGCCCGAGCGTGGACT | Sq-Apt2 |
| TTTCCTTCCCTCTTCTCTCCTTTAAGTGTAATAATGAATCGGCCAACCACCGCCT | Sq-Apt2 |
| TTTCCTTCCCTCTTCTCTCCTTTTTCGCTATTGCCAAGCTTGCATGCGAAGCATA | Sq-Apt2 |
| TTTCCTTCCCTCTTCTCTCCTTTCCCGTCGGGGGACGACGACAGTATCGGGCCTC | Sq-Apt2 |
| TTTCCTTCCCTCTTCTCTCCTTTACCCCGGTTGTTAAATCAGCTCATAGTAACAA | Sq-Apt2 |
| TTTCCTTCCCTCTTCTCTCCTTTCCAATAGGAAACTAGCATGTCAAGGAGCAA | Sq-Apt2 |
| TTTCCTTCCCTCTTCTCTCCTTTAGGAAGATCATTAAATGTGAGCGTTTTTAA | Sq-Apt2 |
| TTTCCTTCCCTCTTCTCTCCTTTTCGACTCTGAAGGGCGATCGGTGCGGCCTC | Sq-Apt2 |
| TTTCCTTCCCTCTTCTCTCCTTTGAGAGGCGACAACATACGAGCCGCTGCAGG | Sq-Apt2 |
| TTTCCTTCCCTCTTCTCTCCTTTGCGCGTACTTTCCTCGTTAGAATC | Sq-Apt2 |
| AAAAGAATAACCGAACTGACCAACTTCATCAAGAGT | Sq3, Sq-Apt1 |
| CGCTAATAGGAATACCCAAAAGAAATACATAAAGGT | Sq3, Sq-Apt1 |

| | |
|---|---|
| GCATCACCAGTATTAGACTTTACAGTTTGAGTAACA | Sq3, Sq-Apt1 |
| TTCATCAACGCACTCCAGCCAGCTGCTGCGCAACTG | Sq3, Sq-Apt1 |
| AAAGCCGGCGAACGTGTGCCGTAA | Sq-Apt1 |
| AATTCCACGTTTGCGTATTGGGCG | Sq-Apt1 |
| TGAGTGTTCAGCTGATTGCCCTTGCGCGGG | Sq-Apt1 |
| AGACAGTCCATTGCCTGAGAGTCTTCATATGT | Sq-Apt1 |
| AATCATACAGGCAAGGCAGAGCATAAAGCTAAGGGAGAAG | Sq-Apt1 |
| GAATTCGTGCCATTCGCCATTCAGTTCCGGCA | Sq-Apt1 |
| TCATATGCGTTATACAAAGGCGTT | Sq-Apt1 |
| CGGGAGAATTTAATGGAAACAGTA | Sq-Apt1 |
| TATATAACGTAAATCGTCGCTATATTTGAA | Sq-Apt1 |
| CCAGCCATCCAGTAATAAAAGGGACGTGGCAC | Sq-Apt1 |
| CTAAACAGGAGGCCGATAATCCTGAGAAGTGTCACGCAAA | Sq-Apt1 |
| GTAGATTTGTTATTAATTTTAAAAAACAATTC | Sq-Apt1 |
| TTAGGATTAGCGGGGTGGAACCTA | Sq-Apt1 |
| AGGCCGGAACCAGAGCCACCACCG | Sq-Apt1 |
| GTCTCTGACACCCTCAGAGCCACATCAAAA | Sq-Apt1 |
| CAAATCAGTGCTATTTTGCACCCAGCCTAATT | Sq-Apt1 |
| AGCTAATGCAGAACGCGAGAAAAATAATATCCTGTCTTTC | Sq-Apt1 |
| GCCATTTGCAAACGTAGAAAATACCTGGCATG | Sq-Apt1 |
| CAACTAAAGTACGGTGGGATGGCT | Sq-Apt1 |
| CATTATTAGCAAAAGAAGTTTTGC | Sq-Apt1 |
| TTAAGAGGGTCCAATACTGCGGATAGCGAG | Sq-Apt1 |
| TTTCACGTCGATAGTTGCGCCGACCTTGCAGG | Sq-Apt1 |
| CCATGTACCGTAACACTGTAGCATTCCACAGATTCCAGAC | Sq-Apt1 |
| AGTCAGGACATAGGCTGGCTGACCTTTGAAAG | Sq-Apt1 |
| GACAACTCGTATTTTTCCTGTGTGAAATTGTTATCCGAGCTC | Sq-Apt1 |
| CCAGGGTGGTTTTGCAAATGAAAAATCTAAA | Sq-Apt1 |
| GCCACGCTGTTTTTTTACCAGTGAG | Sq-Apt1 |
| ACGGGCAAGTTCCAGTTTTTCTGACCTGCAACAGT | Sq-Apt1 |
| GCCAACATTTTTTCCACTATTAAAGAAATAGGGT | Sq-Apt1 |

| | |
|---|---|
| CCAACGTCATCGGAACCCTAAAGGGAGCCCTTTTTTTGAACAATATTACCG | Sq-Apt1 |
| CAAACTATCGGCCTTGCTGGTTTTTGAGCTTGACGGGG | Sq-Apt1 |
| GCGTAACCACCATTTTTGAGTAAAAGAGT | Sq-Apt1 |
| CTGTCCATTTTTATAATCATTTTTTTCTTAATGCGCCCACGCTGC | Sq-Apt1 |
| AATCATGGTCATTTTTTTTTTTGCCCGAACTCAGGTTTAACTTTTTTTTCAGTATGTTAG | Sq-Apt1 |
| ATTAAGACTCCTTTTTAATATACAGTAACAGTACCGAAATTGC | Sq-Apt1 |
| CATAAATCAATTTAGTCAGAGGGTAATTGAG | Sq-Apt1 |
| AACTGAACATTTTTTTTGAATAACC | Sq-Apt1 |
| TTGCTTCTTATATGTATTTTACGCTAACGGAGAATT | Sq-Apt1 |
| TTTTATCTTTTTTATCCAATCGCAAGAGTTGGGT | Sq-Apt1 |
| GCGAGAAAATAAACACCGGAATCATAATTATTTTTTTCGCCCAATAGCAAG | Sq-Apt1 |
| TTTTATTTTCATCGTAGGAATTTTTAGCCTGTTTAGTA | Sq-Apt1 |
| AACATGTAATTTTTTTTGAAACCAATCAA | Sq-Apt1 |
| TAATCGGCCATCCTAATTTTTTTTTTTTTCGAGCCAACAACGCC | Sq-Apt1 |
| AGGACAGATGATTTTTTCACCAGTAGCACCATTACCGACTTGA | Sq-Apt1 |
| GAACCGCCTCTTTACCTAAAACGAAAGAGGC | Sq-Apt1 |
| TGCCACTACTTTTTTTGCCACCCTC | Sq-Apt1 |
| AGAACCGCATTTACCGTTTTACCGATATATACGTAA | Sq-Apt1 |
| ACAACCATTTTTTCATACATGGCTTTTAAGCGCA | Sq-Apt1 |
| AGGAGTGTAAACATGAAAGTATTAAGAGGCTTTTTTTGCGAATAATAATTT | Sq-Apt1 |
| GAGAATAGAAAGGAACAACTATTTTCTCAAGAGAAGGA | Sq-Apt1 |
| ACCGTACTCAGGTTTTTGATCTAAAGTTT | Sq-Apt1 |
| TGTCGTCTCAGCCCTCATATTTTTTTCGCCACCCTCAGGTGTATC | Sq-Apt1 |
| GGAATTAGAGCTTTTTTTTCAGACCAGGCGCGTTGGGAAGATTTTTTTTCCAGGCAAAGC | Sq-Apt1 |
| CCGCTTCTGGTTTTTTCGTTAATAAAACGAACTAAATTATACC | Sq-Apt1 |
| CAGAGGGGGTTTTGCCTTCCTGTAGCCAGCT | Sq-Apt1 |
| CAAAAATAATTTTTTTTGTTTAGAC | Sq-Apt1 |
| TGGATAGCAAGCCCGATTTTTAATCGTAAACGCCAT | Sq-Apt1 |
| ACAAGAGTTTTTTTCGCGTTTTAATTCAAAAAGA | Sq-Apt1 |

| | |
|---|---|
| AAGCGAACAATTGCTGAATATAATGCTGTATTTTTTTGTGAGAAAGGCC GG | Sq-Apt1 |
| GAGTAATGTGTAGGTAAAGATTTTTTGTTTTAAATATG | Sq-Apt1 |
| GATACATTTCGCTTTTTTGACCCTGTAAT | Sq-Apt1 |
| ACTTTTGCATCGGTTGTACTTTTTTTAACCTGTTTAGGACCATTA | Sq-Apt1 |
| GTGTCGTAGACACTCCCAATTCTGCGAACCCATATAACAGTTGATGTGT CGTAGACAC | Sq-Apt1 |
| GTGTCGTAGACACCCATAAATCAAAAATCCAGAAAACGAGAATGAGTG TCGTAGACAC | Sq-Apt1 |
| GTGTCGTAGACACGAAACACCAGAACGAGAGGCTTGCCCTGACGAGTG TCGTAGACAC | Sq-Apt1 |
| GTGTCGTAGACACGAACGAGGGTAGCAACGCGAAAGACAGCATCGGTG TCGTAGACAC | Sq-Apt1 |
| GTGTCGTAGACACGGGATTTTGCTAAACAAATGAATTTTCTGTATGTGT CGTAGACAC | Sq-Apt1 |
| GTGTCGTAGACACGAGAGGGTTGATATAAGCGGATAAGTGCCGTCGTGT CGTAGACAC | Sq-Apt1 |
| GTGTCGTAGACACGCAGGTCAGACGATTGTTGACAGGAGGTTGAGGTGT CGTAGACAC | Sq-Apt1 |
| GTGTCGTAGACACGCGCCAAAGACAAAAGTTCATATGGTTTACCAGTGT CGTAGACAC | Sq-Apt1 |
| GTGTCGTAGACACTTTTTTGTTTAACGTCTCCAAATAAGAAACGAGTGT CGTAGACAC | Sq-Apt1 |
| GTGTCGTAGACACTAAACCAAGTACCGCATTCCAAGAACGGGTATGTGT CGTAGACAC | Sq-Apt1 |
| GTGTCGTAGACACAGTAGGGCTTAATTGAAAAGCCAACGCTCAACGTGT CGTAGACAC | Sq-Apt1 |
| GTGTCGTAGACACAGTCAATAGTGAATTTTTAAGACGCTGAGAAGGTGT CGTAGACAC | Sq-Apt1 |
| GTGTCGTAGACACCAATATAATCCTGATTGATGATGGCAATTCATGTGT CGTAGACAC | Sq-Apt1 |
| GTGTCGTAGACACACATCGCCATTAAAAAAACTGATAGCCCTAAAGTGT CGTAGACAC | Sq-Apt1 |
| GTGTCGTAGACACTTGATTAGTAATAACATTGTAGCAATACTTCTGTGTC GTAGACAC | Sq-Apt1 |
| GTGTCGTAGACACCGGGCGCTAGGGCGCTAAGAAAGCGAAAGGAGGTG TCGTAGACAC | Sq-Apt1 |
| GTGTCGTAGACACATCCTGTTTGATGGTGGCCCCAGCAGGCGAAAGTGT CGTAGACAC | Sq-Apt1 |

| | |
|---|---|
| GTGTCGTAGACACGTAACGCCAGGGTTTTAAGGCGATTAAGTTGGGTGTCGTAGACAC | Sq-Apt1 |
| GTGTCGTAGACACTTTAAATTGTAAACGTATTGTATAAGCAAATAGTGTCGTAGACAC | Sq-Apt1 |
| GTGTCGTAGACACAAATTTTTAGAACCCTTTCAACGCAAGGATAAGTGTCGTAGACAC | Sq-Apt1 |
| TTTCCTTCCCTCTTCTCTCCTTTTTTCATTGAGTAGATTTAGTTTCTATATTT | Sq-Apt1 |
| TTTCCTTCCCTCTTCTCTCCTTTGTCAGGAAGAGGTCATTTTTGCTCTGGAAG | Sq-Apt1 |
| TTTCCTTCCCTCTTCTCTCCTTTAACAGTTAGGTCTTTACCCTGATCCAACAG | Sq-Apt1 |
| TTTCCTTCCCTCTTCTCTCCTTTATACATACAACACTATCATAACATGCTTTA | Sq-Apt1 |
| TTTCCTTCCCTCTTCTCTCCTTTGTGAATATAGTAAATTGGGCTTTAATGCAG | Sq-Apt1 |
| TTTCCTTCCCTCTTCTCTCCTTTACAACGGAAATCCGCGACCTGCCTCATTCA | Sq-Apt1 |
| TTTCCTTCCCTCTTCTCTCCTTTCTCAGCAGGCTACAGAGGCTTTAACAAAGT | Sq-Apt1 |
| TTTCCTTCCCTCTTCTCTCCTTTCAAAAGGTTCGAGGTGAATTTCTCGTCACC | Sq-Apt1 |
| TTTCCTTCCCTCTTCTCTCCTTTGTTAGTAACTTTCAACAGTTTCAAAGGCTC | Sq-Apt1 |
| TTTCCTTCCCTCTTCTCTCCTTTCTTGATACTGAAAATCTCCAAAAAAGCGGAGT | Sq-Apt1 |
| TTTCCTTCCCTCTTCTCTCCTTTAAGACTTTGGCCGCTTTTGCGGGATTAAACAG | Sq-Apt1 |
| TTTCCTTCCCTCTTCTCTCCTTTACTTAGCCATTATACCAAGCGCGAGAGGACTA | Sq-Apt1 |
| TTTCCTTCCCTCTTCTCTCCTTTTTAATTTCCAACGTAACAAAGCTGTCCATGTT | Sq-Apt1 |
| TTTCCTTCCCTCTTCTCTCCTTTACCAGACGGAATACCACATTCAACGAGATGGT | Sq-Apt1 |
| TTTCCTTCCCTCTTCTCTCCTTTGTCAGAAGATTGAATCCCCCTCAACCTCGTTT | Sq-Apt1 |
| TTTCCTTCCCTCTTCTCTCCTTTTAGAGCTTCAGACCGGAAGCAAACCTATTATA | Sq-Apt1 |
| TTTCCTTCCCTCTTCTCTCCTTTAAATATTCCAAAGCGGATTGCATCGAGCTTCA | Sq-Apt1 |

| | |
|---|---|
| TTTCCTTCCCTCTTCTCTCCTTTAGATTTAGACGATAAAAACCAAAAATC GTCAT | Sq-Apt1 |
| TTTCCTTCCCTCTTCTCTCCTTTACCCAAATAACTTTAATCATTGTGATCA GTTG | Sq-Apt1 |
| TTTCCTTCCCTCTTCTCTCCTTTCCCCAGCGGGAACGAGGCGCAGACTAT TCATT | Sq-Apt1 |
| TTTCCTTCCCTCTTCTCTCCTTTGAGTTAAATTCATGAGGAAGTTTCTCTT TGAC | Sq-Apt1 |
| TTTCCTTCCCTCTTCTCTCCTTTCGGGTAAAATTCGGTCGCTGAGGAATG ACA | Sq-Apt1 |
| TTTCCTTCCCTCTTCTCTCCTTTCATAAGGGACACTAAAACACTCACATT AAA | Sq-Apt1 |
| TTTCCTTCCCTCTTCTCTCCTTTTTATGCGATTGACAAGAACCGGAGGTC AAT | Sq-Apt1 |
| TTTCCTTCCCTCTTCTCTCCTTTAGGCTTTTCAGGTAGAAAGATTCAATTA CC | Sq-Apt1 |
| TTTCCTTCCCTCTTCTCTCCTTTAAAAGAATAACCGAACTGACCAACTTC ATCAA | Sq-Apt1 |
| TTTCCTTCCCTCTTCTCTCCTTTTCATTTGCTAATAGTAGTAGCATT | Sq-Apt1 |
| TTTCCTTCCCTCTTCTCTCCTTTGTACCAGGTATAGCCCGGAATAGAACC GCC | Sq-Apt1 |
| TTTCCTTCCCTCTTCTCTCCTTTCAGTGCCCCCCCTGCCTATTTCTTTGCT CA | Sq-Apt1 |
| TTTCCTTCCCTCTTCTCTCCTTTGCCAGCAGCCTTGATATTCACAAACGG GGT | Sq-Apt1 |
| TTTCCTTCCCTCTTCTCTCCTTTAGTTTGCGCATTTTCGGTCATAGAGCCG CC | Sq-Apt1 |
| TTTCCTTCCCTCTTCTCTCCTTTTAGAAAAGGCGACATTCAACCGCAGAA TCA | Sq-Apt1 |
| TTTCCTTCCCTCTTCTCTCCTTTATGAAATGAAAAGTAAGCAGATACAAT CAA | Sq-Apt1 |
| TTTCCTTCCCTCTTCTCTCCTTTATCCCAAAAAAATGAAAATAGCAAGAA ACA | Sq-Apt1 |
| TTTCCTTCCCTCTTCTCTCCTTTTAAGAACGGAGGTTTTGAAGCCTATTAT TT | Sq-Apt1 |
| TTTCCTTCCCTCTTCTCTCCTTTCTTATCACTCATCGAGAACAAGCGGTAT TC | Sq-Apt1 |
| TTTCCTTCCCTCTTCTCTCCTTTAGATTAGTATATAGAAGGCTTATCCAA GCCGT | Sq-Apt1 |
| TTTCCTTCCCTCTTCTCTCCTTTCAGAGAGAACAAAATAAACAGCCATTA AATCA | Sq-Apt1 |

| | |
|---|---|
| TTTCCTTCCCTCTTCTCTCCTTTAAAGTTACGCCCAATAATAAGAGCAGCCTTTA | Sq-Apt1 |
| TTTCCTTCCCTCTTCTCTCCTTTAGGGAAGGATAAGTTTATTTTGTCAGCCGAAC | Sq-Apt1 |
| TTTCCTTCCCTCTTCTCTCCTTTATTAGCGTCCGTAATCAGTAGCGAATTGAGGG | Sq-Apt1 |
| TTTCCTTCCCTCTTCTCTCCTTTAATCCTCAACCAGAACCACCACCAGCCCCCTT | Sq-Apt1 |
| TTTCCTTCCCTCTTCTCTCCTTTTTATTCTGACTGGTAATAAGTTTTAACAAATA | Sq-Apt1 |
| TTTCCTTCCCTCTTCTCTCCTTTGAGCCGCCTTAAAGCCAGAATGGAGATGATAC | Sq-Apt1 |
| TTTCCTTCCCTCTTCTCTCCTTTTAGCAGCATTGCCATCTTTTCATACACCCTCA | Sq-Apt1 |
| TTTCCTTCCCTCTTCTCTCCTTTACCACGGATAAATATTGACGGAAAACCATCGA | Sq-Apt1 |
| TTTCCTTCCCTCTTCTCTCCTTTTGAGTTAACAGAAGGAAACCGAGGGCAAAGAC | Sq-Apt1 |
| TTTCCTTCCCTCTTCTCTCCTTTTGCCAGTTATAACATAAAAACAGGACAAGAAT | Sq-Apt1 |
| TTTCCTTCCCTCTTCTCTCCTTTATTAGACGGAGCGTCTTTCCAGAGCTACAA | Sq-Apt1 |
| TTTCCTTCCCTCTTCTCTCCTTTATAATAACTCAGAGAGATAACCCGAAGCGC | Sq-Apt1 |
| TTTCCTTCCCTCTTCTCTCCTTTTTAAAGGTACATATAAAAGAAACAAACGCA | Sq-Apt1 |
| TTTCCTTCCCTCTTCTCTCCTTTTCACCGGAAACGTCACCAATGAATTATTCA | Sq-Apt1 |
| TTTCCTTCCCTCTTCTCTCCTTTCGCTAATAGGAATACCCAAAAGAAATACATAA | Sq-Apt1 |
| TTTCCTTCCCTCTTCTCTCCTTTACCCTCATTCAGGGATAGCAAGCC | Sq-Apt1 |
| TTTCCTTCCCTCTTCTCTCCTTTCCAGTATGAATCGCCATATTTAGTAATAAG | Sq-Apt1 |
| TTTCCTTCCCTCTTCTCTCCTTTATTTCATGACCGTGTGATAAATAATTCTTA | Sq-Apt1 |
| TTTCCTTCCCTCTTCTCTCCTTTGCTTAGAATCAAAATCATAGGTTTTAGTTA | Sq-Apt1 |
| TTTCCTTCCCTCTTCTCTCCTTTGCGAATTATGAAACAAACATCATAGCGATA | Sq-Apt1 |
| TTTCCTTCCCTCTTCTCTCCTTTATTATCAGTTTGGATTATACTTGCGCAGAG | Sq-Apt1 |

| | |
|---|---|
| TTTCCTTCCCTCTTCTCTCCTTTACAGTTGTTAGGAGCACTAACATATTCCTG | Sq-Apt1 |
| TTTCCTTCCCTCTTCTCTCCTTTATGCGCGTACCGAACGAACCACGCAAATCA | Sq-Apt1 |
| TTTCCTTCCCTCTTCTCTCCTTTAATACCTATTTACATTGGCAGAAGTCTTTA | Sq-Apt1 |
| TTTCCTTCCCTCTTCTCTCCTTTTTAACCGTCACTTGCCTGAGTACTCATGGA | Sq-Apt1 |
| TTTCCTTCCCTCTTCTCTCCTTTTCACACGATGCAACAGGAAAAACGGAAGAACT | Sq-Apt1 |
| TTTCCTTCCCTCTTCTCTCCTTTGATAAAACTTTTTGAATGGCTATTTTCACCAG | Sq-Apt1 |
| TTTCCTTCCCTCTTCTCTCCTTTATTAGAGCAATATCTGGTCAGTTGCAGCAGAA | Sq-Apt1 |
| TTTCCTTCCCTCTTCTCTCCTTTTGGAAGGGAGCGGAATTATCATCAACTAATAG | Sq-Apt1 |
| TTTCCTTCCCTCTTCTCTCCTTTAAATTAATACCAAGTTACAAAATCCTGAATAA | Sq-Apt1 |
| TTTCCTTCCCTCTTCTCTCCTTTCTACCTTTAGAATCCTTGAAAACAAGAAAACA | Sq-Apt1 |
| TTTCCTTCCCTCTTCTCTCCTTTAAATAAGAACTTTTTCAAATATATCTGAGAGA | Sq-Apt1 |
| TTTCCTTCCCTCTTCTCTCCTTTTTTCCCTTTTAACCTCCGGCTTAGCAAAGAAC | Sq-Apt1 |
| TTTCCTTCCCTCTTCTCTCCTTTCTTTGAATTACATTTAACAATTTCTAATTAAT | Sq-Apt1 |
| TTTCCTTCCCTCTTCTCTCCTTTCCAGAAGGTTAGAACCTACCATATCCTGATTG | Sq-Apt1 |
| TTTCCTTCCCTCTTCTCTCCTTTCCTCAATCCGTCAATAGATAATACAGAAACCA | Sq-Apt1 |
| TTTCCTTCCCTCTTCTCTCCTTTAGACAATAAGAGGTGAGGCGGTCATATCAAAC | Sq-Apt1 |
| TTTCCTTCCCTCTTCTCTCCTTTCACCGCCTGAAAGCGTAAGAATACATTCTG | Sq-Apt1 |
| TTTCCTTCCCTCTTCTCTCCTTTGATTTAGATTGCTGAACCTCAAAGTATTAA | Sq-Apt1 |
| TTTCCTTCCCTCTTCTCTCCTTTATTTGCACCATTTTGCGGAACAAATTTGAG | Sq-Apt1 |
| TTTCCTTCCCTCTTCTCTCCTTTTTACCTTTACAATAACGGATTCGCAAAATT | Sq-Apt1 |

| | |
|---|---|
| TTTCCTTCCCTCTTCTCTCCTTTGCATCACCAGTATTAGACTTTACAGTTTGAGT | Sq-Apt1 |
| TTTCCTTCCCTCTTCTCTCCTTTAGAATATCAGACGACGACAATAAA | Sq-Apt1 |
| TTTCCTTCCCTCTTCTCTCCTTTGGAAGGGGGCAAGTGTAGCGGTGCTACAGG | Sq-Apt1 |
| TTTCCTTCCCTCTTCTCTCCTTTGGCGATGTTTTTGGGGTCGAGGGCGAGAAA | Sq-Apt1 |
| TTTCCTTCCCTCTTCTCTCCTTTCTGGTTTGTTCCGAAATCGGCATCTATCAG | Sq-Apt1 |
| TTTCCTTCCCTCTTCTCTCCTTTCTAACTCCCAGTCGGGAAACCTGGTCCACG | Sq-Apt1 |
| TTTCCTTCCCTCTTCTCTCCTTTGTGCTGCCCCAGTCACGACGTTTGAGTGAG | Sq-Apt1 |
| TTTCCTTCCCTCTTCTCTCCTTTATTGACCCGCATCGTAACCGTGAGGGGGAT | Sq-Apt1 |
| TTTCCTTCCCTCTTCTCTCCTTTCAGGAAGTAATATTTTGTTAAAAACGGCGG | Sq-Apt1 |
| TTTCCTTCCCTCTTCTCTCCTTTACCGTTCATTTTTGAGAGATCTCCCAAAAA | Sq-Apt1 |
| TTTCCTTCCCTCTTCTCTCCTTTCCTTTATCATATATTTTAAATGGATATTCA | Sq-Apt1 |
| TTTCCTTCCCTCTTCTCTCCTTTTATCAGGTAAATCACCATCAATATCAATGCCT | Sq-Apt1 |
| TTTCCTTCCCTCTTCTCTCCTTTTAAATTTTTGATAATCAGAAAAGCACAAAGGC | Sq-Apt1 |
| TTTCCTTCCCTCTTCTCTCCTTTAGTTTGAGATTCTCCGTGGGAACAATTCGCAT | Sq-Apt1 |
| TTTCCTTCCCTCTTCTCTCCTTTACGGCCAGTACGCCAGCTGGCGAACATCTGCC | Sq-Apt1 |
| TTTCCTTCCCTCTTCTCTCCTTTAGCTGCATAGCCTGGGGTGCCTAAGTAAAACG | Sq-Apt1 |
| TTTCCTTCCCTCTTCTCTCCTTTTATAAATCGAGAGTTGCAGCAAGCGTCGTGCC | Sq-Apt1 |
| TTTCCTTCCCTCTTCTCTCCTTTAGCACTAAAAAGGGCGAAAAACCGAAATCCCT | Sq-Apt1 |
| TTTCCTTCCCTCTTCTCTCCTTTGGCCCTGAAAAAGAATAGCCCGAGCGTGGACT | Sq-Apt1 |
| TTTCCTTCCCTCTTCTCTCCTTTAAGTGTAATAATGAATCGGCCAACCACCGCCT | Sq-Apt1 |
| TTTCCTTCCCTCTTCTCTCCTTTTTCGCTATTGCCAAGCTTGCATGCGAAGCATA | Sq-Apt1 |

| | |
|---|---|
| TTTCCTTCCCTCTTCTCTCCTTTCCCGTCGGGGGACGACGACAGTATCGGGCCTC | Sq-Apt1 |
| TTTCCTTCCCTCTTCTCTCCTTTACCCCGGTTGTTAAATCAGCTCATAGTAACAA | Sq-Apt1 |
| TTTCCTTCCCTCTTCTCTCCTTTCCAATAGGAAACTAGCATGTCAAGGAGCAA | Sq-Apt1 |
| TTTCCTTCCCTCTTCTCTCCTTTAGGAAGATCATTAAATGTGAGCGTTTTTAA | Sq-Apt1 |
| TTTCCTTCCCTCTTCTCTCCTTTTCGACTCTGAAGGGCGATCGGTGCGGCCTC | Sq-Apt1 |
| TTTCCTTCCCTCTTCTCTCCTTTGAGAGGCGACAACATACGAGCCGCTGCAGG | Sq-Apt1 |
| TTTCCTTCCCTCTTCTCTCCTTTTTCATCAACGCACTCCAGCCAGCTGCTGCGCA | Sq-Apt1 |
| TTTCCTTCCCTCTTCTCTCCTTTGCGCGTACTTTCCTCGTTAGAATC | Sq-Apt1 |
| GCGCGTACTTTCCTCGTTAGAATC | Sq2 |
| AAAGCCGGCGAACGTGTGCCGTAA | Sq2 |
| AATTCCACGTTTGCGTATTGGGCG | Sq2 |
| TTCATCAACGCACTCCAGCCAGCTGCTGCGCA | Sq2 |
| ACTGTTGGAGAGGATCCCCGGGTACCGCTCAC | Sq2 |
| TGAGTGTTCAGCTGATTGCCCTTGCGCGGG | Sq2 |
| GAGAGGCGACAACATACGAGCCGCTGCAGG | Sq2 |
| TCGACTCTGAAGGGCGATCGGTGCGGCCTC | Sq2 |
| AGGAAGATCATTAAATGTGAGCGTTTTTAA | Sq2 |
| CCAATAGGAAACTAGCATGTCAAGGAGCAA | Sq2 |
| AGACAGTCCATTGCCTGAGAGTCTTCATATGT | Sq2 |
| ACCCCGGTTGTTAAATCAGCTCATAGTAACAA | Sq2 |
| CCCGTCGGGGGACGACGACAGTATCGGGCCTC | Sq2 |
| TTCGCTATTGCCAAGCTTGCATGCGAAGCATA | Sq2 |
| AAGTGTAATAATGAATCGGCCAACCACCGCCT | Sq2 |
| GGCCCTGAAAAAGAATAGCCCGAGCGTGGACT | Sq2 |
| AGCACTAAAAAGGGCGAAAAACCGAAATCCCT | Sq2 |
| TATAAATCGAGAGTTGCAGCAAGCGTCGTGCC | Sq2 |
| AGCTGCATAGCCTGGGGTGCCTAAGTAAAACG | Sq2 |
| ACGGCCAGTACGCCAGCTGGCGAACATCTGCC | Sq2 |

| | |
|---|---|
| AGTTTGAGATTCTCCGTGGGAACAATTCGCAT | Sq2 |
| TAAATTTTTGATAATCAGAAAAGCACAAAGGC | Sq2 |
| TATCAGGTAAATCACCATCAATATCAATGCCT | Sq2 |
| CCTTTATCATATATTTTAAATGGATATTCA | Sq2 |
| ACCGTTCATTTTTGAGAGATCTCCCAAAAA | Sq2 |
| CAGGAAGTAATATTTTGTTAAAAACGGCGG | Sq2 |
| ATTGACCCGCATCGTAACCGTGAGGGGGAT | Sq2 |
| GTGCTGCCCCAGTCACGACGTTTGAGTGAG | Sq2 |
| CTAACTCCCAGTCGGGAAACCTGGTCCACG | Sq2 |
| CTGGTTTGTTCCGAAATCGGCATCTATCAG | Sq2 |
| GGCGATGTTTTTGGGGTCGAGGGCGAGAAA | Sq2 |
| GGAAGGGGGCAAGTGTAGCGGTGCTACAGG | Sq2 |
| AATCATACAGGCAAGGCAGAGCATAAAGCTAAGGGAGAAG | Sq2 |
| GAATTCGTGCCATTCGCCATTCAGTTCCGGCA | Sq2 |
| AGAATATCAGACGACGACAATAAA | Sq2 |
| TCATATGCGTTATACAAAGGCGTT | Sq2 |
| CGGGAGAATTTAATGGAAACAGTA | Sq2 |
| GCATCACCAGTATTAGACTTTACAGTTTGAGT | Sq2 |
| AACATTATGTAAAACAGAAATAAATTTTACAT | Sq2 |
| TATATAACGTAAATCGTCGCTATATTTGAA | Sq2 |
| TTACCTTTACAATAACGGATTCGCAAAATT | Sq2 |
| ATTTGCACCATTTTGCGGAACAAATTTGAG | Sq2 |
| GATTTAGATTGCTGAACCTCAAAGTATTAA | Sq2 |
| CACCGCCTGAAAGCGTAAGAATACATTCTG | Sq2 |
| CCAGCCATCCAGTAATAAAAGGGACGTGGCAC | Sq2 |
| AGACAATAAGAGGTGAGGCGGTCATATCAAAC | Sq2 |
| CCTCAATCCGTCAATAGATAATACAGAAACCA | Sq2 |
| CCAGAAGGTTAGAACCTACCATATCCTGATTG | Sq2 |
| CTTTGAATTACATTTAACAATTTCTAATTAAT | Sq2 |
| TTTCCCTTTTAACCTCCGGCTTAGCAAAGAAC | Sq2 |
| AAATAAGAACTTTTTCAAATATATCTGAGAGA | Sq2 |
| CTACCTTTAGAATCCTTGAAAACAAGAAAACA | Sq2 |
| AAATTAATACCAAGTTACAAAATCCTGAATAA | Sq2 |

| | |
|---|---|
| TGGAAGGGAGCGGAATTATCATCAACTAATAG | Sq2 |
| ATTAGAGCAATATCTGGTCAGTTGCAGCAGAA | Sq2 |
| GATAAAACTTTTTGAATGGCTATTTTCACCAG | Sq2 |
| TCACACGATGCAACAGGAAAAACGGAAGAACT | Sq2 |
| TTAACCGTCACTTGCCTGAGTACTCATGGA | Sq2 |
| AATACCTATTTACATTGGCAGAAGTCTTTA | Sq2 |
| ATGCGCGTACCGAACGAACCACGCAAATCA | Sq2 |
| ACAGTTGTTAGGAGCACTAACATATTCCTG | Sq2 |
| ATTATCAGTTTGGATTATACTTGCGCAGAG | Sq2 |
| GCGAATTATGAAACAAACATCATAGCGATA | Sq2 |
| GCTTAGAATCAAAATCATAGGTTTTAGTTA | Sq2 |
| ATTTCATGACCGTGTGATAAATAATTCTTA | Sq2 |
| CCAGTATGAATCGCCATATTTAGTAATAAG | Sq2 |
| CTAAACAGGAGGCCGATAATCCTGAGAAGTGTCACGCAAA | Sq2 |
| GTAGATTTGTTATTAATTTTAAAAAACAATTC | Sq2 |
| ACCCTCATTCAGGGATAGCAAGCC | Sq2 |
| TTAGGATTAGCGGGGTGGAACCTA | Sq2 |
| AGGCCGGAACCAGAGCCACCACCG | Sq2 |
| CGCTAATAGGAATACCCAAAAGAAATACATAA | Sq2 |
| AGGTGGCAGAATTATCACCGTCACCATTAGCA | Sq2 |
| GTCTCTGACACCCTCAGAGCCACATCAAAA | Sq2 |
| TCACCGGAAACGTCACCAATGAATTATTCA | Sq2 |
| TTAAAGGTACATATAAAAGAAACAAACGCA | Sq2 |
| ATAATAACTCAGAGAGATAACCCGAAGCGC | Sq2 |
| ATTAGACGGAGCGTCTTTCCAGAGCTACAA | Sq2 |
| CAAATCAGTGCTATTTTGCACCCAGCCTAATT | Sq2 |
| TGCCAGTTATAACATAAAAACAGGACAAGAAT | Sq2 |
| TGAGTTAACAGAAGGAAACCGAGGGCAAAGAC | Sq2 |
| ACCACGGATAAATATTGACGGAAAACCATCGA | Sq2 |
| TAGCAGCATTGCCATCTTTTCATACACCCTCA | Sq2 |
| GAGCCGCCTTAAAGCCAGAATGGAGATGATAC | Sq2 |
| TTATTCTGACTGGTAATAAGTTTTAACAAATA | Sq2 |
| AATCCTCAACCAGAACCACCACCAGCCCCCTT | Sq2 |

| | |
|---|---|
| ATTAGCGTCCGTAATCAGTAGCGAATTGAGGG | Sq2 |
| AGGGAAGGATAAGTTTATTTTGTCAGCCGAAC | Sq2 |
| AAAGTTACGCCCAATAATAAGAGCAGCCTTTA | Sq2 |
| CAGAGAGAACAAAATAAACAGCCATTAAATCA | Sq2 |
| AGATTAGTATATAGAAGGCTTATCCAAGCCGT | Sq2 |
| CTTATCACTCATCGAGAACAAGCGGTATTC | Sq2 |
| TAAGAACGGAGGTTTTGAAGCCTATTATTT | Sq2 |
| ATCCCAAAAAAATGAAAATAGCAAGAAACA | Sq2 |
| ATGAAATGAAAAGTAAGCAGATACAATCAA | Sq2 |
| TAGAAAAGGCGACATTCAACCGCAGAATCA | Sq2 |
| AGTTTGCGCATTTTCGGTCATAGAGCCGCC | Sq2 |
| GCCAGCAGCCTTGATATTCACAAACGGGGT | Sq2 |
| CAGTGCCCCCCCTGCCTATTTCTTTGCTCA | Sq2 |
| GTACCAGGTATAGCCCGGAATAGAACCGCC | Sq2 |
| AGCTAATGCAGAACGCGAGAAAAATAATATCCTGTCTTTC | Sq2 |
| GCCATTTGCAAACGTAGAAAATACCTGGCATG | Sq2 |
| TCATTTGCTAATAGTAGTAGCATT | Sq2 |
| CAACTAAAGTACGGTGGGATGGCT | Sq2 |
| CATTATTAGCAAAAGAAGTTTTGC | Sq2 |
| AAAAGAATAACCGAACTGACCAACTTCATCAA | Sq2 |
| GAGTAATCTTTTAAGAACTGGCTCCGGAACAA | Sq2 |
| TTAAGAGGGTCCAATACTGCGGATAGCGAG | Sq2 |
| AGGCTTTTCAGGTAGAAAGATTCAATTACC | Sq2 |
| TTATGCGATTGACAAGAACCGGAGGTCAAT | Sq2 |
| CATAAGGGACACTAAAACACTCACATTAAA | Sq2 |
| CGGGTAAAATTCGGTCGCTGAGGAATGACA | Sq2 |
| TTTCACGTCGATAGTTGCGCCGACCTTGCAGG | Sq2 |
| GAGTTAAATTCATGAGGAAGTTTCTCTTTGAC | Sq2 |
| CCCCAGCGGGAACGAGGCGCAGACTATTCATT | Sq2 |
| ACCCAAATAACTTTAATCATTGTGATCAGTTG | Sq2 |
| AGATTTAGACGATAAAAACCAAAAATCGTCAT | Sq2 |
| AAATATTCCAAAGCGGATTGCATCGAGCTTCA | Sq2 |
| TAGAGCTTCAGACCGGAAGCAAACCTATTATA | Sq2 |

| | |
|---|---|
| GTCAGAAGATTGAATCCCCCTCAACCTCGTTT | Sq2 |
| ACCAGACGGAATACCACATTCAACGAGATGGT | Sq2 |
| TTAATTTCCAACGTAACAAAGCTGTCCATGTT | Sq2 |
| ACTTAGCCATTATACCAAGCGCGAGAGGACTA | Sq2 |
| AAGACTTTGGCCGCTTTTGCGGGATTAAACAG | Sq2 |
| CTTGATACTGAAAATCTCCAAAAAAGCGGAGT | Sq2 |
| GTTAGTAACTTTCAACAGTTTCAAAGGCTC | Sq2 |
| CAAAAGGTTCGAGGTGAATTTCTCGTCACC | Sq2 |
| CTCAGCAGGCTACAGAGGCTTTAACAAAGT | Sq2 |
| ACAACGGAAATCCGCGACCTGCCTCATTCA | Sq2 |
| GTGAATATAGTAAATTGGGCTTTAATGCAG | Sq2 |
| ATACATACAACACTATCATAACATGCTTTA | Sq2 |
| AACAGTTAGGTCTTTACCCTGATCCAACAG | Sq2 |
| GTCAGGAAGAGGTCATTTTTGCTCTGGAAG | Sq2 |
| TTTCATTGAGTAGATTTAGTTTCTATATTT | Sq2 |
| CCATGTACCGTAACACTGTAGCATTCCACAGATTCCAGAC | Sq2 |
| AGTCAGGACATAGGCTGGCTGACCTTTGAAAG | Sq2 |
| GACAACTCGTATTTTTTCCTGTGTGAAATTGTTATCCGAGCTC | Sq2 |
| CCAGGGTGGTTTTGCAAATGAAAAATCTAAA | Sq2 |
| GCCACGCTGTTTTTTTACCAGTGAG | Sq2 |
| ACGGGCAAGTTCCAGTTTTTTCTGACCTGCAACAGT | Sq2 |
| GCCAACATTTTTTCCACTATTAAAGAAATAGGGT | Sq2 |
| CCAACGTCATCGGAACCCTAAAGGGAGCCCTTTTTTTGAACAATATTACCG | Sq2 |
| CAAACTATCGGCCTTGCTGGTTTTTGAGCTTGACGGGG | Sq2 |
| GCGTAACCACCATTTTTGAGTAAAAGAGT | Sq2 |
| CTGTCCATTTTTATAATCATTTTTTTCTTAATGCGCCCACGCTGC | Sq2 |
| AATCATGGTCATTTTTTTTTTTGCCCGAACTCAGGTTTAACTTTTTTTCAGTATGTTAG | Sq2 |
| ATTAAGACTCCTTTTTAATATACAGTAACAGTACCGAAATTGC | Sq2 |
| CATAAATCAATTTAGTCAGAGGGTAATTGAG | Sq2 |
| AACTGAACATTTTTTTTGAATAACC | Sq2 |
| TTGCTTCTTATATGTATTTTACGCTAACGGAGAATT | Sq2 |

| | |
|---|---|
| TTTTATCTTTTTTATCCAATCGCAAGAGTTGGGT | Sq2 |
| GCGAGAAAATAAACACCGGAATCATAATTATTTTTTTCGCCCAATAGCAAG | Sq2 |
| TTTTATTTTCATCGTAGGAATTTTTAGCCTGTTTAGTA | Sq2 |
| AACATGTAATTTTTTTTGAAACCAATCAA | Sq2 |
| TAATCGGCCATCCTAATTTTTTTTTTTTTTCGAGCCAACAACGCC | Sq2 |
| AGGACAGATGATTTTTTCACCAGTAGCACCATTACCGACTTGA | Sq2 |
| GAACCGCCTCTTTACCTAAAACGAAAGAGGC | Sq2 |
| TGCCACTACTTTTTTTGCCACCCTC | Sq2 |
| AGAACCGCATTTACCGTTTTACCGATATATACGTAA | Sq2 |
| ACAACCATTTTTTCATACATGGCTTTTAAGCGCA | Sq2 |
| AGGAGTGTAAACATGAAAGTATTAAGAGGCTTTTTTTGCGAATAATAATTT | Sq2 |
| GAGAATAGAAAGGAACAACTATTTTCTCAAGAGAAGGA | Sq2 |
| ACCGTACTCAGGTTTTTGATCTAAAGTTT | Sq2 |
| TGTCGTCTCAGCCCTCATATTTTTTTCGCCACCCTCAGGTGTATC | Sq2 |
| GGAATTAGAGCTTTTTTTTCAGACCAGGCGCGTTGGGAAGATTTTTTTCCAGGCAAAGC | Sq2 |
| CCGCTTCTGGTTTTTTCGTTAATAAAACGAACTAAATTATACC | Sq2 |
| CAGAGGGGGTTTTGCCTTCCTGTAGCCAGCT | Sq2 |
| CAAAAATAATTTTTTTTGTTTAGAC | Sq2 |
| TGGATAGCAAGCCCGATTTTTAATCGTAAACGCCAT | Sq2 |
| ACAAGAGTTTTTTTCGCGTTTTAATTCAAAAAGA | Sq2 |
| AAGCGAACAATTGCTGAATATAATGCTGTATTTTTTTGTGAGAAAGGCCGG | Sq2 |
| GAGTAATGTGTAGGTAAAGATTTTTTGTTTTAAATATG | Sq2 |
| GATACATTTCGCTTTTTTGACCCTGTAAT | Sq2 |
| ACTTTTGCATCGGTTGTACTTTTTTTAACCTGTTTAGGACCATTA | Sq2 |
| GTGTCGTAGACACTCCCAATTCTGCGAACCCATATAACAGTTGATGTGTCGTAGACAC | Sq2 |
| GTGTCGTAGACACCCATAAATCAAAAATCCAGAAAACGAGAATGAGTGTCGTAGACAC | Sq2 |
| GTGTCGTAGACACGAAACACCAGAACGAGAGGCTTGCCCTGACGAGTGTCGTAGACAC | Sq2 |
| GTGTCGTAGACACGAACGAGGGTAGCAACGCGAAAGACAGCATCGGTGTCGTAGACAC | Sq2 |

| | |
|---|---|
| GTGTCGTAGACACGGGATTTTGCTAAACAAATGAATTTTCTGTATGTGT CGTAGACAC | Sq2 |
| GTGTCGTAGACACGAGAGGGTTGATATAAGCGGATAAGTGCCGTCGTGT CGTAGACAC | Sq2 |
| GTGTCGTAGACACGCAGGTCAGACGATTGTTGACAGGAGGTTGAGGTGT CGTAGACAC | Sq2 |
| GTGTCGTAGACACGCGCCAAAGACAAAAGTTCATATGGTTTACCAGTGT CGTAGACAC | Sq2 |
| GTGTCGTAGACACTTTTTTGTTTAACGTCTCCAAATAAGAAACGAGTGT CGTAGACAC | Sq2 |
| GTGTCGTAGACACTAAACCAAGTACCGCATTCCAAGAACGGGTATGTGT CGTAGACAC | Sq2 |
| GTGTCGTAGACACAGTAGGGCTTAATTGAAAAGCCAACGCTCAACGTGT CGTAGACAC | Sq2 |
| GTGTCGTAGACACAGTCAATAGTGAATTTTTAAGACGCTGAGAAGGTGT CGTAGACAC | Sq2 |
| GTGTCGTAGACACCAATATAATCCTGATTGATGATGGCAATTCATGTGT CGTAGACAC | Sq2 |
| GTGTCGTAGACACACATCGCCATTAAAAAAACTGATAGCCCTAAAGTGT CGTAGACAC | Sq2 |
| GTGTCGTAGACACTTGATTAGTAATAACATTGTAGCAATACTTCTGTGTC GTAGACAC | Sq2 |
| GTGTCGTAGACACCGGGCGCTAGGGCGCTAAGAAAGCGAAAGGAGGTG TCGTAGACAC | Sq2 |
| GTGTCGTAGACACATCCTGTTTGATGGTGGCCCCAGCAGGCGAAAGTGT CGTAGACAC | Sq2 |
| GTGTCGTAGACACGTAACGCCAGGGTTTTAAGGCGATTAAGTTGGGTGT CGTAGACAC | Sq2 |
| GTGTCGTAGACACTTTAAATTGTAAACGTATTGTATAAGCAAATAGTGT CGTAGACAC | Sq2 |
| GTGTCGTAGACACAAATTTTTAGAACCCTTTCAACGCAAGGATAAGTGT CGTAGACAC | Sq2 |
